# Engineering Tumor Stroma Morphogenesis Using Dynamic Cell-Matrix Spheroid Assembly

**DOI:** 10.1101/2024.03.19.585805

**Authors:** Michael J. Buckenmeyer, Elizabeth A. Brooks, Madison S. Taylor, Liping Yang, Ronald J. Holewinski, Thomas J. Meyer, Mélissa Galloux, Marcial Garmendia-Cedillos, Thomas J. Pohida, Thorkell Andresson, Brad St. Croix, Matthew T. Wolf

## Abstract

The tumor microenvironment consists of resident tumor cells organized within a compositionally diverse, three-dimensional (3D) extracellular matrix (ECM) network that cannot be replicated in vitro using bottom-up synthesis. We report a new self-assembly system to engineer ECM-rich 3D MatriSpheres wherein tumor cells actively organize and concentrate microgram quantities of decellularized ECM dispersions which modulate cell phenotype. 3D colorectal cancer (CRC) MatriSpheres were created using decellularized small intestine submucosa (SIS) as an orthotopic ECM source that had greater proteomic homology to CRC tumor ECM than traditional ECM formulations such as Matrigel. SIS ECM was rapidly concentrated from its environment and assembled into ECM-rich 3D stroma-like regions by mouse and human CRC cell lines within 4-5 days via a mechanism that was rheologically distinct from bulk hydrogel formation. Both ECM organization and transcriptional regulation by 3D ECM cues affected programs of malignancy, lipid metabolism, and immunoregulation that corresponded with an in vivo MC38 tumor cell subpopulation identified via single cell RNA sequencing. This 3D modeling approach stimulates tumor specific tissue morphogenesis that incorporates the complexities of both cancer cell and ECM compartments in a scalable, spontaneous assembly process that may further facilitate precision medicine.

## Introduction

Three-dimensional (3D) engineered tumor models have emerged as essential tools to replicate the complexities of the tumor microenvironment (TME). The TME is a multifaceted and dynamic niche composed not only of cancer cells, but diverse stromal cell lineages^1^. These cells predominantly manufacture and maintain a 3D tumor-specific extracellular matrix (ECM), which affects colorectal cancer (CRC) phenotype, metastatic invasiveness, and sensitivity to therapy^2–4^. The ECM is composed of a diverse network of proteins and polysaccharides that have many functions. Namely, the ECM provides structural support and can directly influence cell signaling, which is mediated by cell-ECM ligand interactions^5^. In the context of CRC, the ECM undergoes substantial remodeling^6^, leading to alterations in its composition, organization, and mechanical properties. These alterations stimulate CRC tumor survival, proliferation, migration, and invasion, and can contribute to the development of chemoresistance^7^. Thus, the ECM is a critical feature of the TME and is required to accurately mimic in vivo phenotypes that are predictive for drug discovery and precision medicine. Despite its importance, engineering an ECM-rich tumor stroma that reflects the compositional diversity of native tissues in vitro remains a challenge.

Recent technological advancements have ushered in the development of sophisticated 3D models, such as spheroids^8^ and organoids^9,10^, which offer a more accurate representation of the TME, although each presents limitations^11^. Spheroids offer a high-throughput and reproducible method for efficiently screening 3D cancer cell behavior; however, the long-time scale of ECM deposition and contributions from both stromal and cancer cell types are barriers to ECM development within traditional tumor spheroids^12^. Alternatively, the most frequently applied paradigm for 3D ECM organoid culture is to embed tumor cells within hydrogels such as Matrigel and purified Type I Collagen. These models have yielded substantial insights yet are limited in several ways: (1) ECM composition is difficult to modulate and does not reflect the complexity of native tissues, and (2) hydrogel polymerization does not recapitulate the stromal organization that is characteristic of CRC and other tumors^13^. Decellularized tissues are a form of ECM biomaterials that are frequently used as scaffolds to impart tissue-specific ECM composition and complexity. Optimized decellularization techniques remove cells from mammalian tissues to isolate compositionally intact ECM that is reproducible and scalable for commercial applications^14^. For instance, ECM formulations such as scaffolds, particles and hydrogels have been used for tissue reconstruction in vivo and as a substrate to mimic ECM microenvironments in vitro^12,15,16^. Therefore, decellularized tissues could be useful tools to augment tumor models, which to date have primarily involved passive ECM assembly rather than cell-driven tumor morphogenesis that occurs in vivo.

In this study, we developed tumor MatriSpheres: a high-throughput method of in vitro tumor morphogenesis using orthotopic intestine ECM and CRC cells to recapitulate the TME. We found that unlike previous methods, ECM assembly is cell-mediated and cell line specific. Furthermore, MatriSpheres recapitulate CRC transcriptional phenotypes found in vivo that are not found in traditional spheroids alone. We describe the parameters that facilitate cell-mediated ECM organization and concentration in a process that is distinct from bulk hydrogel formation, enables modular and diverse ECM composition, and is compatible with other purified ECM biomaterials such as Matrigel and Type I Collagen. This approach for consistently generating ECM-enhanced 3D tumor spheroids could be used to further understand the complex interplay between cancer cells and ECM and lead to the development of more effective therapies using precision medicine^17^.

## Results

### SIS ECM is an orthotopic ECM biomaterial for modeling the CRC TME

The ECM of tissues is highly complex; it is encoded by hundreds of genes, is shaped by dozens of post-translational modifications including incompletely characterized polysaccharides such as glycosaminoglycans (GAGs), and matrix bound vesicles^18^. We therefore explored biologically derived materials that preserve this natural complexity as a facile and reproducible source for harnessing ECM diversity that cannot be recreated via bottom-up synthesis approaches. Our initial aim was to generate and proteomically characterize an ECM biomaterial that consists of the most abundant, naturally occurring ECM components found within the CRC TME. To accomplish this, we decellularized porcine small intestine submucosa as a reproducible orthotopic tissue ECM source that has previously been established as a biocompatible scaffold for tissue engineering applications (**Fig. 1a**)^19^. Histologic staining demonstrated that cells were removed from decellularized tissues while retaining a dense collagen-rich microarchitecture. Partial enzymatic digestion of lyophilized SIS ECM resulted in a liquid ECM dispersion that could be used as a supplement for 3D cultures. Transmission electron micrographs (TEM) of SIS ECM digest highlighted loosely associated, fibrous ECM proteins along with nanosized structures consistent with matrix-bound extracellular vesicles within the ECM dispersion. Further characterization indicated double stranded DNA reduction from 4124±902.4 ng/mg in native tissue to 214.9±20.56 ng/mg in decellularized SIS ECM material and substantial DNA fragmentation with no bands detected > 350 bp in decellularized samples (**Fig. 1b,c**).

**Figure 1:**
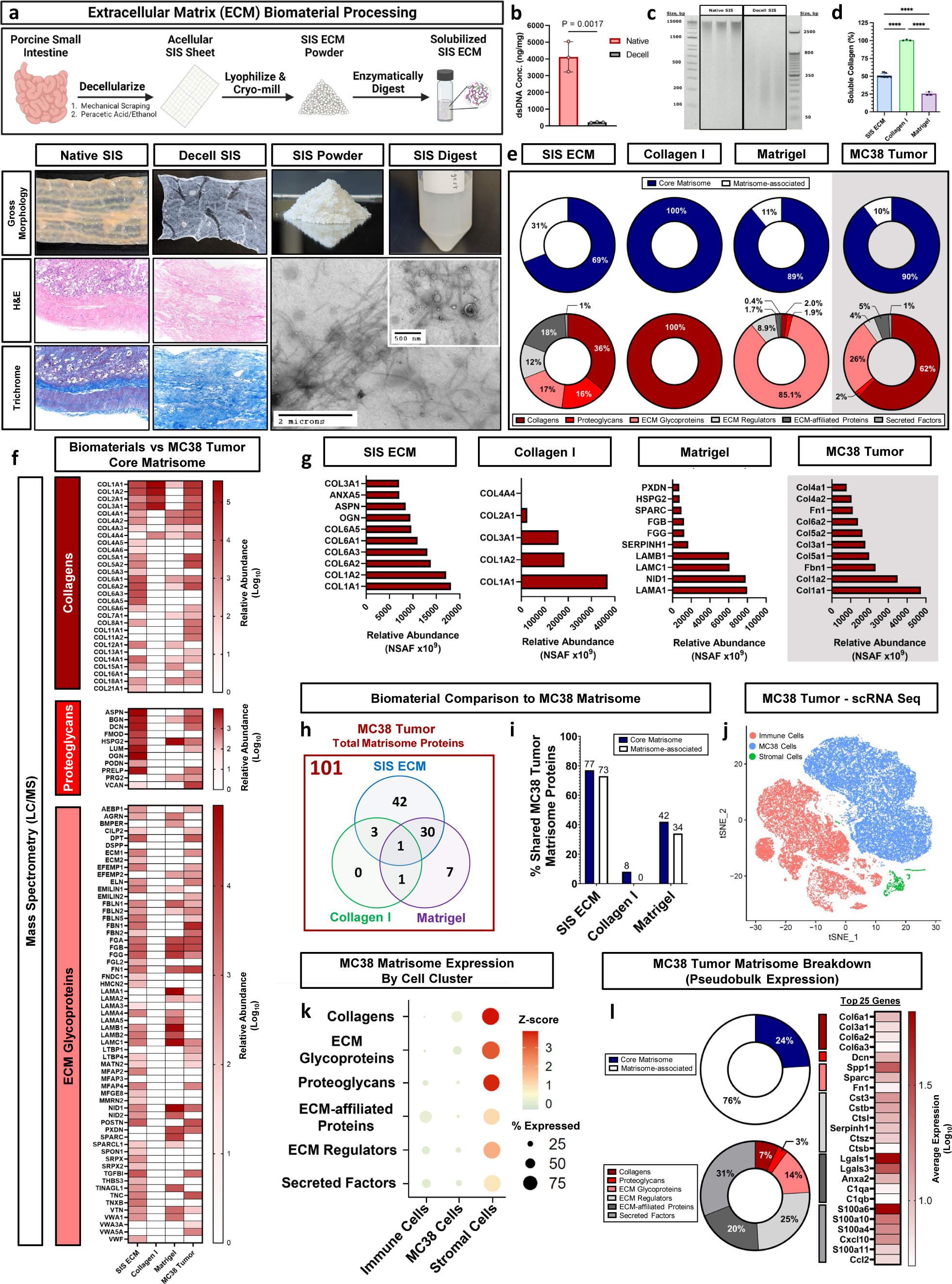
SIS ECM matrisome composition reflects CRC tumor ECM diversity. a. SIS ECM processing workflow and characterization. Histological staining with H&E and Trichrome shows the removal of cells and retention of dense collagen. Transmission electron m Figure 1: SIS ECM matrisome composition reflects CRC tumor ECM diversity a. SIS ECM processing workflow and characterization. Histological staining with H&E and Trichrome shows the removal of cells and retention of dense collagen. Transmission electron micrographs (TEM) of digested SIS ECM highlight the presence of intact fibrils and extracellular vesicles. b. PicoGreen assay indicates a significant reduction in dsDNA content and c. DNA gel electrophoresis demonstrates a substantial decrease in DNA fragment size post-decellularization. d. Sircol assay displays variance in soluble collagen content of ECM biomaterials. e. Matrisome category breakdown and f. Core matrisome composition between ECM biomaterials compared with MC38 tumor ECM based upon relative abundance, detected by mass spectrometry (LC-MS). g. Top 10 matrisome proteins by relative abundance. h. Total shared number and i. percentage of matrisome proteins between ECM biomaterials and MC38 tumor. j. tSNE plot of in vivo cell populations within MC38 tumors (Immune, MC38, and Stromal cells) based upon clustering of scRNA Seq expression data. k. Matrisome expression based upon cell type within MC38 tumors. l. Pseudobulk expression data of MC38 tumor matrisome and top 25 matrisome genes expressed. Plotted data are the mean ± SD. Statistics were calculated by a one-way ANOVA followed by a Tukey’s multiple comparisons test. Statistical significance where p < 0.0001 is denoted with ****.

We compared SIS ECM composition to other naturally derived biomaterials that are frequently utilized to mimic the ECM for in vitro tumor modeling. These include Matrigel, which is a standard tool for 3D tumor organoid generation, and purified Type I Collagen (Collagen I), that is one of the most abundant ECM types in solid tumors and a popular ECM biomaterial for augmenting cell culture applications. Although these biomaterials represent critical components of the ECM, we hypothesized that they provide only a fraction of the broad ECM diversity found within colorectal tumors. One obvious difference in ECM composition were the ratios of total soluble non-crosslinked fibrillar Collagen content. We found that SIS ECM is composed of 50.9% ± 2.8% soluble collagen by mass compared to Matrigel which is 25.5% ± 2.3% (**Fig. 1d**). These results highlighted general compositional differences in Collagen between the ECM biomaterials.

We used mass spectrometry to comprehensively characterize and estimate relative proportions of the remaining proteins within SIS ECM compared to the other ECM biomaterials using preparation methods that emphasize identification sensitivity and coverage. Protein categorization was defined by the matrisome, a list of 1,062 known gene-encoded ECM proteins^20^. SIS ECM was the most compositionally diverse material, with 145 matrisome proteins detected compared to 69 in Matrigel, while Collagen I was the least varied with 5 proteins. SIS ECM diversity was in part due to greater representation of matrisome-associated constituents in addition to structural core matrisome proteins (**Fig. 1e**). We then sought to quantify homology between these ECM biomaterials and in vivo CRC tumor ECM composition. We isolated CRC tumor ECM from syngeneic MC38 mouse tumors since they have a complete TME with intact immune compartment, unlike tumor xenografts. We identified 101 total matrisome proteins within MC38 tumor ECM, which was composed primarily of Collagens (62%) and ECM Glycoproteins (26%). Core matrisome analysis showed that SIS ECM contained the greatest diversity of all ECM biomaterials with Collagens (23), Proteoglycans (10) and ECM Glycoproteins (47) (**Fig. 1f**). The most abundant matrisome proteins in SIS ECM were fibrillar collagens *(Col1a1, Col1a2, Col6a1, Col6a2 and Col6a3)*, proteoglycans *(Ogn and Aspn) and* ECM-affiliated protein *(Anxa5)*, which mediates extracellular vesicle binding^21^ (**Fig. 1g**). We confirmed that commercially available Collagen I biomaterials were predominantly composed of Collagen I chains *Col1a1* and *Col1a2*, but also substantial Type III Collagen (*Col3a1)*, while Matrigel was enriched with the basement membrane ECM Glycoprotein laminin-111 (*Lama1, Nid1, Lamb1, and Lamc1*). We compared homology between ECM biomaterials and in vivo MC38 tumor ECM and found that out of the 101 matrisome proteins identified within MC38 tumor ECM, 42 were exclusively found in SIS ECM suggesting that it provides tissue-specific constituents that are absent in traditional ECM biomaterials (**Fig. 1h**). In contrast, Matrigel provided 7 exclusive proteins. In summary, SIS ECM contained the highest percentage of shared MC38 tumor ECM matrisome proteins (∼75%), which provides a robust and diverse ECM biomaterial for modeling the CRC tumor stroma in vitro (**Fig. 1i**).

The ECM is the sum contribution of all resident cells within a tissue, and a probable advantage of a decellularized tissue ECM is capturing these contributions that would otherwise be missing from a 3D culture system. We hypothesized that proteins represented in decellularized SIS ECM were the result of expression by multiple cell types and were not solely expressed in 3D tumor monoculture. We defined three primary cell compartments from the in vivo MC38 TME using single cell RNA sequencing (scRNA Seq): (1) MC38 cancer cells, (2) immune cells and (3) stromal cells (**Fig. 1j**). Each of these cell compartments expressed unique ECM gene signatures in the production or maintenance of tumor ECM (**Fig. 1k**). We found that stromal cells (i.e., fibroblasts, endothelial cells, pericytes) represented the lowest proportion of total cells but account for the highest percentage of total matrisome gene expression compared to substantially lower expression from more abundant MC38 cancer cells and immune cells. This suggests that in vitro cancer cell monocultures may lack robust ECM production and could indicate that supplementation with ECM biomaterials may be required to effectively model tumor cell-ECM interactions. Examining the scRNA Seq data more closely, MC38 tumor cell matrisome expression predominantly showed a dependence on matrisome-associated genes with a less prevalent expression of core matrisome genes (**Fig. 1l**). ECM Regulators (31%) and Secreted Factors (25%) represented the genes with the highest level of matrisome expression. This shift toward matrisome-associated genes could indicate that mRNA expression from established tumors is in response to pre-existing ECM rather than robust de novo structural ECM production.

We then assessed several colorectal cancer cell lines (MC38, CT26 and HT-29) for core matrisome expression using bulk RNA sequencing to determine how ECM expression profiles differ in traditional 3D spheroid monocultures in vitro **(Supplementary Fig. 1a-b)**. Examining the core matrisome genes, we found shared expression of 28 Collagen isoforms, 12 Proteoglycans, and 101 ECM Glycoproteins across all three cell lines **(Supplementary Fig. 1c)**. We identified a disparity in proteomic conservation of ECM biomaterials to predicted core matrisome expression. Specifically, we found that SIS ECM was the most similar (∼50%) to average CRC spheroid matrisome genes, whereas Matrigel showed greater than 2-fold reduction in overlapping matrisome proteins (∼22%), followed by Collagen I (∼4%), which had the least similar ECM profile **(Supplementary Fig. 1d)**. This suggests that decellularized SIS ECM biomaterials have greater similarity and capacity to replicate to the striking compositional diversity of ECM proteins produced in tumors and supports its use as an orthotopic scaffold for modeling the CRC TME. Additionally, 124 core matrisome genes (18 Collagens, 81 ECM Glycoproteins and 25 Proteoglycans) expressed within in vivo MC38 tumors were not found in vitro **(Supplementary Fig. 1e)**. The top five matrisome genes exclusively found in in vivo MC38 tumors were Srgn, Mfap5, Fgl2, Col15a1 and Spon1 **(Supplementary Fig. 1f)**. Each of these genes were predominantly expressed by non-cancer cells (immune cells and/or fibroblasts) within the MC38 TME. These data support that tumor ECM production and modifiers arise from diverse cell populations within the TME and cannot be deposited by cancer cells alone in vitro.

### Decellularized ECM biomaterials spontaneously organize into 3D stromal microenvironments during MatriSphere assembly in vitro

We sought to utilize the tissue mimetic diversity of SIS ECM to engineer 3D TME morphogenesis in vitro. Organoid methods, where isolated cells are seeded within a hydrogel (such as Matrigel), have traditionally been used to impart ECM signals in a 3D tumor context. However, hydrogel polymerization via monomer crosslinking does not involve resident cell participation during ECM assembly. Thus, we identified conditions to facilitate cancer cell and decellularized ECM co-assembly to build tumor tissues in vitro in high-throughput well plate systems. We evaluated two murine (MC38 and CT26) and one human (HT-29) CRC cell lines and found that microgram quantities (over a range of concentrations, 8-125 µg/mL) of decellularized ECM digest are assembled into an ECM dense stroma within MatriSpheres in ultra-low attachment culture (**Fig. 2a, Supplementary Fig. 2a)**. All three cell lines formed compact 3D spheroids as cells alone and as MatriSpheres with low concentration (125 µg/mL) SIS ECM dispersion supplementation (**Fig. 2b**). Purified Type I Collagen (matched to SIS ECM soluble collagen content) and Matrigel formation efficacy depended on specific cell line and ECM type. MC38 and CT26 spheroids formed with the addition of Type I Collagen and Matrigel whereas HT-29 cells did not, instead forming cell multilayer discs with Type I Collagen and unconsolidated clusters with Matrigel rather than a consistent, cohesive structure. As a control derived from stem cell spheroid models^16^, we validated that SIS ECM particles (**Supplementary Fig. 2b**) could be incorporated into tumor spheroid via sedimentation rather than cell mediated assembly in CRC cells. MatriSpheres had more consistent morphology and size than other ECM types (**Fig. 2b-c, Supplementary Fig. 2c**). Compared to cells alone spheroids (MC38: 770 ± 30 µm, CT26: 870 ± 14 µm, HT-29: 776 ± 30 µm), MatriSpheres diameters were larger with CT26 (919 ± 46 µm) and HT-29 (990 ± 63 µm), but smaller in size with MC38 (748 ± 41 µm) suggesting that there are cell line dependent effects of how they utilize ECM from their environment. Matrigel incorporation yielded the largest diameter structures but were variable in all 3 cell lines after 7 days, with diameters ranging from 968 ± 141 µm in HT-29 to 1,090 ± 56 µm in CT26 compared to cells alone spheroids.

**Figure 2:**
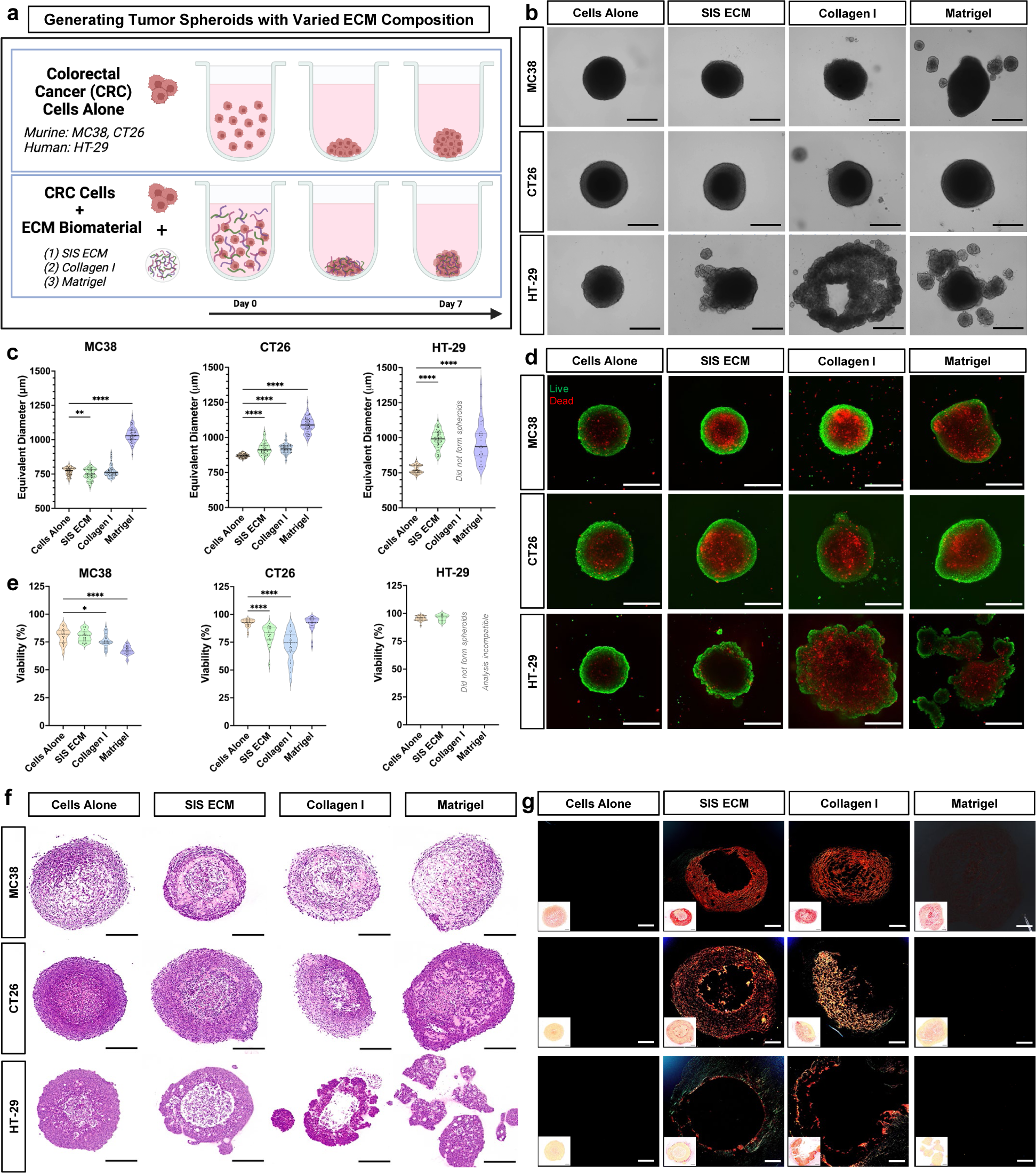
CRC cells assemble and organize ECM to form 3D MatriSpheres. a. Colorectal cancer cells are seeded as cells alone or with low concentration of ECM to generate spheroids in ultra-low attachment round-bottom 96-well plates for seven days of culture. b. Representative brightfield images of spheroids after 7 days of culture. Scale bars: 200 µm. c. Quantification of spheroid diameters after 7 days of culture (n ≥ 37 for groups that were quantified). d. Live-dead images of spheroids after 7 days of culture. e. Quantification of cell viability within the spheroids after 7 days of culture (n ≥ 13 for groups that were quantified) f. Representative H&E images of tumor spheroid sections demonstrates differences in micro-tissue organization. Scale bars: 200 µm. g. Picrosirius red (PSR) stained spheroids imaged with polarized light displays the presence of dense collagen fibers in SIS ECM and Type I Collagen groups. Scale bars: 100 µm. Statistics were calculated by a one-way ANOVA followed by a Tukey’s multiple comparisons test with significance values p < 0.05 is denoted with *, ≤ 0.01 with **, ≤ 0.001 with ***, and ≤ 0.0001 with ****.

We confirmed that MC38 and HT-29 MatriSpheres maintained a similar viability (74-99%) to cells alone controls (83-98%) but was lower for CT26 MatriSpheres (70-90%) (**Fig. 2d-e**). Other ECMs showed more pronounced decreases in variability, notably with Collagen I supplementation in MC38 and CT26 cultures and Matrigel with MC38. As expected from previous reports of 3D tumor spheroids^12^, we observed viable cells predominantly at the MatriSphere cortex with dead cells towards the center via live/dead staining. Furthermore, CellTiter-Glo quantification of metabolically active cells (**Supplementary Fig. 2d**) was generally consistent with spheroid size except for CT26 cells alone and with Matrigel, which had different diameters but were metabolically similar. We also enzymatically dissociated 3D spheroids and MatriSpheres at 7 days and found averages ranging from 13,000 to 97,000 cells per spheroid, which was greater than the initial seeding of 2,500 cells per spheroid showing net proliferation (**Supplementary Fig. 2e**). These results support that differences in diameters between ECM groups are due to effects on cell proliferation and survival rather than compaction or space occupied by the ECM alone.

We performed detailed histologic analyses using a modified tumor spheroid microarray^22^ and confirmed that SIS ECM is assembled and organized into collagen dense stroma-like regions within MatriSpheres after 7 days (**Fig. 2f**). ECM type impacted stromal morphology and composition of across CRC cell lines. We observed particularly dense ECM regions within certain conditions such as MC38 MatriSpheres, as shown by H&E and Masson’s Trichrome (**Supplementary Fig. 2f**). Picrosirius red (PSR) stained sections imaged with polarized light revealed that these stroma-like regions were indeed collagen-rich and formed birefringent fibrils (**Fig. 2g**). Consistent with morphologic analysis, collagen assembly was cell line-specific, with no detectable fibrillar collagen in cells alone spheroid conditions. This lack of endogenous ECM production highlights the need to provide ECM supplements for in vitro models. Concentrically-oriented large diameter collagen fibrils (red staining) were abundant in both the MC38 and CT26 cell lines when adding SIS ECM or Type I Collagen. Collagen in HT-29 MatriSpheres was enriched near the periphery in low diameter bundles (green staining) rather than the interior. Though Matrigel assembly was pronounced, it is composed of non-fibrillar collagens as confirmed by PSR staining. These observations recapitulate dense collagen dense stromal regions often pronounced in tumors from patients^23^. Taken together, we have shown that decellularized ECM organizes with CRC cell lines to create viable ECM MatriSpheres with a collagen rich stroma.

### MatriSphere formation is distinct from ECM hydrogel polymerization and traditional spheroid assembly

We observed that low ECM concentrations were sufficient to develop organized, collagen-dense microenvironments within MatriSpheres and that ECM composition altered 3D formation kinetics compared to cells alone. These ECM concentrations are well below those previously reported for hydrogel formation^24^, suggesting an alternative mechanism of ECM assembly that we investigated using live cell time lapse imaging (**Fig. 3a-b, supplementary videos**). Cells alone in ultra-low attachment conditions displayed similar spheroid formation behavior for all three cell lines; cells sediment at the bottom of the well, aggregate within an hour of seeding, and then proceed to grow outward within the first day (**Supplementary Vid. 1,5,9:** days 0-3). However, ECM addition altered kinetics and stages of the MatriSphere formation process (**Supplementary Vid. 2-4,6-8,10-11:** days 0-3). We classified the stages of MatriSphere assembly with SIS ECM as: (1) rapid formation of multiple smaller cell clusters within hours of seeding, (2) coalescence of these clusters via directed motion towards the center of the well to form a single larger MatriSphere over the following four to five days and (3) outward growth thereafter. Collagen I supplementation was similar to SIS ECM except with the HT-29 line which did fully coalesce at 7 days despite being composed of the same soluble collagen content as SIS ECM. Formation with Matrigel supplementation more closely resembled traditional spheroid assembly with cells alone with MC38 formation and demonstrated a very rapid and localized cluster formation phase with CT26 and HT-29 cells (**Supplementary Vid. 8 & 12**) that concludes within 24 hours. These results show distinct 3D formation characteristics that are dependent on ECM composition. Since HT-29 cells formed 3D structures more slowly than cells alone with each of the tested ECM biomaterials, we evaluated HT-29 assembly over longer timepoints comparing cells alone, SIS ECM at working concentrations, and Type I Collagen at a reduced concentration (25 µg/mL) (**Supplementary Fig. 3a**). SIS ECM MatriSpheres continued to evolve beyond 7 days, forming a more compact structure by day 10. Likewise, reduced Type I Collagen enabled single structure formation by 7 days whereas it was inhibited at higher concentrations. HT-29 spheroid and MatriSphere diameter and viable cell numbers increased over 7-21 days via CellTiter-Glo analysis (**Supplementary Fig. 3b-c**). This demonstrates that MatriSphere and ECM composition can be tailored to facilitate ECM assembly in more challenging cell lines, and that MatriSpheres are viable in long term cultures.

**Figure 3:**
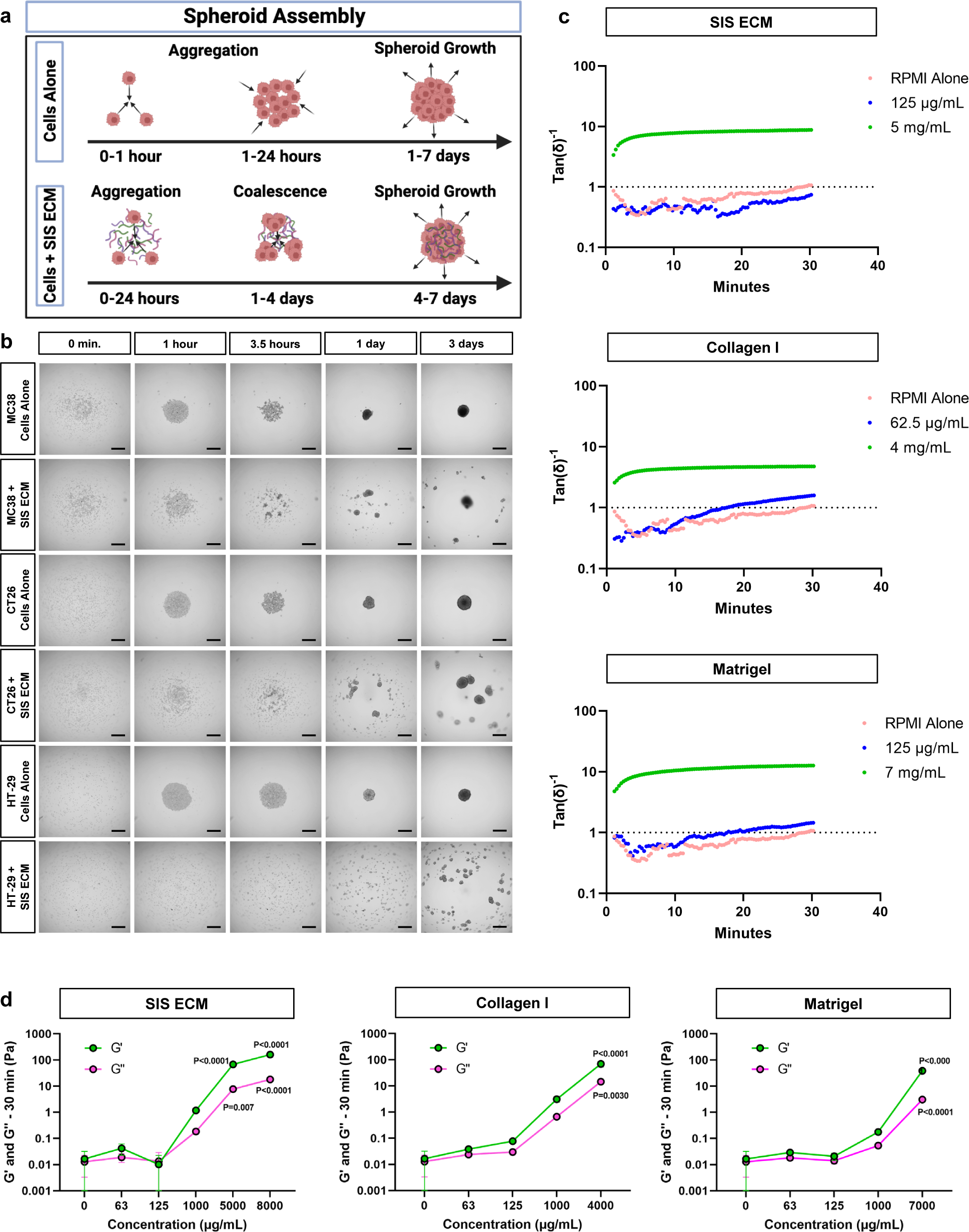
Cell-mediated ECM assembly delays formation kinetics. a. Schematic overview of cell assembly and spheroid formation mechanisms for cells alone and SIS-ECM MatriSpheres. b. Representative images of formation over the first 3 days of culture during time-lapse imaging. Scale bar: 500 µm. c. Tan(δ)^-1^ plotted as an indicator of gelation of ECM biomaterials (without cells) at working concentrations used in MatriSphere formation and traditional hydrogel formation measured by rheology (n ≥ 2 per group) d. Averaged storage (G’) and loss (G’’) moduli of ECM biomaterials at varied concentrations observed at 30 minutes after exposure to 37°C. A 2-way ANOVA with Dunnett’s multiple comparison test was used to assess significant differences (p-value < 0.05) within or between groups.

These complex MatriSphere formation behaviors and spontaneous ECM assembly motivated us to elucidate the differences between ECM organization during spheroid assembly from other well characterized mechanisms of gelation and to gain insights into whether tumor cells orchestrate this process. Our live cell imaging experiments did not show increased opacity during MatriSphere formation that is characteristic of bulk ECM hydrogel formation. Further, rheometry of ECM biomaterials confirmed that there was no spontaneous SIS ECM gelation with MatriSphere conditions in the absence of cells (125 µg/mL SIS ECM) and remained indistinguishable from media only controls. Only SIS ECM concentrations above 1,000 µg/mL resulted in stable and robust gelation defined by sigmoidal increase in storage and loss moduli (G’, G’’), and Tan(δ)^-1^ greater than 1 (**Fig. 3c,d**). Similar rheologic behaviors were found for Type I Collagen and Matrigel, and G’ remained below 0.05 Pa for each biomaterial at working concentrations. These results were supported by turbidimetric gelation kinetics assays showing no difference in turbidity at working concentrations of the SIS ECM and Matrigel below 125 µg/mL and Collagen I below 62.5 µg/mL compared to media alone **(Supplementary Fig. 3e)**. In contrast, turbidity of ECM biomaterials increased with ECM concentrations above 125 µg/mL **(Supplementary Fig. 3f)**. These results suggest that ECM biomaterial incorporation is not a passive process driven by ECM properties alone and is likely cell-mediated since we only detect ECM assembly within the final 3D cell structure. We hypothesize that the cells slowly concentrate the ECM within the primary spheroid and create an extratumoral ECM mesh that connects and envelopes small cell clusters, guiding spheroid assembly. The lack of visible turbidity and rheologic gelation suggest a cell mediated and highly localized ECM assembly process, as an emergent phenomenon that is independent of ECM hydrogel formation.

### Decellularized ECM promotes spatial phenotypic organization and heterogeneity within CRC MatriSpheres

Our histologic analysis (**Fig. 2**) showed that decellularized SIS ECM assembled into distinct networks of collagen rich stroma during MatriSphere formation, and this assembly required cell participation (**Fig. 3**) creating heterogenous intraspheroidal morphologies. Intratumoral heterogeneity arises from both cancer cell intrinsic genetic variation and environmental factors that can produce discrete regions of hypoxia and ECM density that in turn influence cancer cell mesenchymal transition, proliferation, migration, and drug resistance^25–27^. Therefore, we sought to characterize spatial organization of cellular features within MatriSpheres, and whether these correlate with ECM composition using multiplex fluorescence histology. We defined ECM rich regions by staining fibrillar collagen with a collagen hybridizing peptide (CHP, **Fig. 4a-b,e, Supplementary Fig. 4a,c-i**)^28^. The large dynamic range of the CHP (**Supplementary Fig. 4c**) allowed us to infer collagen density across ECM conditions and cell lines with variable ECM incorporation: the MC38 line with the greatest collagen density, CT26 was intermediate, and HT-29 was lowest in MatriSpheres (**Supplementary Fig. 4c,d,f,h**). As expected, CHP intensity was greater within MatriSpheres and with Type I Collagen supplementation rather than Matrigel, which has little fibrillar collagen (**Fig. 1d**). Collagen rich morphology was consistent with our PSR results and enabled fluorescent co-localization with cell-cell adhesion cadherin junctions, E-cadherin (E-CAD) and N-cadherin (N-CAD), proliferation marker (Ki67), and Carbonic Anhydrase IX (CA IX) as an environmental feature. We found that the organization of CRC cells and ECM within MatriSpheres dictated cadherin expression patterns, though overall expression was similar across conditions. CT26 cells were the exception, with increased N-CAD expression for each of the ECM-incorporated groups compared to cells alone and the highest expression with Matrigel (**Fig. 4c**). Spatially, N-CAD positive cells were enriched at the periphery in the cells alone control groups and Matrigel-containing MC38 and CT26 spheroids.

**Figure 4:**
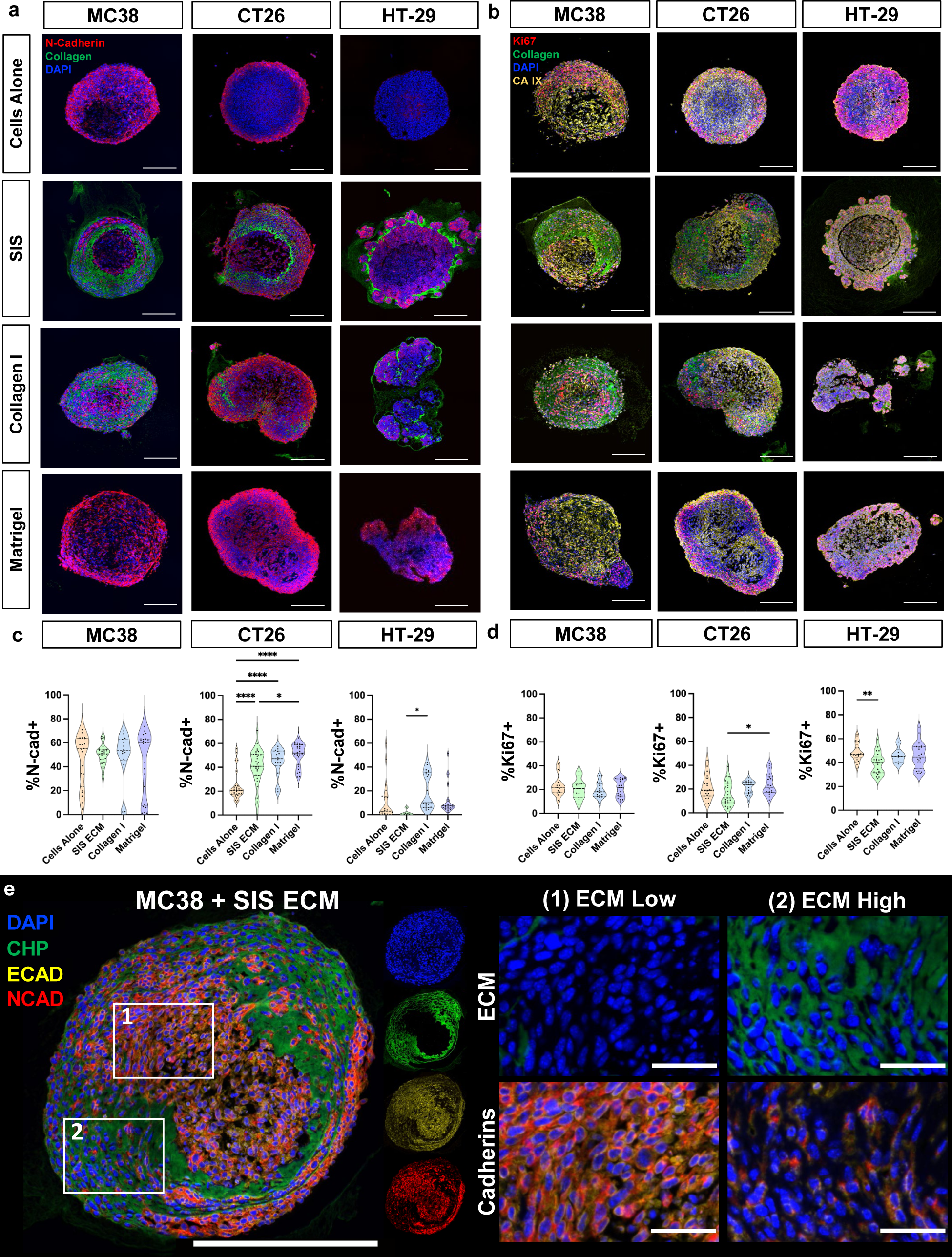
CRC MatriSpheres display heterogenous phenotypes with enhanced cell-ECM interaction. a. Multiplex fluorescence staining for CHP (denatured fibrillar collagen, green), N-CAD (red), and DAPI (blue) within MatriSphere histologic sections. Scale bar: 200 µm. b. Multiplex fluorescence staining for CHP (denatured fibrillar collagen, green), Ki67 (proliferation, red), CA IX (hypoxia, yellow), and DAPI (blue) within MatriSphere histologic sections. Scale bar: 200 µm. c. Quantification of N-CAD expression in representative MatriSphere sections. Statistics were calculated by a one-way ANOVA followed by a Tukey’s multiple comparisons test. d. Quantification of Ki67 expression in representative MatriSphere sections. Statistics were calculated by a one-way ANOVA followed by a Tukey’s multiple comparisons test. e. Representative MC38 + SIS ECM MatriSphere section stained for CHP (denatured fibrillar collagen, green), E-CAD (yellow), N-CAD (red), and DAPI (blue). Cadherin expression is higher in ECM low regions than in ECM low regions. Full image scale bar: 250 µm. Inset image scale bar: 50 µm. Statistical significance where p < 0.05 is denoted with *, ≤ 0.01 with **, ≤ 0.001 with ***, and ≤ 0.0001 with ****.

Furthermore, cadherin expressing cells (either E-CAD or N-CAD) segregated into cell-dense tumor nests within MatriSpheres whereas cadherin negative cells were enriched in ECM dense regions within MC38 MatriSpheres (**Fig. 4e**). This suggests that these cells may rely more on cell-matrix interactions rather than cell-cell interactions when there is a high amount of ECM present though it is unknown whether this represents a phenotypic switch or preferential organization during assembly. This organization was consistent in CT26 MatriSpheres **(Supplementary Fig. 4a)**. HT-29 MatriSpheres had the lowest area of high ECM density, which resulted in predominantly cadherin positive cells. ECM dense tumor stroma has been implicated in affecting nutrient diffusion and sequestering factors such as cytokines and growth factors^29^, which could impact the ability of cells to proliferate or increase hypoxia within the spheroid. Decellularized tissues such as SIS ECM preserve this complexity and have been previously shown to maintain the ability to bind such soluble factors^30^. We used Ki67 as a marker for proliferation (**Fig. 4b,d**) to distinguish actively proliferating from quiescent cells and carbonic anhydrase 9 (CA IX, **Fig. 4b, Supplementary Fig. 4b**) as a hypoxia marker associated with poor prognosis in CRC patients^31^. All conditions expressed CA IX indicating hypoxia, with no differences between groups. The total number of proliferating cells was also similar between ECM conditions, and were primarily cell line dependent (20%, 30%, and 50% for MC38, CT26, and HT-29, respectively). All spheroids contained similar levels of proliferative cells, with minor differences indicating that ECM conditions did not affect proliferation alone during homeostatic culture conditions. These results show that ECM utilization and organization are cell line dependent, suggesting dynamic cell-cell and cell-ECM interactions that determines the final structure and phenotype.

### CRC MatriSpheres’ unique transcription profiles recapitulate in vivo tumor heterogeneity

We determined the effect of SIS ECM on CRC spheroid phenotype after MatriSphere formation (7 days) using bulk RNA sequencing in murine (MC38 and CT26) and human (HT-29) cell lines. MatriSpheres and traditional spheroids separated into distinct phenotypic clusters for each cell line via principal component analyses (PCA), identifying 112, 165, and 28 high-significance differentially expressed genes (DEGs, adjusted P-val < 0.01 and FC > 2) in MC38, CT26, and HT-29 MatriSpheres, respectively (**Fig. 5b**). These results indicate that MatriSpheres are transcriptionally distinct from traditional cells alone spheroids. Specific gene regulation patterns in MatriSpheres were cell line dependent. The top 10 DEGs in MC38 MatriSpheres were mostly downregulated including several ECM genes (*Col1a1, Col28a1, Dmp1, Vtn, Aspn*) (**Fig. 5c**); however, increased expression of the long non-coding RNA *Gm20427* accounted for the highest fold change in MC38 spheroids. In contrast to MC38 spheroids, the top DEGs from CT26 spheroids were largely upregulated in response to SIS ECM. The highest absolute fold changes for CT26 included genes associated with CRC aggressiveness and invasion, including *Tnfrsf11b*, which codes for osteoprotegerin, *Mmp13*, and zinc finger family of proteins (*Zfp704, Zfp936*)^32,33^. For HT-29 cells, increased *Plat* (plasminogen activator, tissue type) was the largest fold change (4.73) in MatriSpheres, along with upregulation of multiple APOBEC (apolipoprotein B mRNA-editing enzyme, catalytic polypeptide) genes (*Apobec3g, Apobec3c, Apobec3f*) within the top 10. APOBEC3s are known to play to a role in cancer mutagenesis and APOBEC3G is highly expressed in colon cancer^34^.

**Figure 5:**
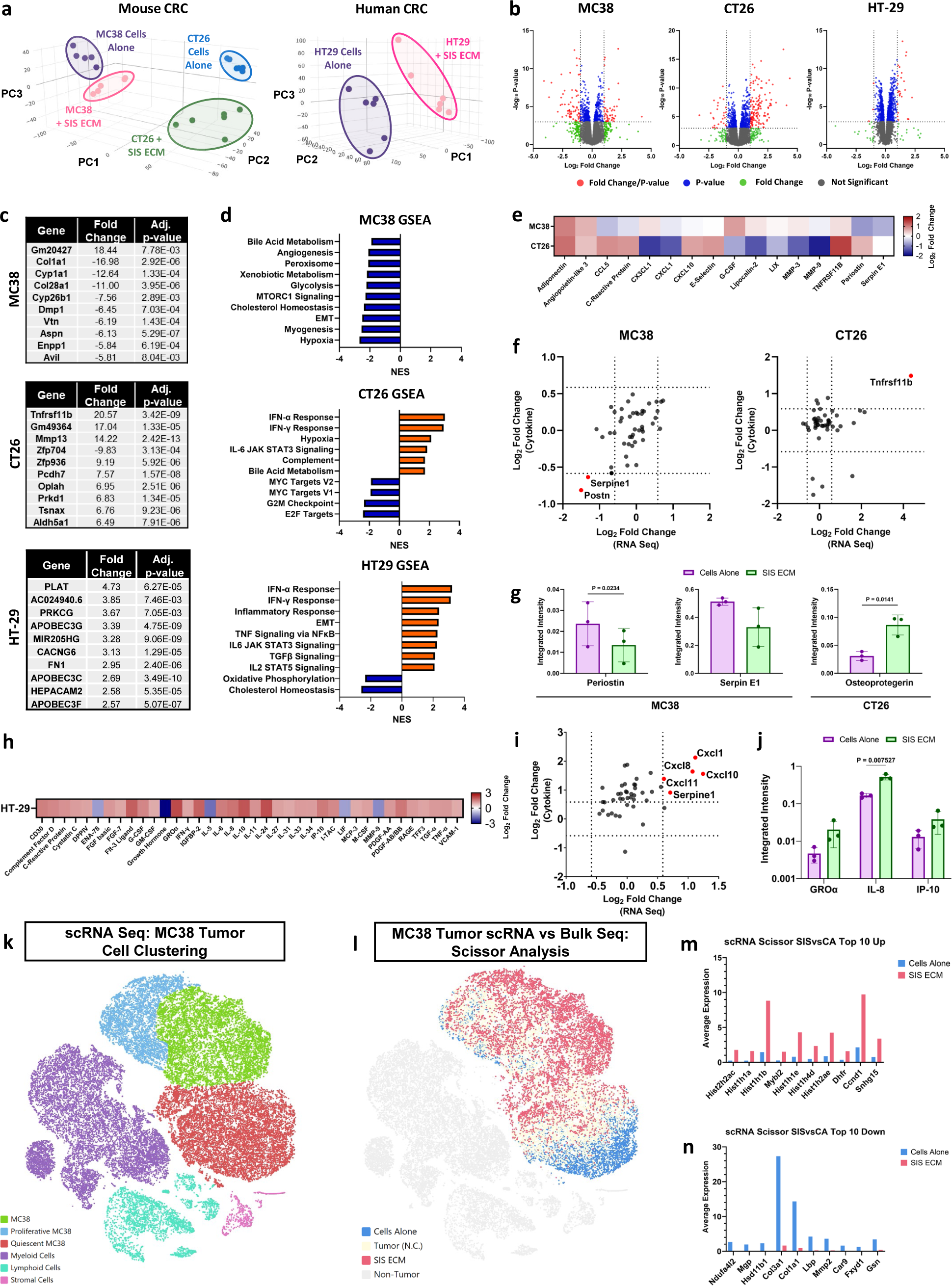
MatriSpheres alter CRC transcriptome and secretome capturing in vivo tumor heterogeneity. a. PCA plots clustering RNA Seq expression data of both mouse and human CRC spheroids cultured with and without SIS ECM at day 7. b. Volcano plots of differentially expressed genes across all three cells line as compared to cells alone spheroid controls. c. Top 10 genes for each CRC cell line by fold change. d. Top 10 gene set enrichment based upon GSEA. Significantly upregulated (orange) and downregulated (blue) gene sets. e. (MC38 and CT26), h. (HT-29) Heatmap displaying top cytokine fold change between cell culture supernatant from MatriSpheres compared to cells alone spheroids at Day 7. f. (MC38 and CT26), i. (HT-29). Modified correlation plot identifying similarities between conserved transcriptome and secretome signatures g. (MC38 and CT26), j. (HT-29) Chemiluminescent intensity values of correlated proteins. k. tSNE plot of cell clusters identified within in vivo MC38 tumors based upon scRNA Seq expression data. l. tSNE plot comparing bulk RNA Seq expression level data from in vitro MC38 MatriSpheres and cell alone spheroids to scRNA Seq expression of in vivo MC38 cancer cells. m. Top 10 upregulated and n. downregulated genes defining the high confidence correlation of in vitro MC38 tumor spheroids with in vivo cancer cells. Significant differences were defined as p-value < 0.01 for RNA Seq data.

We applied gene set enrichment analysis (GSEA) to infer broad phenotypic regulation in MatriSpheres. The top 10 most notable pathways in MC38 MatriSpheres (highest normalized enrichment score, NES), were all negatively correlated relative to cells alone (**Fig. 5d**). The largest impact was found in the suppression of the hypoxia signaling pathway. Additionally, several metabolism-related pathways were downregulated in response to SIS ECM. Within the hypoxia gene set, there were 68 leading edge genes, including *Serpine1*, which is a serine proteinase inhibitor that blocks plasminogen activation and limits fibrin degradation **(Supplementary Fig. 5g)**. High expression of serpin E1 has been implicated in CRC TME remodeling, specifically aiding immune cell infiltration^35^. GSEA for CT26 MatriSpheres revealed an inflammatory response led by the upregulation of interferon signaling pathways such as IFN-α and IFN-γ, *Sp140* an activator of STAT1, as well as other IFN regulatory factors (*Irf9*, *Irf7, Irf1*) and IFN-induced transmembrane proteins (Ifitm3, Ifitm1, Iftim2) **(Supplementary Fig. 5h)**^36–38^. Similar to CT26 spheroids’ response, we found a positive enrichment of several pathways within HT-29 spheroids involved in inflammation, including IFN-α, IFN-γ and inflammatory response signaling pathways. We also noticed an increase in the epithelial to mesenchymal transition (EMT) pathway, which is an indicator of metastatic potential, and negative regulation of cholesterol homeostasis. IFN-α response pathway was the most perturbed gene set with 61 leading edge genes and a NES of 3.21 **(Supplementary Fig. 5i)** and the IFN response gene (*Ifitm1*) was the highest ranked gene, which has been proposed as a marker of advanced clinical stage and poor survival in CRC patients^39^. We inferred more generalizable gene regulation across 8 shared DEGs between MC38 and CT26 MatriSpheres **(Supplementary Fig. 5c)**. *Mmp13* expression was upregulated and both *Aspn* and *Dlx2* were downregulated in mouse cell lines **(Supplementary Fig. 5d)**.

Many of the transcriptional changes in MatriSpheres corresponded with a functional CRC cell secretory phenotype and increases in 16 cytokines compared to traditional spheroids (FC > 1.5) increased (**Fig. 5e**). In MC38 MatriSpheres, we found downregulation of both Serpin E1 (*Serpine1*) and Periostin (*Postn*), which agrees with RNA sequencing (**Fig. 5f,g**). For CT26 spheroids, only osteoprotegerin (Tnfrsf11b) was significantly increased in MatriSpheres. In the human HT-29 line, we found 38 cytokines had an absolute fold-change ≥ 2 (**Fig. 5h**). However, only three pro-inflammatory cytokines (GROα, IL-8, and IP-10) correlated in both secretion and RNA sequencing analyses (**Fig. 5i,j**). These cytokines have been identified as essential factors in the wound healing response and have been cited as pro-tumor markers leading to poor survival in CRC patients.

A primary aim of this study was to establish an in vitro model of the TME that specifically recapitulates in vivo ECM conditions. We therefore sought to benchmark MatriSpheres using in vivo MC38 tumors analyzed via single cell RNA sequencing (scRNAseq) and to characterize MC38 cancer cells in their native stromal and immunocompetent environment. We broadly defined three primary clusters: MC38 Tumor, Immune and Stromal cells (**Fig. 5k**). Further, we identified expected immune and stromal microenvironment subpopulations, and found MC38 cells clustered into proliferative and quiescent MC38 cells based on proliferation markers (*Mki67, Pcna, etc*). We found that SIS ECM MatriSpheres recapitulated a subpopulation of in vivo MC38 cells using Scissor analysis^40^, which determined a phenotypic correlation score compared with in vitro MatriSpheres and traditional MC38 spheroids with cells alone (**Fig. 5l**). Out of 20,796 MC38 tumor cells, 34.5% positively correlated with MatriSpheres containing SIS ECM and 18.6% for Cells Alone spheroids. Expression profiles of MC38 SIS ECM MatriSpheres were predominantly clustered within medium to highly Proliferative MC38 cell populations (P-MC38), while MC38 Cells Alone spheroids correlated almost exclusively with Quiescent MC38 cells (Q-MC38). This analysis indicates that SIS ECM MatriSpheres can drive unique CRC phenotypes observed within primary in vivo tumor cells. A DEG analysis between these distinct MC38 cancer cell populations in vivo showed that cells more similar to SIS ECM MatriSpheres upregulated histone genes (Hist2h2ac, Hist1h1a, Hist1h1b, Hist1h1e, Hist1h4d, Hist1h2ae) and proto-oncogene, Mybl2, and downregulated ECM protein and remodeling genes (Col3a1, Col1a1, Mmp2). Aberrant histone modification and expression can disrupt DNA folding and replication dynamics, which may have several consequences in cancer^41,42^. Mybl2 overexpression has been linked to reduced CRC patient survival, while knocking down Mybl2 expression in SW480 cells dysregulated cell cycle and proliferation^43^. These results suggest that in vitro phenotypes may improve modeling of specific cancer cell subpopulations.

### CRC MatriSpheres conserve upstream regulators linked to TME target genes

We next identified putative pathway activation trends with SIS ECM MatriSpheres utilizing Ingenuity Pathway Analysis (IPA). We found MatriSpheres most strongly regulated signaling pathways annotated as cancer-associated, immune-associated, and metabolism-associated across cell lines and thus were the focus of further analysis. The most significant change in MC38 MatriSpheres was decreased activity in metabolism-associated pathways, specifically cholesterol biosynthesis, which agreed with the GSEA findings **(Supplementary Fig. 6a)**. Cholesterol metabolism has several potential implications within the TME and this dysfunction within cancer cells could be exploited for therapeutic benefit^44^. The top DEGs in each category showed decreases in collagen expression (*Col1a1*, *Col28a1*), immune-associated pathways involving *Il2rg* and *Rras,* and metabolism-associated genes Mvd and Dgat2 **(Supplementary Fig. 6b)**^45^.

CT26 MatriSpheres likewise predicted immune modulation resulting from ECM interactions, activating pathways that include IFN signaling and cytokine storm that may indicate pro-inflammatory signatures **(Supplementary Fig. 6c)**. Cancer-associated pathways influenced by SIS ECM in CT26 spheroids were driven by an upregulation in 2’-5’ oligoadenylate synthetases 2 and 3 (*Oas2*, *Oas3*) **(Supplementary Fig. 6d)** also associate with immune regulation, which are enzymes activated by IFN signaling cascades and has been shown to establish an immunosuppressive microenvironment^46^. IFN-related genes (Prkd1 and Irf7) were also enriched in the presence of SIS ECM. Protein kinase D1 (Prkd1) interacts with β-catenin to control proliferation in CRC^47^ and elevated Irf7 expression correlates with greater immune cell-infiltrates and poor survival in CRC patients^48^.

HT-29 MatriSpheres showed strong predicted activation of cytokine production, fibrosis, and wound healing, as well as a significant inhibition of cholesterol biosynthesis signaling **(Supplementary Fig. 6e)**. We found both Prkcg (protein kinase c gamma) and Fn1 (fibronectin) upregulated within the fibrosis signaling pathways **(Supplementary Fig. 6f)**. Prkcg has been shown to increase CRC cell migration, whereas its inhibition increases cell adhesion and proliferation^49^. Elevated levels of fibronectin have also been implicated in poor prognosis and survival in CRC patients^50,51^. IL-8 signaling contained the highest fold change from immune-associated pathways with significant increases in both Cxcl1 (CXC motif ligand 1) and Cxcl8 (CXC motif ligand 8). Cxcl1 and Cxcl8 initiate neutrophil recruitment during wound healing, but they also play critical role in CRC progression and metastasis by suppressive myeloid cell recruitment^52^.

To summarize the conserved ECM stroma effects on CRC cell expression, we discovered three shared pathway activity patterns predicted across all 3 cell lines: Superpathway of Cholesterol Biosynthesis, Role of Chondrocytes in Rheumatoid Arthritis, and Pyroptosis Signaling (**Fig. 6a**). Additionally, 22 other pathways were shared between at least 2 out of 3 cells lines with high activation significance (|z-scores| ≥ 3) (**Fig. 6b**). CT26 and HT-29 spheroids showed the most similarity within the cancer and immune-associated pathways 61% (11/18) and 55% (12/22) of the total pathways shared between them. The most significant pathways between the CT26 and HT-29 were related to IFN signaling, specifically the Role of PKR in IFN Induction and Antiviral Response (Avg. z-score: 3.46) and IFN Signaling (Avg. z-score: 3.38). In general, MC38 pathway activation was less similar to CT26 and HT-29, however, several metabolism-associated pathways were shared between MC38 and HT-29. Cholesterol metabolism has been shown to dramatically alter the TME and could be exploited to improve cancer therapies^53^.

**Figure 6:**
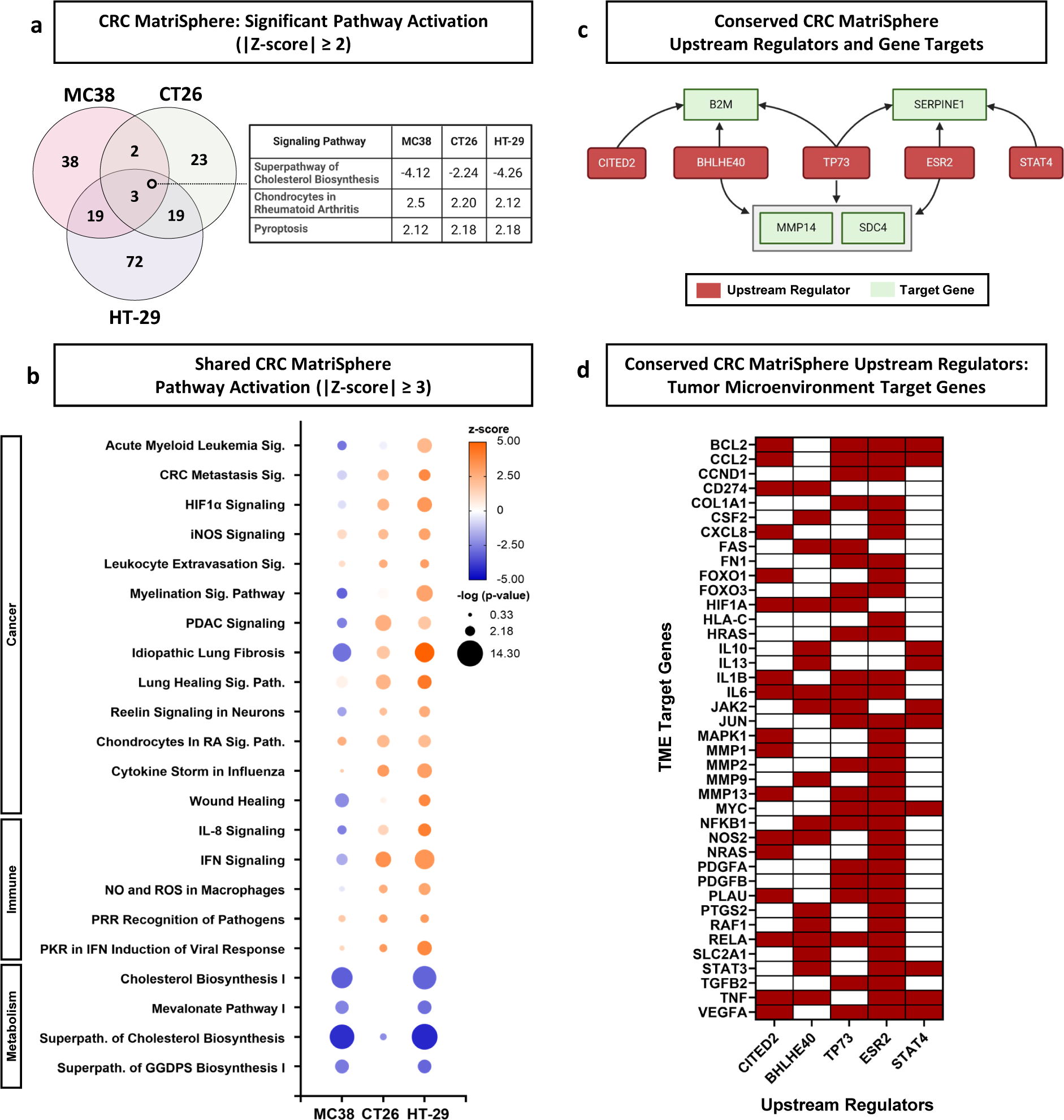
Conserved ECM effects on CRC pathway activation and potential TME target genes. a. Venn diagram showing the number of predicted pathways and their activity based upon RNA transcription patterns across CRC MatriSpheres. Filtered by |z-scores| ≥ 2. The table highlights the 3 significantly overlapping pathways and the corresponding z-scores across all CRC cell lines. b. Bubble plot displaying shared significant pathways. Pathways shown contained two or more CRC cell lines with |z-scores| ≥ 3. c. Flow chart of interactions between IPA predicted upstream regulators and target genes conserved between all CRC cell lines. d. Heatmap of target genes within the IPA tumor microenvironment pathway that have interactions with the 5 identified upstream regulators.

We probed pathway activation profiles to identify common predicted regulators and gene targets predicted in CRC MatriSpheres across all 3 cell lines. We found 10 upstream regulators predicted and measured by IPA software corresponding with 35 overlapping genes. From that list, we filtered based upon significance (p-value < .001 and |z-score| > 2) and identified 10 shared upstream regulators across CRC cells lines **(Supplementary Fig. 6g)**. We narrowed these interactions to 5 conserved upstream regulators CITED2, BHLHE40, TP73, ESR2 and STAT4 that directly interacted with 4 common gene targets B2M, SERPINE1, MMP14 and SDC4 (**Fig. 6c; Supplementary Figs. 6h,i**). We hypothesize that these regulator-target interactions provide insights of possible pathways that are dependent upon ECM interactions that are otherwise absent in traditional spheroid models. Across all 3 CRC MatriSpheres cell lines, IPA predicted CITED2 gene inactivation with strong confidence, which would indicate this regulator may have a similar signal transduction stimulated by SIS ECM. CITED2 (Cbp/P300 Interacting Transactivator with Glu/Asp Rich Carboxy-Terminal Domain) is a transcription regulator that acts as a competitive inhibitor to HIF-1α signaling. It has also been suggested that CITED2 can alter activation states of macrophages^54^, regulate cell proliferation^55^, and mediate colorectal cell invasion^56^. Lastly, we cross-referenced these shared regulators to determine their role in the TME signaling pathway and found 40 target genes that associate with at least 2 upstream regulators, out of 89 with at least one regulator interaction (**Fig. 6d; Supplementary Fig. 6j**). These findings suggest that SIS ECM or stromal interactions in MatriSpheres influence CRC signaling pathways beyond the transcriptional level, which may lead to broad TME regulation.

## Discussion

In this study, we developed MatriSpheres; a 3D culture system that incorporates orthotopic decellularized ECM within spheroids through cell-directed self-assembly. This spontaneously creates a tumor-like ECM stroma that induces phenotypic changes mimicking in vivo tumor heterogeneity. This facile and reproducible approach displays characteristics of tumor morphogenesis and is rheologically and structurally distinct from traditional hydrogel formation used in organoid methods. This model system displayed a dynamic reciprocity between CRC cells and a proteomically complex intestinal ECM. Specifically, the cells organized the partially-solubilized matrix, and in turn, the integrated matrix influenced CRC MatriSphere transcriptome and secretome phenotypes. Gene set enrichment and pathway analyses showed that SIS ECM influenced matrix remodeling, immune cell signaling, cell cycle, and lipid metabolism, which each play critical roles in cancer progression. Most notably, in vitro MatriSpheres demonstrated improved correlation with in vivo CRC cells over traditional spheroids. MatriSpheres provide a unique tool to seamlessly enhance spheroid complexity as well as their applicability for disease modeling and high-throughput therapeutic screening.

A significant finding of this study was that MatriSpheres containing decellularized ECM biomaterials enhance CRC spheroid ECM diversity and morphological complexity. In vitro tumor cultures have been unable to reliably predict therapeutic responses observed in the clinic, which is hypothesized due to a lack of TME hallmarks and do not reflect tumor heterogeneity. More complex 3D in vitro models (spheroid, organoid, and 3D bioprinting) and in vivo models (patient-derived xenografts) have been used to capture the complexity of solid tumors. However, with increasing complexity comes a greater financial burden, reduced analysis throughput, and higher variability, which could limit their implementation and highlights the importance of a balance of practicality and predictiveness. Therefore, our initial objectives in MatriSphere development were to (1) bridge the gap in 3D model complexity between traditional spheroids and ECM-competent organoids, (2) facilitate orthotopic ECM composition using decellularized tissues, and (3) maintain the ease of standard spheroid workflows.

Traditional spheroids grown in low-adherence conditions are an economical option to recapitulate some features of solid tumors such as diffusion gradients of oxygen and nutrients. In addition, they provide an opportunity to conduct assays and downstream analyses of individual objects per well which reduces sample to sample variability. However, spheroid assembly is not a given, as many cell lines and primary cells are unable to form compact spherical structures with cancer cells alone. Additionally, most formed tumor spheroids lack a mature ECM stroma, which is a hallmark of many advanced tumors. Therefore, in the absence of a robust ECM stimulus, spheroids may fail to capture the complex phenotypic and functional in vivo tumor responses. Organoids improve upon several aspects of spheroid cultures through the addition of tissue-derived ECM biomaterials, such as Matrigel. The presence of basement membrane ECM proteins stimulates cancer stem cell growth, guides cell organization and induces phenotypic similarities to tumor cells^57,58^. Although organoids are a useful tool for tissue modeling, they have several limitations, such as specialized media formulations (e.g. growth factors), potential outgrowth of normal cells, and a lack of TME components (e.g. fibrillar ECM proteins and stromal cell types)^59^.

A benefit of MatriSpheres is their modular design, which allows cancer cells to assemble a multitude of ECM types, derived from different tissue sources, to match a tumor of interest. Every tissue in the body possesses a unique ECM composition and architecture, including intestinal tissues in which CRC arises. Decellularization has allowed researchers to isolate tissue-specific ECM scaffolds that contain unique biochemical composition^60^. The ECM can be processed into multiple forms and has been used clinically for its innate regenerative properties^61,62^. Previously, spheroids made from human adipose derived stem cells or breast cancer cells were augmented with ECM particles derived from collagen I, bone, brain, cartilage, adipose, lung, and spleen and showed discordant cell responses to varied ECM biomaterial composition^16^. Another study showed that orthotopic decellularized intestinal ECM hydrogels could improve gastrointestinal organoid maturation and function suggesting that Matrigel alone may not be the most effective ECM biomaterial for tumor modeling^63^. Our proteomics analysis indicated that Matrigel represents a quarter of the matrisome homology as orthotopic intestinal ECM when compared to MC38 tumor ECM, supporting the use of an alternative mass producible ECM source. Another unique advantage of MatriSpheres is their ability to enable tumor cells to rapidly develop a mature ECM compartment by self-assembly rather than *de novo* synthesis that is otherwise absent due to the lack of specialized cell types, such as fibroblasts, which are professional ECM generators^64^. In traditional cells alone spheroid models, we find small amounts of matrix deposited over short time scales (2-7 days) and are generally devoid of fibrillar collagens that provide structural and mechanical support for virtually all solid tumors^65^. This was confirmed by single cell analysis of in vivo MC38 tumor cells which showed relatively low matrisome expression compared to stromal cells. In other words, this method provides user-defined ECM composition without the need for heterotypic cell cultures that may prevent the study of direct cancer cell responses to their changing microenvironment. Though CRC cells likely do not generate the bulk ECM stroma of tumors, recent studies have shown their contributions create a highly localized pericellular matrix that promotes drug resistant phenotypes^66^. This suggests the importance of ECM in tumor modeling but requires further investigation of cell contributions toward matrix deposition in addition to the observed stroma formation from supplemented ECM biomaterials.

The primary objective of a 3D tumor culture system is to recapitulate in vivo phenotypes under controlled in vitro conditions that are not possible in simplified models. Thus, we asked whether MC38 MatriSpheres would provide a tumor-like niche that would drive similar responses to in vivo MC38 cancer cells. We found that MatriSpheres induced unique transcriptomic phenotypes that correlated with multiple cancer cell subpopulations identified in vivo. Interestingly, single cell Scissor analysis^40^ revealed that in vivo MC38 tumors stratified into subpopulations that matched either MatriSpheres or cells alone spheroids. Our results suggest that a single culture condition may never fully capture in vivo tumor heterogeneity and that there is utility in examining tumor spheroids that are both ECM rich and poor. This may address a challenge in treating solid tumors where cancer cells populate heterogeneous TME niches^67,68^. These characteristics have hindered clinical efficacy of targeted cancer therapies based upon genetic mutations alone rather than the influence of the complete TME^69^. These findings illustrate the importance of utilizing ECM biomaterials for solid tumor modeling to mimic the diverse cancer cell phenotypes observed in vivo.

We found that the concept of dynamic reciprocity was a promising framework to describe both ECM organization within MatriSpheres and subsequent phenotypic regulation. Dynamic reciprocity, coined several decades ago by Bissell et al., describes the bidirectional relationship between cells and their extracellular environment^70^. The first principle of dynamic reciprocity was at work during MatriSphere formation, as cancer cells directed assembly of the ECM dispersion into defined networks encapsulating small cellular clusters, or nests. Distinct differences in ECM organization were observed between cell lines thus demonstrating cell-specific ECM utilization. For example, both MC38 and CT26 lines showed intraspheroidal ECM uptake; however, MC38 MatriSpheres displayed regions of hyper densified collagen fibers in contrast to a more even distribution of thinner collagen fibers in CT26 spheroids. In contrast, HT-29 cells preferentially assembled a peri-spheroidal collagen barrier with limited ECM assembly toward the spheroid core. In addition to cell-dependent ECM assembly, the second principle of dynamic reciprocity was evident wherein each CRC line responded differently to ECM via changes to their transcriptomes and secretomes. An advantage of MatriSpheres compared to organoids is the ability to decouple the contribution of ECM in 3D cultures by comparing to traditional spheroids. We identified multiple ECM-driven changes in CRC cells, such as matrix remodeling^71^, immune regulation, cell cycle and cholesterol metabolism. ECM-associated gene expression was particularly interesting and reinforces dynamic reciprocity. MatriSpheres typically downregulated structural ECM proteins (e.g. Collagens) and enhanced ECM protease (e.g. matrix metalloprotease) expression. We found that ECM engagement affected other cancer cell-intrinsic gene expression programs that potentially shape the TME including immunomodulation of inflammatory gene signatures and cytokines. Namely, Type I and II IFN-associated genes were heavily upregulated in CT26 and HT-29 spheroids in response to SIS ECM. To date, there has been a vast amount of literature documenting the important role of IFN signaling in altering tumor homeostasis and responsiveness to immunotherapy^72,73^. This inflammatory profile was supported by the enhanced cancer cell secretion of pro-inflammatory cytokines (GROα, IL-8 and IP-10) within HT-29 spheroids. This would indicate that the ECM is inducing cancer cells to recruit immune cells to the tumor site, but it is unknown whether this recruitment would polarize these cells toward a pro- or anti-tumor phenotype. We observed that ECM also played a causal role in cancer cell metabolism, robustly inhibiting cholesterol metabolism pathways across spheroids in each of the tested CRC cell lines, but most prominently in MC38 and HT-29 spheroids. Cholesterol metabolism is a critical aspect of cellular maintenance and has become an intriguing target for cancer therapy^74^. Although no single gene associated with this pathway was substantially altered, the collective downregulation of these genes resulted in the highest activation score among all pathways. This result predicts that the ECM within the spheroid significantly modifies the lipid biosynthesis machinery within cancer cells and could provide a model for studying the mechanism of ECM regulation of cancer cell metabolism^75,76^. MatriSpheres highlight the influence of cancer cells on the TME as well as their plasticity and capacity for being manipulated by the ECM^77^.

An unexpected finding in the present study was the stark difference between MatriSphere assembly and existing 3D culture models. Though the MatriSphere formation mechanism was not fully determined, we characterized several unique aspects of cell-ECM engagement. For example, ECM assembly is not simply hydrogel formation, but rather is an emergent phenomenon that concentrates ECM molecules at sub-gelation thresholds within intercellular spaces, creating a dense ECM-rich stroma. Cells only spheroids aggregate almost exclusively through cell-cell adhesions due to the low ECM abundance, which allows the cells to form quickly and homogeneously within the 3D structure. In contrast, SIS ECM fibers are slowly concentrated and encapsulate small clusters of cells that are pulled into a single dominant spheroid. It is not clear whether this process can be described as intercellular gelation^78^; however, our results demonstrate that the cells are active participants in not only aggregating into spheroid nests, but also the concentration and co-assembly of decellularized ECM from the surrounding environment. Rheological testing and live cell imaging confirmed that in the absence of cells, ECM concentrations at or below 125 µg/mL failed to form visible crosslinks. This highlights the importance of the cells in triggering ECM assembly and organization within MatriSpheres. Previous studies of synthetic peptide hydrogel precursors provide a precedent for cancer cell mediated matrix assembly and suggests a role for cancer cell produced fibronectin^79^. This process is reminiscent of ECM deposition, organization, and maintenance, which is poorly understood during in vivo tumor morphogenesis. It is likely that ECM receptors such as integrins direct spatial ECM organization, much like cell-cell junction proteins have been shown to drive spheroid organization, as explained by the differential adhesion hypothesis^80,81^. This framework may be useful to describe MatriSpheres, where integrin-ECM and ECM-ECM interactions compete to lower the free energy of the final system. We found that cells expressing low levels of both N-CAD and E-CAD were more likely to assemble in ECM rich regions, suggesting a driving force for this organization. Further analysis of these ECM assembly characteristics may provide insights into tumor morphogenesis.

3D model scalability and reproducibility are important considerations for pre-clinical applications, and the facile methods for MatriSphere generation demonstrate several practical advantages over more complex systems such as organoids or bioprinting. These methods are suitable for high throughput analysis in both 96- and 384-well formats **(Supplementary Fig. 7**), similar to traditional spheroids. Importantly, they can be formed from minute quantities of decellularized tissues. Even at the highest concentrations used in this study, only 12.5 µg (dry wt.) was necessary per 100 µL well. Commercial ECM scaffold production in the medical device industry is expensive, and this method uses this material sparingly where 1g of ECM would support a theoretical maximum 80,000 96-well conditions. This is 8-10 fold less material than creating ECM hydrogels for organoid formation above 1 mg/mL. Image analysis fits seamlessly with other 3D workflows, and we showed compatibility with microarray methods for bulk histologic evaluation **(Supplementary Fig. 8**). This opens the possibility of use in high throughput drug screening techniques.

As with any in vitro model, there are several limitations and questions that deserve further investigation. There are many additional components within the TME, most notably immune cells and fibroblasts that support tumors via paracrine and metabolic factors that were not studied here. However, the modular nature of this approach could accommodate their use in a heterotypic co-culture system. Additionally, MatriSpheres facilitate control over ECM composition and allow the user to study the impact of diverse composition by using other decellularized tissues, which may have a role in cancer cell phenotype switching during migration to other microenvironments during metastasis. Here, we utilized a healthy orthotopic ECM, but uncharacterized factors within tumor ECM may be essential for improved modeling of a specific tumor type^82,83^. Porcine ECM was used in the present study as a consistent, reliable orthotopic source for CRC culture, and though ECM is highly conserved among mammalian species, xenogeneic effects may affect these described interactions. The small ECM quantities used in MatriSphere formation makes human sourced tumor ECM a practical possibility. This finding supports the use of this method for high-throughput drug screening and precision medicine approaches. There are several avenues for further investigation into the nature of these ECM-cell interactions. We observed qualitatively dense ECM assembly within MatriSpheres, however future studies are required to characterize intraspheroidal stiffness. There are several studies that highlight the importance of ECM stiffness and its role in mechanotransduction in cancer cell signaling, which could substantially alter cancer cell response within the TME^84^. Further characterization is necessary to determine the extent of cancer cell ECM deposition and remodeling at the protein level, as transcriptomic analysis suggests several critical upstream regulators that may influence TME dynamics.

## Conclusion

We introduce MatriSpheres as a novel 3D in vitro tumor model, which enables the enrichment of decellularized ECM to mimic the dense stroma observed within solid tumors. It is the first system to our knowledge that promotes cell-mediated intercellular stroma formation from decellularized ECM in vitro that may have implications to tumorigenesis. This modular approach uses a simple to implement protocol that adds TME complexity without disrupting standard spheroid workflows and retains the capacity for high-throughput drug screening. ECM biomaterials initiated unique cancer cell-directed spheroid formation and adhesion dynamics that simulated tumor morphogenesis with cell line-specific ECM spatial organization. SIS ECM induced significant changes in CRC cell transcriptome and secretome phenotypes, which better recapitulated in vivo CRC tumor cell heterogeneity compared to cells alone spheroids. We believe that MatriSpheres offer a tool to augment in vitro 3D disease models and improve preclinical drug evaluation for clinical translation.

## Methods

### SIS ECM Decellularization and Processing

Porcine small intestines were obtained from Tissue Source LLC (Zionsville, IN) and were kept frozen at −30°C until decellularization. Intestines were decellularized as described previously^19,85^. Briefly, small intestines were thawed and rinsed with MilliQ water to remove residual intestinal fluid. Tissues were cut longitudinally to create a sheet and the muscle layers were removed by mechanical delamination. Smaller strips (∼6 inches wide) of scraped SIS were transferred to a flask containing MilliQ water to remove any loose residual tissue. To disinfect tissues, SIS strips were washed in a 0.1% peracetic acid and 4% ethanol solution in MilliQ water on a mechanical shaker at 300 rpm for 2 hours at ambient temperature. SIS was washed by alternating MilliQ and 1X PBS two times for 15 minutes each at 300 rpm. After washing, wet SIS ECM was transferred to 50 mL conical tubes and frozen at −80°C. Decellularized tissues were lyophilized and milled into a fine powder using a cryogenic grinder (SPEX SamplePrep, Cat. No. 6875). SIS ECM powder was enzymatically digested with pepsin from porcine gastric mucosa (Sigma, Cat. No. P6887, 1 mg/mL) and hydrochloric acid (0.01 M) at a stock ECM concentration of 10 mg/mL under constant magnetic stirring (between 700-1500 rpm) for 48 hours at room temperature. Low concentrations of SIS ECM digest were used for all CRC tumor spheroid cultures.

### ECM Characterization

#### DNA Isolation and Quantification

To validate the removal of immunogenic materials within decellularized tissues, we quantified double-stranded DNA (dsDNA) content. First, we isolated and purified total DNA from native and decellularized small intestine using a DNeasy Blood and Tissue Kit (Qiagen, Cat. No. 69504) in accordance with the manufacturer’s instructions. Briefly, 10-25 mg of lyophilized samples were digested using proteinase K solution and lysed overnight in a water bath at 56°C. Tissue lysate was passed through a DNeasy membrane to bind DNA and remove contaminants. Purified DNA was eluted in 200 µL of AE buffer and stored at −80°C until use. A Quant-iT PicoGreen assay was used for a highly-sensitive fluorescence-based readout of dsDNA content. Native and decellularized DNA samples were serially-diluted from 1:200 to 1:1600 and 1:20 to 1:160, respectively. A dsDNA standard was used as a control and provided a reference range from 31.25-2000 ng/mL. 100 µL of diluted samples and standards were plated onto a 96-well plate and 100 µL of Quant-iT reagent was added. The samples were mixed for 5 minutes and the fluorescence intensity was determined at 480/520 nm excitation and emission wavelengths. DNA fragment size was assessed using a PowerSnap gel electrophoresis (Thermo Fisher) system. A 2% agarose E-gel with SYBR Safe was loaded with diluted samples (1:10 Decell or 1:20 Native) and run for 26 minutes before imaging via transillumination.

#### Sircol Assay

ECM biomaterial solutions were prepared from stock concentrations and assayed using the Sircol kit (S1000, Biocolor Life Science Assay Kits, UK). Diluted samples and standards were mixed with 1 mL of saturated Sirius Red solution in picric acid. The dye-binding incubation step (30 minutes with gentle mechanical agitation) was performed at 4°C to prevent gelation of the Matrigel. Samples were spun at 13,000 rpm for 10 minutes. Excess dye and solution were removed and 750 µL ice-cold acid salt were added to each sample. Samples were spun at 13,000 rpm for 10 minutes. This solution was removed and 1,000 µL alkali reagent were added to each sample and the collagen-bound dye pellets were solubilized by vortexing the samples. 200 µL of each sample (triplicate) and standard (duplicate) were transferred to a 96-well plate. The absorbance was measured at 556 nm using a plate reader (SpectraMax i7, Molecular Devices).

### ECM Biomaterial Digestion and LC/MS analysis

#### MC38 Tumor ECM MS prep

Mass spectrometry was used to compare differences in composition between ECM biomaterials and acellular MC38 tumor ECM. Tumors were established in the right and left flanks of 6-10-week-old C57BL/6 mice (Jackson Lab) with bilateral subcutaneous injections of 500,000 MC38 cells per injection. Ethical approval for the animal experiments was provided by the Institutional Animal Care and Use Committee at NCI Frederick (ASP No. 20-063). Tumors were harvested after two weeks (1 cm in diameter) and were frozen until thawed for future use. To prepare MC38 tumor ECM, we diced MC38 tumors into small pieces and processed with the following steps. Briefly, tissues were washed in MilliQ water then rinsed in hypotonic and hypertonic solutions. Trypsin-EDTA treatment, followed by washes in 4% sodium deoxycholate were designed to detach cells then release cytoplasmic nuclear contents. Residual DNA was broken down by a DNase treatment then 3% Triton X-100 was used to clean up any remaining cellular components. Peracetic acid and ethanol were combined to reduce pyrogenic material. The remaining tumor ECM was washed several times in alternating MilliQ water and 1X PBS washes then lyophilized for downstream analysis.

#### Digestion

For decellularized SIS-ECM powdered samples, 0.54-0.83 mg was treated with 300 µL of EasyPep Lysis buffer (Thermo Fisher PN A45735, provided in EasyPep 96-well plat kit A57864) and treated with 50 µL each of reducing solution and alkylation solution provided with the EasyPep kit. The decellularized SIS-ECM samples were sonicated 3×25 sec at 15% amplitude then all samples were heated at 95°C for 10 minutes. SIS-ECM samples were treated with 50 µL of 0.2 µg/µL trypsin/LysC while the Collagen 1 and Matrigel, samples were treated with 10 µL of 0.2 µg/µL trypsin/LysC. Samples were incubated at 37°C overnight for 18 hrs with shaking at 1000 rpm. Added 100 µL of stop solution provided with the EasyPep kit and cleaned using the EasyPep 96-well plate and eluted with 300 µL of the elution buffer, aliquoted into 20 µL and 250 µL and dried. The 20 µL aliquot was used to assess the peptide concentration using the Pierce Quantitative Colorimetric Peptide Assay (Thermo PN 23275).

#### LC/MS analysis of peptides

Each sample was resuspended to a concentration of 1 µg/µL in 0.1% FA and 1 μL was analyzed using a Dionex U3000 RSLC in front of a Orbitrap Eclipse (Thermo) equipped with a FAIMS interface and an EasySpray ion source. Solvent A consisted of 0.1%FA in water and Solvent B consisted of 0.1%FA in 80%ACN. Loading pump consisted of Solvent A and was operated at 7 μL/min for the first 6 minutes of the run then dropped to 2 μL/min when the valve was switched to bring the trap column (Acclaim™ PepMap™ 100 C18 HPLC Column, 3μm, 75μm I.D., 2cm, PN 164535) in-line with the analytical column EasySpray C18 HPLC Column, 2μm, 75μm I.D., 25cm, PN ES902). The gradient pump was operated at a flow rate of 300 nL/min. Each run used a linear LC gradient of 5-7%B for 1min, 7-30%B for 34 min, 30-50%B for 15min, 50-95%B for 4 min, holding at 95%B for 7 min, then re-equilibration of analytical column at 5%B for 17 min. MS acquisition employed the TopSpeed method at three FAIMS compensation voltages (−45, −60, −75) with a 1 second cycle time for each voltage and the following parameters: Spray voltage was 2200V and ion transfer temperature was 300 ⁰C. MS1 scans were acquired in the Orbitrap with resolution of 120,000, AGC of 4e5 ions, and max injection time of 50 ms, mass range of 350-1600 m/z; MS2 scans were acquired in the Orbitrap a with resolution of 15,000, AGC of 5e4, max injection time of 22ms, HCD energy of 30%, isolation width of 1.06 Da, intensity threshold of 2.5e4 and charges 2-5 for MS2 selection. Advanced Peak Determination, Monoisotopic Precursor selection (MIPS), and EASY-IC for internal calibration were enabled and dynamic exclusion was set to a count of 1 for 15 sec.

#### Database search and post-processing analysis

MS files were searched with Proteome Discoverer 2.4 using the Sequest node. Data was searched against the *Mus musculus* (Matrigel), *Bos taurus* (Collagen 1), or *Sus scrofa* (ECM) Uniprot databases. The SIS-ECM, Matrigel, and Collagen I samples were searched using a full tryptic digest. and allowed for 2 max missed cleavages with a minimum peptide length of 6 amino acids and maximum peptide length of 40 amino acids, an MS1 mass tolerance of 10 ppm, MS2 mass tolerance of 0.02 Da, fixed carbamidomethyl (+57.021) on cysteine and variable oxidation on methionine and proline (+15.995 Da). Percolator was used for FDR on all searches except the Collagen I sample in which Fixed PSM was used. NSAF values for each protein within a sample were calculated by taking the spectral counts of the protein and dividing by the number of amino acids to give the SCP/L value, which was then divided by the total PSMs of the sample to give the NSAF.

### CRC Cell culture

MC38 murine cancer cells were provided by the McVicar Lab and CT26 cells were purchased from the American Type Culture Collection (Manassas, VA). HT-29 human cancer cells were obtained from the NCI Division of Cancer Treatment and Diagnosis Tumor Repository (Frederick, MD). Cells were grown in Roswell Park Memorial Institute (RPMI) media supplemented with L-glutamine (Thermo Fisher 11875119), 10% fetal bovine serum (Thermo Fisher 16000044), and 1% penicillin streptomycin (Thermo Fisher 15140122). Cells were maintained at 37°C with 5% CO_2_ humidified air.

### Spheroid Formation and Culture

Single cells in suspension were counted with a hemocytometer and a single cell suspension at 2x the final cell seeding density was prepared in complete RPMI and kept on ice until mixed with the ECM-containing solution. ECM-containing solutions were prepared at 2x the final seeding concentration (16-250 µg/mL) and kept on ice until mixed with cells. The cell and ECM solutions were mixed 1:1 and transferred to a sterile basin for cell seeding. Spheroids were seeded by pipetting 100 µL of the cell and ECM containing solution into the inner 60 wells of a 96-well ultra-low attachment round bottom well plate (Corning 7007). Spheroids were cultured at 37°C with 5% CO_2_ humidified air by replenishing the media every two to three days. At day seven, 100 µL of media was removed from each well and saved for cytokine analysis by storing at −80°C or discarded. Spheroids were collected for RNA-Seq analysis by storing in TRIzol Reagent (Thermo Fisher 15596018) or fixed by adding 100 µL 8% paraformaldehyde (PFA, Electron Microscopy Sciences 15710) to each well for a final concentration of 4% PFA and fixed at 4°C for at least 48 hours. The spheroids were washed 3x with 1x PBS after fixation and kept at 4°C until histological processing.

### Spheroid Diameters Analysis

Spheroid diameters were measured on days 3, 5, and & 7 of the culture using a Celigo Image Cytometer (Nexcelom, Lawrence, MA) using the tumorsphere setting to acquire brightfield images of the samples. The largest object within each well was determined to be the spheroid for quantification and spheroid diameters were analyzed through the instrument’s software. Spheroids were manually traced in ImageJ (NIH, Bethesda, MD) for cases where the analysis parameters could not be adjusted to capture the actual spheroid diameter. The equivalent diameters reported are calculated from the spheroid area and as if the spheroid diameter was an actual circle.

### Spheroid Microarray Embedding & Processing

Cancer cell spheroids were embedded into agarose similarly as described previously^22^ using a 3D printed plastic mold *(see description below)*. Briefly, the wells of the mold were filled with water and all air bubbles were removed. PFA fixed spheroids that were washed 3x for 10 minutes each with 1x PBS were transferred to the microarray mold by pipetting. The spheroids settled to the bottom of the mold by gravity. A molten hot solution of 3% agarose was pipetted into the mold and a histology cassette was placed on top and additional agarose was added to fill the mold. The mold was transferred to an incubator set to 65°C for 5 minutes, then cooled on the bench for an additional 25 minutes. The microarray was removed from the mold and transferred to 30% ethanol for at least 3 hours, followed by 50% and 70% ethanol for 3 hours each before automated processing and paraffin embedding. The automated processing of the microarrays was: 70% ethanol, 80% ethanol, 95% ethanol, 100% ethanol x 3 each step for 3 hours, xylene x 2 each step for 4.5 hours followed by 4x paraffin wax for 3 hours x 2, 2 hours, then 1 hour.

### 3D Printing of Spheroid Microarray Mold

We designed a microarray mold that would allow us to increase the throughput of our histological staining and imaging of MatriSpheres. The mold was sized to fit a standard tissue embedding cassette (Sigma Cat. #Z672122) to avoid shifting during agarose curing. The 3D model of the mold was drafted using SolidWorks 3D CAD software (v2022, Dassault Systèmes, Vélizy-Villacoublay, France). The file was then exported to .STL format, processed in Formlabs PreForm software (v3.27.1, Formlabs Inc., Sommerville, MA, USA), and printed on a Formlabs printer. Parts were fabricated from a biocompatible and autoclavable material, Biomed Clear resin (Formlabs Inc.), using a Formlabs 3B printer (Formlabs Inc.). The parts were post-processed using a Form Wash solvent system (Formlabs Inc.) and the Form Cure UV heated chamber system (Formlabs Inc.).

### LIVE/DEAD Spheroid Staining and Imaging

Spheroids at day 7 of culture were stained with Calcein AM and Ethidium-homodimer (Thermo L3224) in accordance with the manufacturer’s instructions. Stained spheroids were imaged with the ImageXpress under confocal microscopy at 4X magnification with an offset ∼200 µm and z-stack range at depths between 100 and 300 µm. For three-dimensional spheroid imaging, resolution significantly diminishes after 300 µm without optical clearing. However, we were most interested in sampling both the outer edge and core of the spheroid, which we can image at the noted depths. After imaging, brightfield, FITC and TRITC images were analyzed using a custom module editor on MetaXpress software. Briefly, an image mask was determined from brightfield images to identify the spheroid area. The fluorescent channels were used to identify Live (FITC) and Dead (TRITC) cells within the spheroid region.

### CellTiter-Glo 3D Analysis

Cancer spheroids were cultured for seven days and 100 μL of media were removed from each of the wells. The plates were brought to room temperature and 100 μL of room temperature CellTiter-Glo 3D (Promega G9683) were added to each of the wells. The plates were incubated on a shaker at 151 rpm for five minutes, followed by an additional 25 minutes on the bench. Then, 100 μL was transferred from each well into a white opaque 96-well plate and the luminescence was read by a plate reader (Molecular Devices).

### Spheroid Dissociation and Counting

Cancer spheroids were cultured for seven days and transferred to 1.5 mL tubes for each spheroid condition. The spheroids were pelleted in the bottom of the tube by centrifugation at 200xg for five minutes. The media was removed and 150 μL of serum free RPMI containing 0.25 mg/mL Liberase TL (Roche 054010120001) were added to the samples. The samples were incubated in a water bath for two minutes followed by 20 minutes on a shaker at 200 rpm, both at 37°C. The spheroids were then dissociated with pipetting and the enzymatic reaction was quenched by adding 15 μL FBS. Then, 14.2 μL NuncBlue Live reagent and 11.4 μL NuncGreen Dead reagent (Thermo) were added to each of the tubes and incubated at 37°C for 15 minutes. Cell counts from each sample were obtained with a Countess II FL Automated Cell Counter (Thermo).

### Live-Spheroid Formation Acquisition

After cells and ECM had been plated as described above, the plate was transferred to an ImageXpress Micro Confocal System (Molecular Devices, San Jose, CA) with environmental controls set to 37°C and 5% CO_2_. Within 5 minutes of plating, images acquisition began using a 10x objective and were obtained every 15 minutes for the culture period. Imaging was suspended temporarily during the spheroid feeding. We observed differences in the time needed for spheroid formation during incubator conditions compared with spheroids that were formed during the timelapse imaging. It took longer for the spheroids to form in the imaging experiments, and we suspect this may be a result of the movement of the plate during the image acquisition.

### Rheological Characterization

To determine the mechanical properties of the ECM biomaterials, we used a Discovery HR20 rheometer (TA Instruments – Schneider Lab) and a 40 mm stainless steel parallel plate geometry (Cat. No. 511400.945). ECM biomaterials were prepared at various concentrations (62.5, 125, 1000 and 4000, 5000 and 7000 µg/mL) to determine the effects of ECM concentration on gelation compared to RPMI media controls. Briefly, the rheometer temperature was set to 4°C to prevent premature gelation and ∼700 µL of sample was pipetted onto the center of the Peltier plate. The geometry was slowly lowered to establish a uniform spreading and mineral oil was added to the edge of the sample to avoid evaporation during testing. A brief temperature ramp was performed to 37°C to mimic the culture conditions followed by a time sweep.

### Immunohistochemistry

Spheroid sections were deparaffinized in xylenes 3x for 3 minutes each followed by rehydration in 100% ethanol 2x, 95% ethanol, 70% ethanol, and type I water 3x. Next, sections were refixed in 10% neutral buffered formalin (Sigma HT501128-4L) for 15 minutes. Slides were then washed in type I water and TBS-T for 3 minutes each. 10 mM sodium citrate (Sigma) buffer at pH 6 was preheated in a vegetable steamer for 20 minutes to reach 95°C. Slides were heated in the buffer for 20 minutes and then cooled at room temperature for 20 minutes. Following antigen retrieval, endogenous peroxidases were blocked with 3% H_2_O_2_ (Sigma H1009-100ML) for 15 minutes. Slides were washed 2x in TBS-T. Aldehydes were quenched with glycine in TBS-T for 5 minutes at room temperature. Blocking buffer (10% BSA (Sigma A9647-100G), 5% Goat Serum (Vector Laboratories S-1000-20), 0.5% Tween-20 (Sigma) in TBS-T) was incubated for a minimum of 30 minutes on the sections at room temperature before primary antibody incubation. Primary antibodies (Abcam) were incubated at 4°C overnight in a humidity chamber and diluted in blocking buffer at the following concentrations: E-Cadherin (ab76319): 1/10000, N-Cadherin (ab76011): 1/8000, Carbonic Anhydrase IX (CA9, ab243660): 1/2000, and Ki-67 (ab16667): 1/1000. A FITC-conjugated collagen hybridizing peptide (CHP, 3Helix FLU300) was heated on a heating block for 15 minutes at 80°C for 15 minutes and then allowed to cool to room temperature. The CHP was diluted to 1 µM and incubated overnight at 4°C with the last primary antibody in each panel. After primary antibody and CHP incubations, slides were washed 3x in TBS-T for 3 minutes. Rabbit-on-Rodent Secondary HRP antibody (Biocare Medical RMR622H) was added to the slides for 20 minutes at room temperature. Next, slides were washed 4x for 3 minutes each. Opal reagents (Akoya Biosciences) were added to slides: Opal 570 (1:150) for CA9 and E-Cadherin and Opal 650 (1:250) for Ki67 and N-Cadherin. For subsequent stains in a panel, slides were reheated in citrate buffer at 95°C for 20 minutes and allowed to cool 20 minutes before proceeding. Once all antibody stains were performed, 1 µg/mL DAPI was added to slides for 5 minutes at room temperature and then washed 2x with type I water. Coverslips were mounted with Dako Fluorescent Mounting Media (Agilent S302380-2) and allowed to dry. A rabbit IgG isotype (Abcam ab172730) and primary deletes were performed as controls. Slides were imaged with an ImageXpress Micro Confocal Imaging System using a 20X water immersion objective. The images were analyzed in MetaXpress with multi-wavelength cell scoring.

### In Vitro CRC Spheroid Bulk RNA Isolation, Sequencing, and Analysis

After 7 days of culture, CRC spheroids were transferred to 5 mL tubes in batches of 10, the excess media was removed by pipetting, 1 mL of TRIzol (Invitrogen) was added to each sample, and the samples were stored at −80°C until further processing. Samples were thawed on ice and the RNA was extracted using the RNeasy Micro Kit (QIAGEN, Germantown, MD) in accordance with the manufacturer’s instructions. RNA Samples were then pooled and sequenced on a NovaSeq 6000 S1 using Illumina Stranded mRNA Prep and paired-end sequencing. Cutadapt (v1.18) was used to trim reads and samples for adapters and low-quality bases before alignment to the reference genomes (hg38 for human and mm10 for mouse) and annotation of transcripts using STAR (v2.7.0f) in 2-pass mode^86^. Expression quantification was performed using STAR, RSEM (v1.3.1)^87^, and the gene annotations for human (GENCODE_30) and mouse (GENCODE_M21), respectively. Downstream analysis and visualization were performed within the NIH Integrated Analysis Platform (NIDAP) using R programs developed on the Foundry platform (Palantir Technologies). The RSEM gene counts matrix was imported into the NIDAP platform where genes were filtered for low counts (<1 cpm) and normalized by quantile normalization using the limma package^88^. Differentially expressed genes were calculated using limma-Voom and batch removal was performed on mouse samples using ComBat^89^. The fgsea package^90^ was used to perform pre-ranked GSEA.

Datasets were imported to Ingenuity Pathway Analysis (IPA, Qiagen) for calculating pathway activation patterns or z-scores.

### In Vivo MC38 Tumor Establishment and Collection for scRNA Seq

Seven-month-old C57BL/6 mice were obtained from Charles River Laboratories and injected with 1x 10^6^ cells in 100 µL PBS per mouse. Ethical approval for the animal experiments was provided by the Institutional Animal Care and Use Committee at NCI Frederick (ASP No. 23-078). Tumors were carefully excised from the mice when the size reached up to 250-500 mm^3^, which was usually after 10-14 days. The tumor tissues were washed with cold sterile PBS. Fat and necrotic areas and blood vessels were removed from the samples. The MC38 tumor single cell suspension preparation began by preparing the enzyme mix in accordance with the protocol of the Miltenyi Tumor Dissociation Kit (mouse) and placing on ice. One mL of ready-to-use enzyme mixture was then added to sterile petri dishes for each sample and placed on ice. Subsequently, the tumor tissue was placed in the dish with the enzyme mix and cut into small pieces of 2– 4 mm^3^ using sterile blades. Tissue pieces were then transferred into the gentleMACS C Tube containing the remaining enzyme mix. The C Tube were tightly closed and positioned upside down onto the sleeve of the gentleMACS Dissociator. The program was run in accordance with the instruction of the Miltenyi Tumor Dissociation Kit (mouse). Following completion of the program, cells were centrifuged at 300g x 10 min followed by one wash using cold PBS at 300g x 10 min. Red blood cells were removed using anti-Ter-119 microbeads (Miltenyi) according to the manufacturer’s instruction. Cells were counted and spun down at 300g x 10 min. Dead cells were removed using dead cell removal kit (Miltenyi) in accordance with the protocol. Cells were counted and the viability was determined, ensuring that the cell viability was higher than 90%. Finally, cells were cryopreserved in a solution of 90% FBS + 10% DMSO until further processing.

### MC38 Tumor RNA Isolation, scRNA Sequencing, and Analysis

The frozen MC38 single cell suspension was thawed in 37°C water bath and promptly transferred into 5 mL of DMEM containing 10% FBS. Subsequently, the cells were centrifuged at 350 x 5 min. The cell pellet was then washed three times with PBS + 0.04% BSA at 350 x 5 min before further processing. Dead cells need to be removed if the cell viability is lower than 70%. Aiming to capture 10,000 cells, 16,000 cells per sample were used for generating scRNAseq libraries following the guidance of 10x Genomics Chromium Next GEM Single Cell 3’ Gene Expression version 3.1. Briefly, RT reagents, cell suspension, along with the gel beads, were loaded onto Chromium NextGEM chip G (separate channels for different samples). Gel Beads-in-emulsion (GEMs) generated using the 10X Genomics Chromium Controller were then transferred and incubated in a thermal cycler for GEM-RT incubation. The barcoded cDNAs generated were then purified, amplified, cleaned up. Subsequently, Bioanalyzer (Agilent Technologies) was run to determine cDNA concentration. The libraries were prepared from the cDNA according to the guidance of 3’ gene expression dual index library construction. The libraries were sequenced on NovaSeq_SP system using read lengths: 28bp (Read 1), 10bp (Index i7), 10bp (Index i5) and 90bp (Read 2) (Illumina).

### In Vivo MC38 scRNA-seq Processing and Analysis

Single-cell RNA sequencing data was processed using the Seurat package (version 4.4.0) in R (version 4.2.2)^91^. Raw sequencing data was first preprocessed to remove low-quality cells and mitochondrial genes. Next, the data was normalized and scaled using the NormalizeData and the ScaleData functions, respectively. The multiple scRNA-seq datasets were integrated using the harmony package (version 1.1.0)^92^. Then, the cells were grouped into clusters using PCA for dimensionality reduction and visualized using t-SNE projection. Cell types were assigned using scType method^93^ in combination with manual annotation. Pseudo-bulk data for each sample was extracted using AverageExpression function from Seurat. T-SNE plots and violin plots reflecting individual gene expression were generated using Loupe Browser software (10x Genomics, version 6.4.1). The dotplots reflecting expression of each gene sets in the different cell types were made using custom functions for Z-score calculation and plotting. The heatmaps were done using the ComplexHeatmap package (version 2.14.0)^94,95^. Scissor package (version 2.0.0) was used to identify the most highly phenotype-associated cell subpopulations^40^. Scissor uses the bulk assays corresponding to each phenotype to annotate the phenotype-associated cells from scRNA-seq data. SIS ECM MatriSpheres and traditional MC38 spheroids RNA-seq data was used to identify to which phenotype each cell from MC38 scRNA-seq data was associated.

### Cytokine Arrays

To assess the effects of SIS ECM on CRC cell cytokine secretion, we used both mouse (111 analytes) and human (105 analytes) proteome profiler XL kits (R&D Systems, Cat No. ARY028 and ARY022B). We screened the cell culture supernatant from Day 7 spheroid cultures with and without SIS ECM. Briefly, 100 µL of media was collected from each well (n = 30) and batched by row (10 wells/row) for a total of 1 mL (n = 3 replicates). Media was frozen at −80°C and thawed on ice before use. For each sample array, 400 µL of cell culture supernatant were used to determine the presence of cytokines. Bound cytokines were detected using streptavidin-HRP conjugated antibodies and chemi reagent, which produced a chemiluminescent signal, captured on an Amersham Imager 680 digital imaging system (GE Life Sciences). Integrated intensity was determined using an ImageJ plugin (Protein Array Analyzer) and exported to Excel for data analysis.

### Upstream Regulators and Target Genes Analysis

Ingenuity Pathway Analysis was used to identify conserved upstream regulators and target genes within CRC MatriSpheres compared to cells alone spheroids. IPA software predicted 6,670 combined upstream regulators associated with the addition of SIS ECM across all three cell lines. We narrowed this list of upstream regulators by 91.3% using exclusion criteria based upon |z-score| ≥ 2 and p-value < 0.001. From this list of 580 significant upstream regulators, we found 10 upstream regulators shared between each of the three CRC cells lines. We then combined the list of target genes associated with each of the upstream regulators and found 35 common target genes. The selected regulators and target genes were put back into IPA to create an interaction plot **(Supplemental Fig. 6g**). This was subsequently narrowed to only direct and shared interactions between upstream regulators and target genes **(Supplemental Fig. 6h,i**). The top 5 upstream regulators were then superimposed onto the Tumor Microenvironment Pathway in IPA to determine their role in TME signaling **(Supplementary Fig. 6j,k**).

### Statistical Analysis

Statistical Analyses were performed in GraphPad Prism (v9). For data with a normal distribution and three or more groups, statistics were calculated by a one-way ANOVA followed by a Tukey’s or Sidak’s multiple comparisons test. A two-way ANOVA followed by a Dunnett’s or Sidak’s multiple comparisons test were used to calculate statistics for data with two variables and three or more groups. P values are indicated in the figure legends.

## Author Contributions

M.J.B., E.A.B., and M.T.W. developed the concept for MatriSpheres and prepared the manuscript. M.J.B. and E.A.B. established the methods for MatriSphere formation and characterized the model. M.J.B., E.A.B. and M.S.T. decellularized and processed SIS ECM. R.J.H. and T.A. performed mass spectrometry on ECM biomaterials and M.J.B. analyzed the data. E.A.B. carried out Sircol assays of ECM biomaterials. M.J.B., M.G.C., and T.J.P. designed and 3D printed the spheroid microarray mold. M.J.B. and E.A.B. performed spheroid cultures, live-dead assays, and spheroid dissociation. E.A.B. imaged and quantified spheroid diameters and ran CellTiterGlo 3D assays. M.S.T. and E.A.B. stained, imaged, and analyzed spheroid microarray slides. M.J.B. and E.A.B. isolated RNA and T.J.M. assisted with Bulk RNA Sequencing workflows. M.J.B. analyzed the Bulk and scRNA Seq datasets using GSEA, IPA and Loupe Browser. M.J.B. performed and analyzed cytokine arrays. L.Y., B.S.C. generated MC38 tumors for scRNA Seq and M.G. performed scRNA Seq analysis. All authors reviewed and edited the final manuscript.

## Acknowledgements

This research was supported by the Intramural Research Program of the National Institutes of Health (NIH), National Cancer Institute (NCI), Center for Cancer Research (CCR) and Cancer Innovation Laboratory (CIL). We would like to thank our CCR colleagues from the labs of Dr. Daniel McVicar, Dr. Howard Young, Dr. Scott Durum, Dr. Joost Oppenheim, Dr. Stephen Anderson and Dr. Joel Schneider for their sharing of knowledge and resources. We appreciate the support given by the following core facilities: (1) Optical Microscopy and Analysis Laboratory – Dr. Stephen Lockett and Dr. William Heinz (2) CCR Sequencing Facility – Dr. Maggie Cam and the (3) Protein Characterization Laboratory. Thanks to Dr. Kaitlin Fogg for her insights on potential mechanisms of MatriSphere assembly. Illustrations were created with BioRender.com.

## Data and code availability

Data and code will be made available upon reasonable request. All proteomic data has been uploaded to public databases MassIVE server. Bulk and scRNA sequencing datasets have been deposited in GEO. Code used to analyze quantified mRNA expression is available on GitHub.

## References

1 de Visser, K. E. & Joyce, J. A. The evolving tumor microenvironment: From cancer initiation to metastatic outgrowth. Cancer Cell 41, 374–403 (2023). 10.1016/j.ccell.2023.02.016

2 Brauchle, E. et al. Biomechanical and biomolecular characterization of extracellular matrix structures in human colon carcinomas. Matrix Biol 68-69, 180-193 (2018). 10.1016/j.matbio.2018.03.016

3 Devarasetty, M. et al. Simulating the human colorectal cancer microenvironment in 3D tumor-stroma co-cultures in vitro and in vivo. Sci Rep 10, 9832 (2020). 10.1038/s41598-020-66785-1

4 Lu, P., Weaver, V. M. & Werb, Z. The extracellular matrix: A dynamic niche in cancer progression. Journal of Cell Biology 196, 395–406 (2012). 10.1083/jcb.201102147

5 Cox, T. R. The matrix in cancer. Nat Rev Cancer 21, 217–238 (2021). 10.1038/s41568-020-00329-7

6 Nebuloni, M. et al. Insight On Colorectal Carcinoma Infiltration by Studying Perilesional Extracellular Matrix. Scientific Reports 6, 22522 (2016). 10.1038/srep22522

7 Karlsson, S. & Nystrom, H. The extracellular matrix in colorectal cancer and its metastatic settling - Alterations and biological implications. Crit Rev Oncol Hematol 175, 103712 (2022). 10.1016/j.critrevonc.2022.103712

8 Raghavan, S. et al. Comparative analysis of tumor spheroid generation techniques for differential in vitro drug toxicity. Oncotarget 7 (2016).

9 Giobbe, G. G. et al. Extracellular matrix hydrogel derived from decellularized tissues enables endodermal organoid culture. Nature Communications 10 (2019). 10.1038/s41467-019-13605-4

10 Fujii, M. et al. A Colorectal Tumor Organoid Library Demonstrates Progressive Loss of Niche Factor Requirements during Tumorigenesis. Cell Stem Cell 18, 827–838 (2016). 10.1016/j.stem.2016.04.003

11 LeSavage, B. L., Suhar, R. A., Broguiere, N., Lutolf, M. P. & Heilshorn, S. C. Next-generation cancer organoids. Nat Mater 21, 143–159 (2022). 10.1038/s41563-021-01057-5

12 Ferreira, L. P., Gaspar, V. M., Mendes, L., Duarte, I. F. & Mano, J. F. Organotypic 3D decellularized matrix tumor spheroids for high-throughput drug screening. Biomaterials 275, 120983 (2021). 10.1016/j.biomaterials.2021.120983

13 van Pelt, G. W. et al. The tumour–stroma ratio in colon cancer: the biological role and its prognostic impact. Histopathology 73, 197–206 (2018). 10.1111/his.13489

14 Sadtler, K. et al. Proteomic composition and immunomodulatory properties of urinary bladder matrix scaffolds in homeostasis and injury. Semin Immunol 29, 14–23 (2017). 10.1016/j.smim.2017.05.002

15 Gonzalez-Fernandez, T., Tenorio, A. J., Saiz, A. M., Jr. & Leach, J. K. Engineered Cell-Secreted Extracellular Matrix Modulates Cell Spheroid Mechanosensing and Amplifies Their Response to Inductive Cues for the Formation of Mineralized Tissues. Adv Healthc Mater 11, e2102337 (2022). 10.1002/adhm.202102337

16 Beachley, V. Z. et al. Tissue matrix arrays for high-throughput screening and systems analysis of cell function. Nat Methods 12, 1197–1204 (2015). 10.1038/nmeth.3619

17 Klemm, F. & Joyce, J. A. Microenvironmental regulation of therapeutic response in cancer. Trends in Cell Biology 25, 198–213 (2015). 10.1016/j.tcb.2014.11.006

18 Huleihel, L. et al. Matrix-bound nanovesicles within ECM bioscaffolds. Science Advances 2 (2016). 10.1126/sciadv.1600502

19 Badylak, S. F. et al. The use of xenogeneic small intestinal submucosa as a biomaterial for Achilles tendon repair in a dog model. J Biomed Mater Res 29, 977–985 (1995). 10.1002/jbm.820290809

20 Naba, A. et al. The matrisome: in silico definition and in vivo characterization by proteomics of normal and tumor extracellular matrices. Mol Cell Proteomics 11, M111 014647 (2012). 10.1074/mcp.M111.014647

21 Bolean, M. et al. Matrix vesicle biomimetics harboring Annexin A5 and alkaline phosphatase bind to the native collagen matrix produced by mineralizing vascular smooth muscle cells. Biochim Biophys Acta Gen Subj 1864, 129629 (2020). 10.1016/j.bbagen.2020.129629

22 Gabriel, J., Brennan, D., Elisseeff, J. H. & Beachley, V. Microarray Embedding/Sectioning for Parallel Analysis of 3D Cell Spheroids. Scientific Reports 9 (2019). 10.1038/s41598-019-52007-w

23 Ueno, H. et al. Prognostic value of desmoplastic reaction characterisation in stage II colon cancer: prospective validation in a Phase 3 study (SACURA Trial). Br J Cancer 124, 1088–1097 (2021). 10.1038/s41416-020-01222-8

24 Freytes, D. O., Martin, J., Velankar, S. S., Lee, A. S. & Badylak, S. F. Preparation and rheological characterization of a gel form of the porcine urinary bladder matrix. Biomaterials 29, 1630–1637 (2008). 10.1016/j.biomaterials.2007.12.014

25 Shen, Y. et al. Reduction of Liver Metastasis Stiffness Improves Response to Bevacizumab in Metastatic Colorectal Cancer. Cancer Cell 37, 800–817 e807 (2020). 10.1016/j.ccell.2020.05.005

26 Baker, A. M., Bird, D., Lang, G., Cox, T. R. & Erler, J. T. Lysyl oxidase enzymatic function increases stiffness to drive colorectal cancer progression through FAK. Oncogene 32, 1863–1868 (2013). 10.1038/onc.2012.202

27 Rahbari, N. N. et al. Anti-VEGF therapy induces ECM remodeling and mechanical barriers to therapy in colorectal cancer liver metastases. Sci. Transl. Med. 8 (2016). 10.1126/scitranslmed.aaf5219

28 Horn, L. A. et al. Remodeling the tumor microenvironment via blockade of LAIR-1 and TGF-β signaling enables PD-L1–mediated tumor eradication. The Journal of Clinical Investigation 132 (2022). 10.1172/JCI155148

29 Valkenburg, K. C., de Groot, A. E. & Pienta, K. J. Targeting the tumour stroma to improve cancer therapy. Nat Rev Clin Oncol 15, 366–381 (2018). 10.1038/s41571-018-0007-1

30 Hynes, R. O. The extracellular matrix: not just pretty fibrils. Science 326, 1216–1219 (2009). 10.1126/science.1176009

31 van Kuijk, S. J. et al. Prognostic Significance of Carbonic Anhydrase IX Expression in Cancer Patients: A Meta-Analysis. Front Oncol 6, 69 (2016). 10.3389/fonc.2016.00069

32 Wang, Y., Liu, Y., Huang, Z., Chen, X. & Zhang, B. The roles of osteoprotegerin in cancer, far beyond a bone player. Cell Death Discovery 8 (2022). 10.1038/s41420-022-01042-0

33 Liu, S., Sima, X., Liu, X. & Chen, H. Zinc Finger Proteins: Functions and Mechanisms in Colon Cancer. Cancers 14, 5242 (2022). 10.3390/cancers14215242

34 Butler, K. & Banday, A. R. APOBEC3-mediated mutagenesis in cancer: causes, clinical significance and therapeutic potential. Journal of Hematology & Oncology 16 (2023). 10.1186/s13045-023-01425-5

35 Wang, S., Pang, L., Liu, Z. & Meng, X. SERPINE1 associated with remodeling of the tumor microenvironment in colon cancer progression: a novel therapeutic target. BMC Cancer 21 (2021). 10.1186/s12885-021-08536-7

36 Tanagala, K. K. K. et al. SP140 inhibits STAT1 signaling, induces IFN-γ in tumor-associated macrophages, and is a predictive biomarker of immunotherapy response. Journal for ImmunoTherapy of Cancer 10, e005088 (2022). 10.1136/jitc-2022-005088

37 Sharma, B. R. et al. The Transcription Factor IRF9 Promotes Colorectal Cancer via Modulating the IL-6/STAT3 Signaling Axis. Cancers 14, 919 (2022). 10.3390/cancers14040919

38 Chu, P.-Y. et al. IFITM3 promotes malignant progression, cancer stemness and chemoresistance of gastric cancer by targeting MET/AKT/FOXO3/c-MYC axis. Cell & Bioscience 12 (2022). 10.1186/s13578-022-00858-8

39 Yu, F. et al. IFITM1 promotes the metastasis of human colorectal cancer via CAV-1. Cancer Letters 368, 135–143 (2015). 10.1016/j.canlet.2015.07.034

40 Sun, D. et al. Identifying phenotype-associated subpopulations by integrating bulk and single-cell sequencing data. Nat Biotechnol 40, 527–538 (2022). 10.1038/s41587-021-01091-3

41 Amatori, S., Tavolaro, S., Gambardella, S. & Fanelli, M. The dark side of histones: genomic organization and role of oncohistones in cancer. Clin Epigenetics 13, 71 (2021). 10.1186/s13148-021-01057-x

42 Jung, G., Hernández-Illán, E., Moreira, L., Balaguer, F. & Goel, A. Epigenetics of colorectal cancer: biomarker and therapeutic potential. Nature Reviews Gastroenterology & Hepatology 17, 111–130 (2020). 10.1038/s41575-019-0230-y

43 Ren, F. et al. MYBL2 is an independent prognostic marker that has tumor-promoting functions in colorectal cancer. Am J Cancer Res 5, 1542–1552 (2015).

44 Giacomini, I. et al. Cholesterol Metabolic Reprogramming in Cancer and Its Pharmacological Modulation as Therapeutic Strategy. Front Oncol 11, 682911 (2021). 10.3389/fonc.2021.682911

45 Xu, L. et al. Quantitative proteomics reveals that distant recurrence-associated protein R-Ras and Transgelin predict post-surgical survival in patients with Stage III colorectal cancer. Oncotarget 7, 43868–43893 (2016). 10.18632/oncotarget.9701

46 Li, X. Y. et al. OAS3 is a Co-Immune Biomarker Associated With Tumour Microenvironment, Disease Staging, Prognosis, and Treatment Response in Multiple Cancer Types. Front Cell Dev Biol 10, 815480 (2022). 10.3389/fcell.2022.815480

47 Sundram, V. et al. Protein Kinase D1 attenuates tumorigenesis in colon cancer by modulating β-catenin/T cell factor activity. Oncotarget 5, 6867–6884 (2014). 10.18632/oncotarget.2277

48 Chen, Y.-J. et al. Interferon regulatory factor family influences tumor immunity and prognosis of patients with colorectal cancer. Journal of Translational Medicine 19 (2021). 10.1186/s12967-021-03054-3

49 Dowling, C. M. et al. Expression of protein kinase C gamma promotes cell migration in colon cancer. Oncotarget 8, 72096–72107 (2017). 10.18632/oncotarget.18916

50 Cai, X. et al. Down-regulation of FN1 inhibits colorectal carcinogenesis by suppressing proliferation, migration, and invasion. Journal of Cellular Biochemistry 119, 4717–4728 (2018). 10.1002/jcb.26651

51 Yi, W., Xiao, E., Ding, R., Luo, P. & Yang, Y. High expression of fibronectin is associated with poor prognosis, cell proliferation and malignancy via the NF-κB/p53-apoptosis signaling pathway in colorectal cancer. Oncology Reports 36, 3145–3153 (2016). 10.3892/or.2016.5177

52 Łukaszewicz-Zając, M., Pączek, S., Mroczko, P. & Kulczyńska-Przybik, A. The Significance of CXCL1 and CXCL8 as Well as Their Specific Receptors in Colorectal Cancer. Cancer Management and Research Volume 12, 8435–8443 (2020). 10.2147/cmar.s267176

53 Huang, B., Song, B.-L. & Xu, C. Cholesterol metabolism in cancer: mechanisms and therapeutic opportunities. Nature Metabolism 2, 132–141 (2020). 10.1038/s42255-020-0174-0

54 Kim, G.-D. et al. CITED2 Restrains Proinflammatory Macrophage Activation and Response. Molecular and Cellular Biology 38, e00452–00417 (2018). 10.1128/MCB.00452-17

55 Chou, Y. T. et al. CITED2 functions as a molecular switch of cytokine-induced proliferation and quiescence. Cell Death & Differentiation 19, 2015–2028 (2012). 10.1038/cdd.2012.91

56 Bai, L. & Merchant, J. L. A role for CITED2, a CBP/p300 interacting protein, in colon cancer cell invasion. FEBS Letters 581, 5904–5910 (2007). 10.1016/j.febslet.2007.11.072

57 Neal, J. T. et al. Organoid Modeling of the Tumor Immune Microenvironment. Cell 175, 1972–1988.e1916 (2018). 10.1016/j.cell.2018.11.021

58 Tuveson, D. & Clevers, H. Cancer modeling meets human organoid technology. Science 364, 952–955 (2019). doi:10.1126/science.aaw6985

59 Veninga, V. & Voest, E. E. Tumor organoids: Opportunities and challenges to guide precision medicine. Cancer Cell 39, 1190–1201 (2021). 10.1016/j.ccell.2021.07.020

60 Cramer, M. C. & Badylak, S. F. Extracellular Matrix-Based Biomaterials and Their Influence Upon Cell Behavior. Annals of Biomedical Engineering 48, 2132–2153 (2020). 10.1007/s10439-019-02408-9

61 Spang, M. T. & Christman, K. L. Extracellular matrix hydrogel therapies: In vivo applications and development. Acta Biomaterialia 68, 1–14 (2018). 10.1016/j.actbio.2017.12.019

62 Hussey, G. S., Dziki, J. L. & Badylak, S. F. Extracellular matrix-based materials for regenerative medicine. Nature Reviews Materials 3, 159–173 (2018). 10.1038/s41578-018-0023-x

63 Kim, S. et al. Tissue extracellular matrix hydrogels as alternatives to Matrigel for culturing gastrointestinal organoids. Nature Communications 13 (2022). 10.1038/s41467-022-29279-4

64 Yamauchi, M., Barker, T. H., Gibbons, D. L. & Kurie, J. M. The fibrotic tumor stroma. The Journal of Clinical Investigation 128, 16–25 (2018). 10.1172/JCI93554

65. Sun, B. The mechanics of fibrillar collagen extracellular matrix. Cell Reports Physical Science 2 (2021).

66 Chen, Y. et al. Oncogenic collagen I homotrimers from cancer cells bind to a3b1 integrin and impact tumor microbiome and immunity to promote pancreatic cancer. Cancer Cell 40, 818–834.e819 (2022). 10.1016/j.ccell.2022.06.011

67 Gavish, A. et al. Hallmarks of transcriptional intratumour heterogeneity across a thousand tumours. Nature 618, 598–606 (2023). 10.1038/s41586-023-06130-4

68 Bejarano, L., Jordāo, M. J. C. & Joyce, J. A. Therapeutic Targeting of the Tumor Microenvironment. Cancer Discovery 11, 933–959 (2021). 10.1158/2159-8290.Cd-20-1808

69 O’Dwyer, P. J. et al. The NCI-MATCH trial: lessons for precision oncology. Nature Medicine 29, 1349–1357 (2023). 10.1038/s41591-023-02379-4

70 Bissell, M. J., Hall, H. G. & Parry, G. How does the extracellular matrix direct gene expression? Journal of Theoretical Biology 99, 31–68 (1982). 10.1016/0022-5193(82)90388-5

71 Winkler, J., Abisoye-Ogunniyan, A., Metcalf, K. J. & Werb, Z. Concepts of extracellular matrix remodelling in tumour progression and metastasis. Nature Communications 11, 5120 (2020). 10.1038/s41467-020-18794-x

72 Gocher, A. M., Workman, C. J. & Vignali, D. A. A. Interferon-γ: teammate or opponent in the tumour microenvironment? Nature Reviews Immunology 22, 158–172 (2022). 10.1038/s41577-021-00566-3

73 Schmitt, M. & Greten, F. R. The inflammatory pathogenesis of colorectal cancer. Nature Reviews Immunology 21, 653–667 (2021). 10.1038/s41577-021-00534-x

74 Xiao, M. et al. Functional significance of cholesterol metabolism in cancer: from threat to treatment. Experimental & Molecular Medicine 55, 1982–1995 (2023). 10.1038/s12276-023-01079-w

75 Xiao, Z., Dai, Z. & Locasale, J. W. Metabolic landscape of the tumor microenvironment at single cell resolution. Nature Communications 10, 3763 (2019). 10.1038/s41467-019-11738-0

76 Demicco, M., Liu, X.-Z., Leithner, K. & Fendt, S.-M. Metabolic heterogeneity in cancer. Nature Metabolism 6, 18–38 (2024). 10.1038/s42255-023-00963-z

77 Yuan, S., Almagro, J. & Fuchs, E. Beyond genetics: driving cancer with the tumour microenvironment behind the wheel. Nature Reviews Cancer (2024). 10.1038/s41568-023-00660-9

78 Guo, J. et al. Cell spheroid creation by transcytotic intercellular gelation. Nature Nanotechnology 18, 1094–1104 (2023). 10.1038/s41565-023-01401-7

79 Wang, H., Feng, Z. & Xu, B. Intercellular Instructed-Assembly Mimics Protein Dynamics To Induce Cell Spheroids. Journal of the American Chemical Society 141, 7271–7274 (2019). 10.1021/jacs.9b03346

80 Stevens, A. J. et al. Programming multicellular assembly with synthetic cell adhesion molecules. Nature 614, 144–152 (2023). 10.1038/s41586-022-05622-z

81 Foty, R. A. & Steinberg, M. S. Differential adhesion in model systems. WIREs Developmental Biology 2, 631–645 (2013). 10.1002/wdev.104

82 Schneider, G., Schmidt-Supprian, M., Rad, R. & Saur, D. Tissue-specific tumorigenesis: context matters. Nature Reviews Cancer 17, 239–253 (2017). 10.1038/nrc.2017.5

83 Deasy, S. K. & Erez, N. A glitch in the matrix: organ-specific matrisomes in metastatic niches. Trends in Cell Biology 32, 110–123 (2022). 10.1016/j.tcb.2021.08.001

84 Ishihara, S. & Haga, H. Matrix Stiffness Contributes to Cancer Progression by Regulating Transcription Factors. Cancers 14, 1049 (2022).

85 Hodde, J. P., Record, R. D., Tullius, R. S. & Badylak, S. F. Retention of endothelial cell adherence to porcine-derived extracellular matrix after disinfection and sterilization. Tissue Eng 8, 225–234 (2002). 10.1089/107632702753724996

86 Dobin, A. et al. STAR: ultrafast universal RNA-seq aligner. Bioinformatics 29, 15–21 (2013).

87 Li, B. & Dewey, C. N. RSEM: accurate transcript quantification from RNA-Seq data with or without a reference genome. BMC bioinformatics 12, 1–16 (2011).

88 Ritchie, M. E. et al. limma powers differential expression analyses for RNA-sequencing and microarray studies. Nucleic acids research 43, e47–e47 (2015).

89 Leek, J. T. et al. sva: Surrogate variable analysis. R package version 3, 882–883 (2019).

90 Korotkevich, G. et al. Fast gene set enrichment analysis. bioRxiv, 060012 (2021). 10.1101/060012

91 R: A Language and Environment for Statistical Computing (R Foundation for Statistical Computing, Vienna, Austria, 2022).

92 Korsunsky, I. et al. Fast, sensitive and accurate integration of single-cell data with Harmony. Nature Methods 16, 1289–1296 (2019). 10.1038/s41592-019-0619-0

93 Ianevski, A., Giri, A. K. & Aittokallio, T. Fully-automated and ultra-fast cell-type identification using specific marker combinations from single-cell transcriptomic data. Nature Communications 13, 1246 (2022). 10.1038/s41467-022-28803-w

94 Gu, Z., Eils, R. & Schlesner, M. Complex heatmaps reveal patterns and correlations in multidimensional genomic data. Bioinformatics 32, 2847–2849 (2016). 10.1093/bioinformatics/btw313

95 Gu, Z. Complex heatmap visualization. iMeta 1, e43 (2022). 10.1002/imt2.43

