## Supplemental Materials for "Engineering Tumor Stroma Morphogenesis Using Dynamic Cell-Matrix Spheroid Assembly"

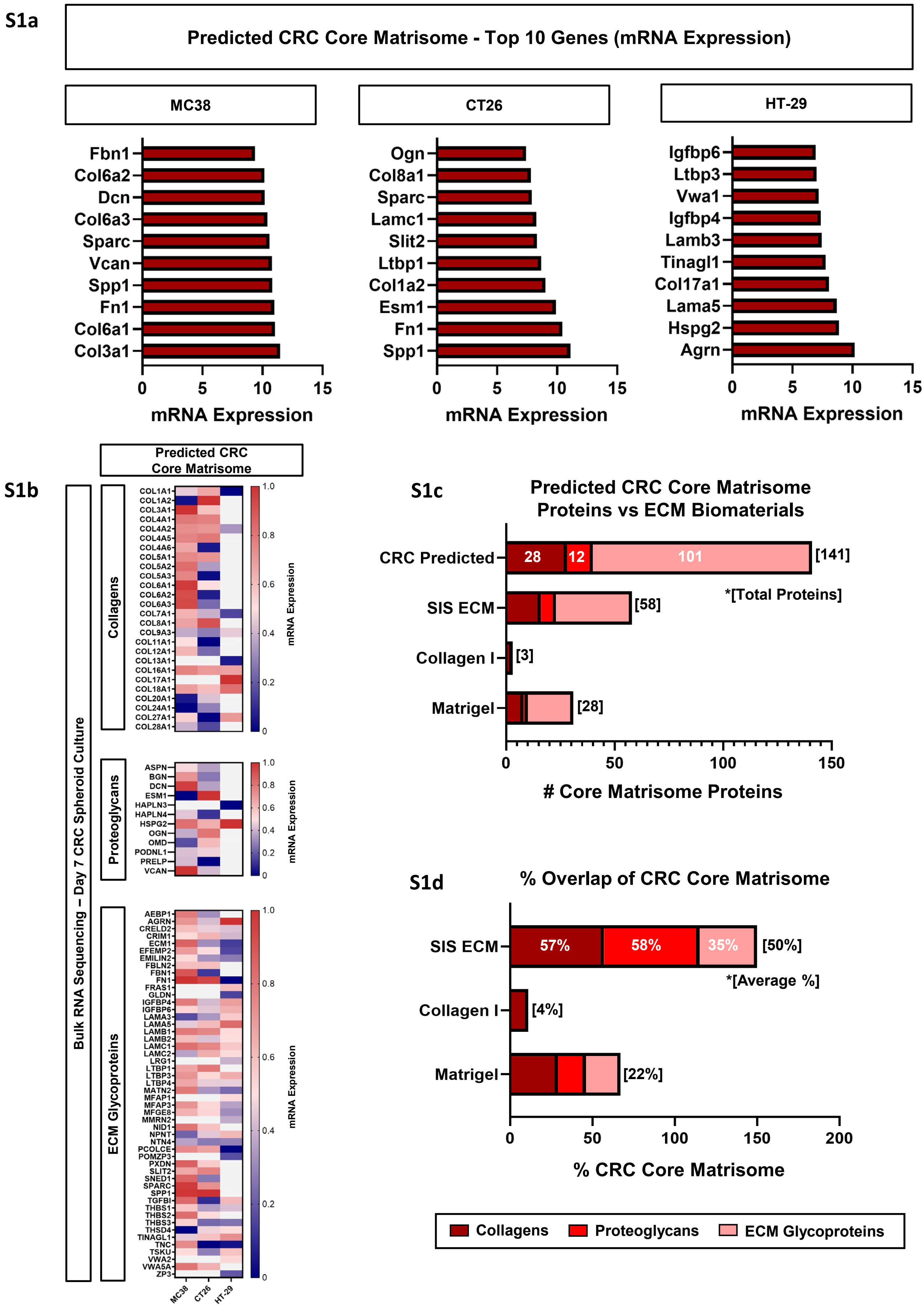

S1a
Predicted CRC Core Matrisome - Top 10 Genes (mRNA Expression)
MC38
Fbn1
Col6a2
Dcn
Col6a3
Sparc
Vcan
Spp1
Fn1
Col6a1
Col3a1
0
5
10
15
mRNA Expression
CT26
Ogn
Col8a1
Sparc
Lamc1
Slit2
Ltbp1
Col1a2
Esm1
Fn1
Spp1
0
5
10
15
mRNA Expression
HT-29
Igfbp6
Ltbp3
Vwa1
Igfbp4
Lamb3
Tinagl1
Col17a1
Lama5
Hspg2
Agrn
0
5
10
15
mRNA Expression
S1b
Predicted CRC Core Matrisome
Bulk RNA Sequencing – Day 7 CRC Spheroid Culture
Collagens
COL1A1
COL1A2
COL3A1
COL4A1
COL4A2
COL4A5
COL4A6
COL5A1
COL5A2
COL5A3
COL6A1
COL6A2
COL6A3
COL7A1
COL8A1
COL9A3
COL11A1
COL12A1
COL13A1
COL16A1
COL17A1
COL18A1
COL20A1
COL24A1
COL27A1
COL28A1
1.0
0.8
0.6
0.4
0.2
0
mRNA Expression
Proteoglycans
ASPN
BGN
DCN
ESM1
HAPLN3
HAPLN4
HSPG2
OGN
OMD
PODNL1
PRELP
VCAN
1.0
0.8
0.6
0.4
0.2
0
mRNA Expression
ECM Glycoproteins
AEBP1
AGRN
CRELD2
CRIM1
ECM1
EFEMP2
EMILIN2
FBLN2
FBN1
FN1
FRAS1
GLDN
IGFBP4
IGFBP6
LAMA3
LAMA5
LAMB1
LAMB2
LAMC1
LAMC2
LRG1
LTBP1
LTBP3
LTBP4
MATN2
MFAP1
MFAP3
MFGE8
MMRN2
NID1
NPNT
NTN4
PCOLCE
POMZP3
PXDN
SLIT2
SNED1
SPARC
SPP1
TGFBI
THBS1
THBS2
THBS3
THSD4
TINAGL1
TNC
TSKU
VWA2
VWA5A
ZP3
1.0
0.8
0.6
0.4
0.2
0
mRNA Expression
MC38
CT26
HT-29
S1c
Predicted CRC Core Matrisome Proteins vs ECM Biomaterials
CRC Predicted
28
12
101
[141]
*[Total Proteins]
SIS ECM
[58]
Collagen I
[3]
Matrigel
[28]
0
50
100
150
### Core Matrisome Proteins
S1d
% Overlap of CRC Core Matrisome
SIS ECM
57%
58%
35%
[50%]
*[Average %]
Collagen I
[4%]
Matrigel
[22%]
0
50
100
150
200
% CRC Core Matrisome
Collagens
Proteoglycans
ECM Glycoproteins

S1e

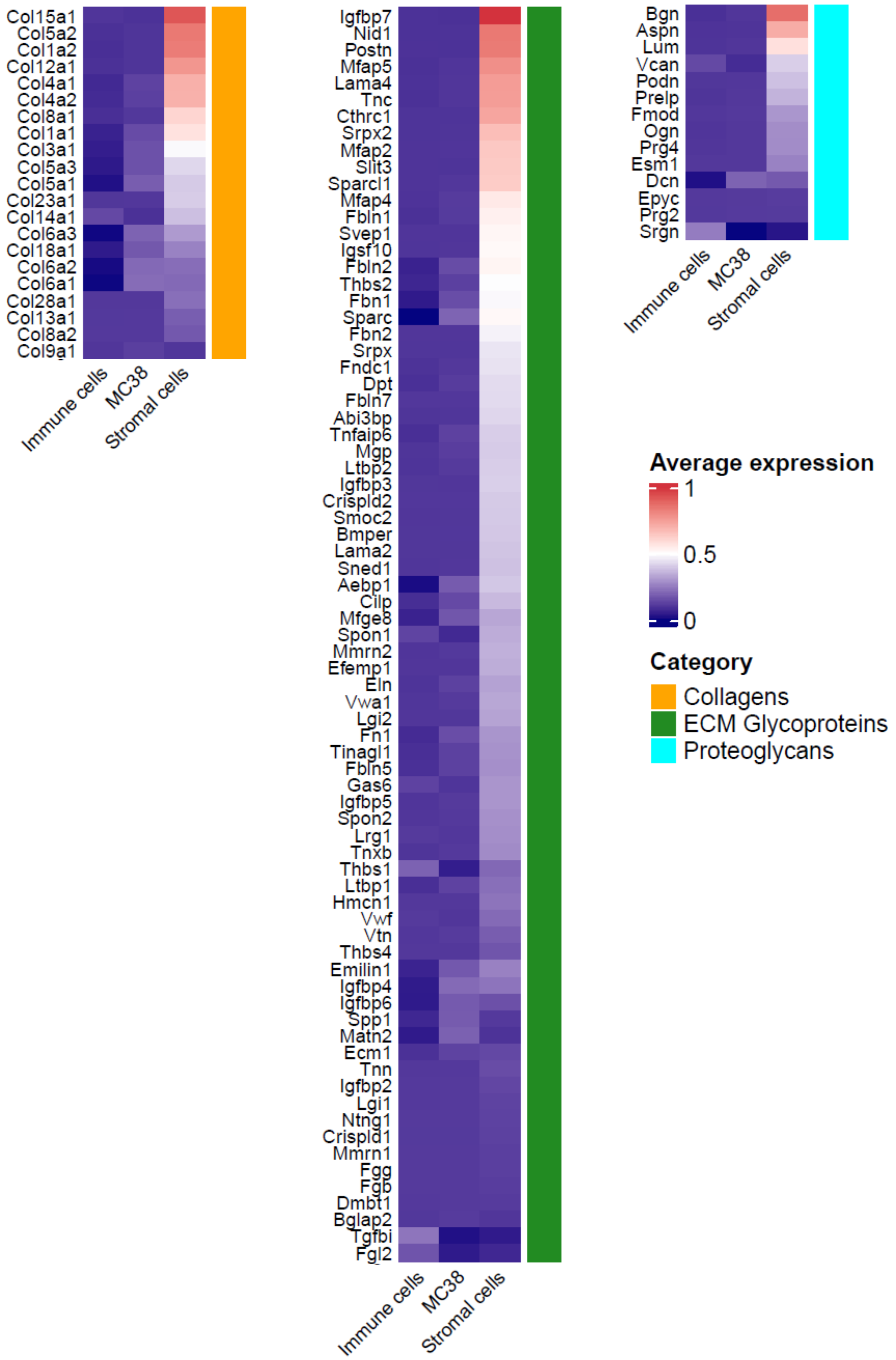

Col15a1
Col5a2
Col1a2
Col12a1
Col4a1
Col4a2
Col8a1
Col1a1
Col3a1
Col5a3
Col5a1
Col23a1
Col14a1
Col6a3
Col18a1
Col6a2
Col6a1
Col28a1
Col13a1
Col8a2
Col9a1
Immune cells
MC38
Stromal cells
Igfbp7
Nid1
Postn
Mfap5
Lama4
Tnc
Cthrc1
Srpx2
Mfap2
Slit3
Sparcl1
Mfap4
Fbln1
Svep1
Igsf10
Fbln2
Thbs2
Fbn1
Sparc
Fbn2
Srpx
Fndc1
Dpt
Fbln7
Abi3bp
Tnfaip6
Mgp
Ltbp2
Igfbp3
Crispld2
Smoc2
Bmper
Lama2
Sned1
Aebp1
Cilp
Mfge8
Spon1
Mmrn2
Efemp1
Eln
Vwa1
Lgi2
Fn1
Tinagl1
Fbln5
Gas6
Igfbp5
Spon2
Lrg1
Tnxb
Thbs1
Ltbp1
Hmcn1
Vwf
Vtn
Thbs4
Emilin1
Igfbp4
Igfbp6
Spp1
Matn2
Ecm1
Tnn
Igfbp2
Lgi1
Ntng1
Crispld1
Mmrn1
Fgg
Fgb
Dmbt1
Bglap2
Tgfbi
Fgl2
Immune cells
MC38
Stromal cells
Bgn
Aspn
Lum
Vcan
Podn
Prelp
Fmod
Ogn
Prg4
Esm1
Dcn
Epyc
Prg2
Srgn
Immune cells
MC38
Stromal cells
Average expression
1
0.5
0
Category
Collagens
ECM Glycoproteins
Proteoglycans

S1f

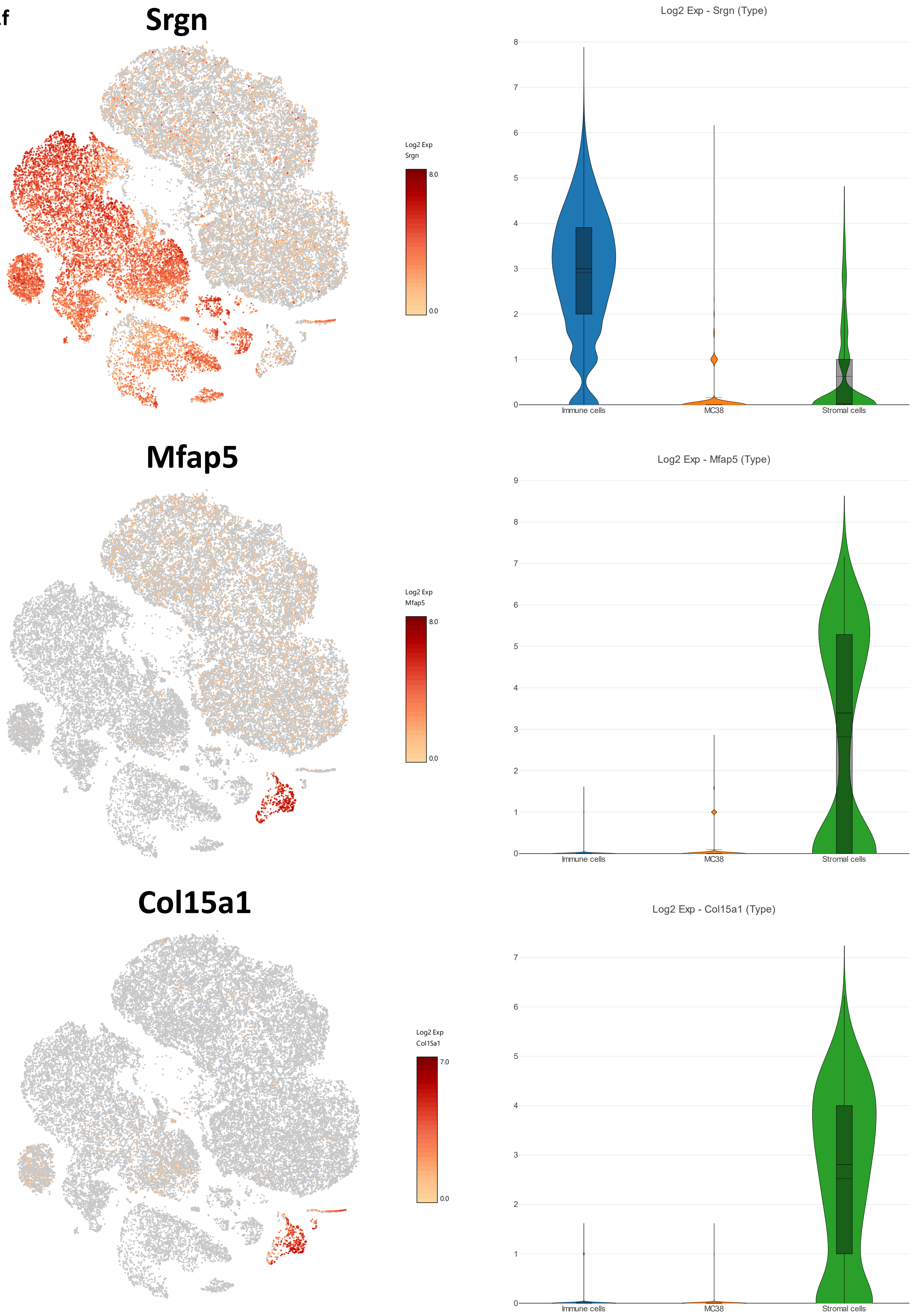
Srgn
Log2 Exp
Srgn
8.0
0.0
Log2 Exp - Srgn (Type)
8
7
6
5
4
3
2
1
0
Immune cells
MC38
Stromal cells
Mfap5
Log2 Exp
Mfap5
8.0
0.0
Log2 Exp - Mfap5 (Type)
9
8
7
6
5
4
3
2
1
0
Immune cells
MC38
Stromal cells
Col15a1
Log2 Exp
Col15a1
7.0
0.0
Log2 Exp - Col15a1 (Type)
7
6
5
4
3
2
1
0
Immune cells
MC38
Stromal cells

**S1f (cont.)**

### Fgl2

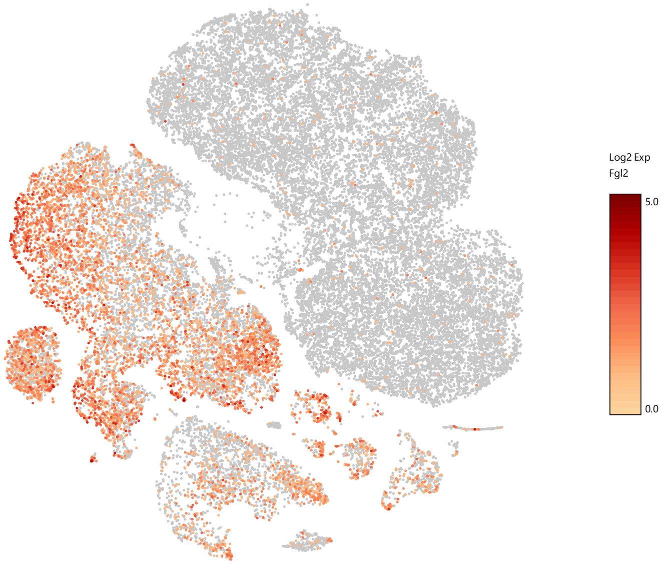


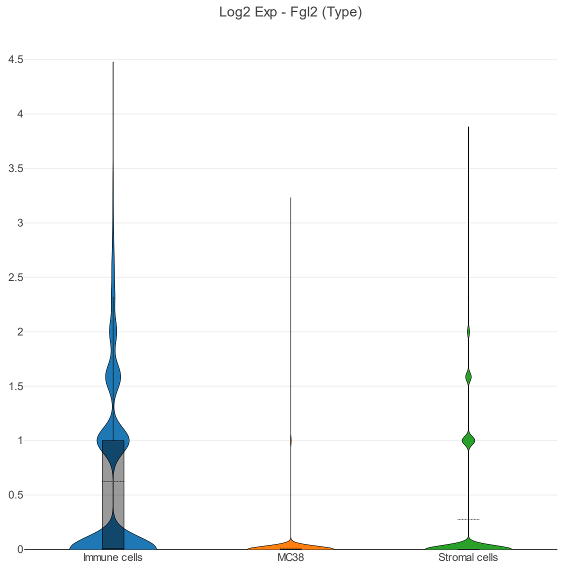


### Spon1

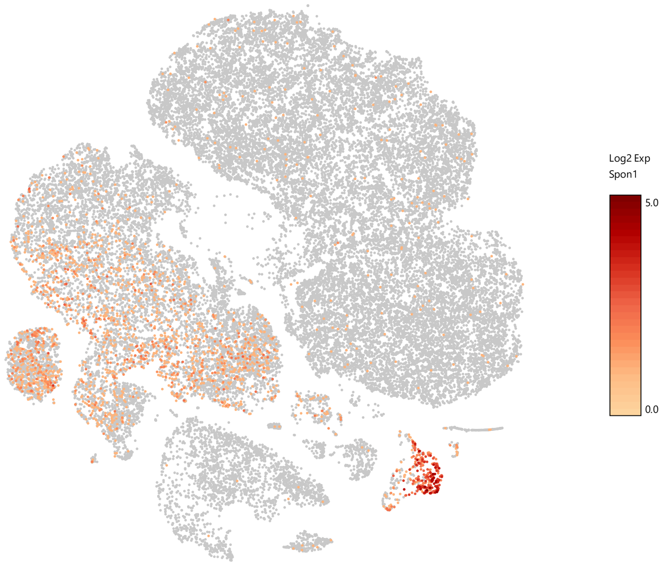


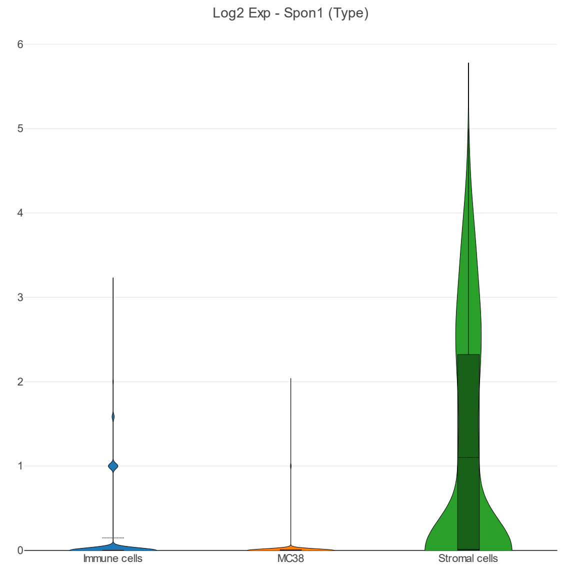

S1g

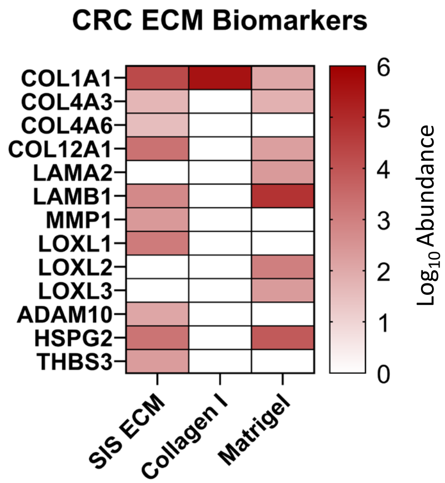
CRC ECM Biomarkers
COL1A1
COL4A3
COL4A6
COL12A1
LAMA2
LAMB1
MMP1
LOXL1
LOXL2
LOXL3
ADAM10
HSPG2
THBS3
SIS ECM
Collagen I
Matrigel
6
5
4
3
2
1
0
Log10 Abundance

**Figure S1**: a. Top 10 CRC core matrisome genes based upon bulk RNA Seq of cells alone spheroids at Day 7 of culture. b. Heatmap of core matrisome genes and their relative expression patterns. Expression data was normalized within each cell line column. c. Bar chart showing the overlap between core matrisome proteins identified within ECM biomaterials (LC-MS) vs. the number of CRC core matrisome genes identified from the cells alone spheroids. d. Percentage similarity between the ECM biomaterials and CRC spheroids. e. Core matrisome expression by cell type from in vivo MC38 tumors determined by scRNA Seq. f. tSNE and violin plots of relative expression of the top 5 matrisome genes predominantly expressed by non-cancer cells within in vivo MC38 tumors. g. Heatmap of CRC ECM biomarkers found within ECM biomaterials.

S2a

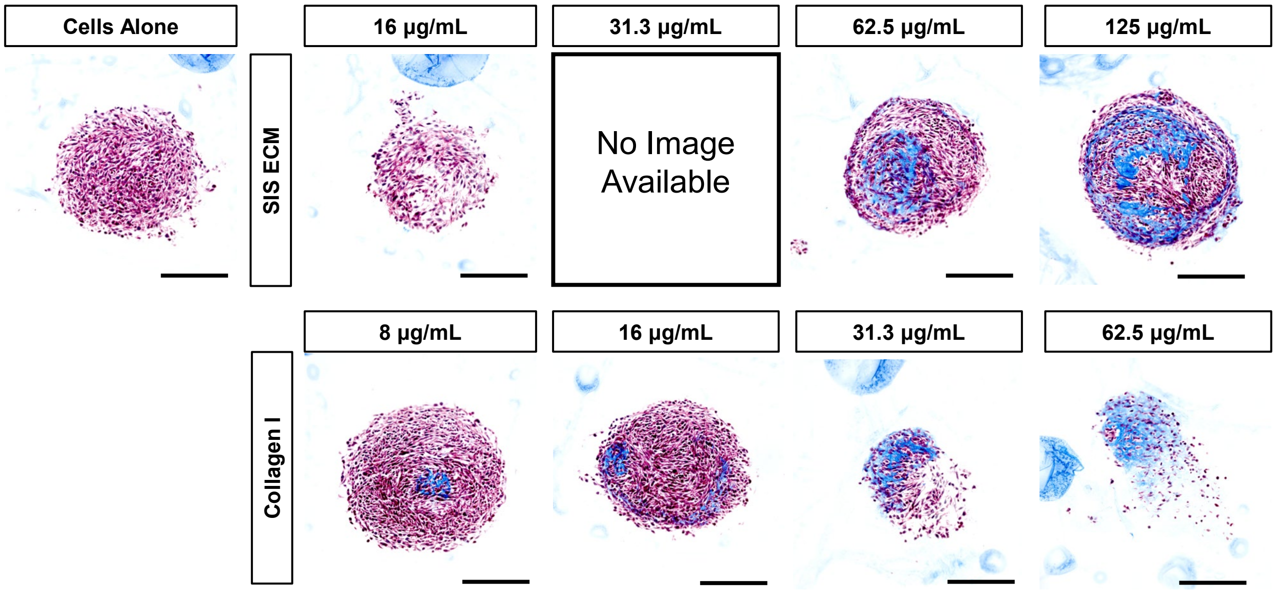
Cells Alone
16 µg/mL
31.3 µg/mL
62.5 µg/mL
125 µg/mL
SIS ECM
No Image Available
8 µg/mL
16 µg/mL
31.3 µg/mL
62.5 µg/mL
Collagen I

S2b

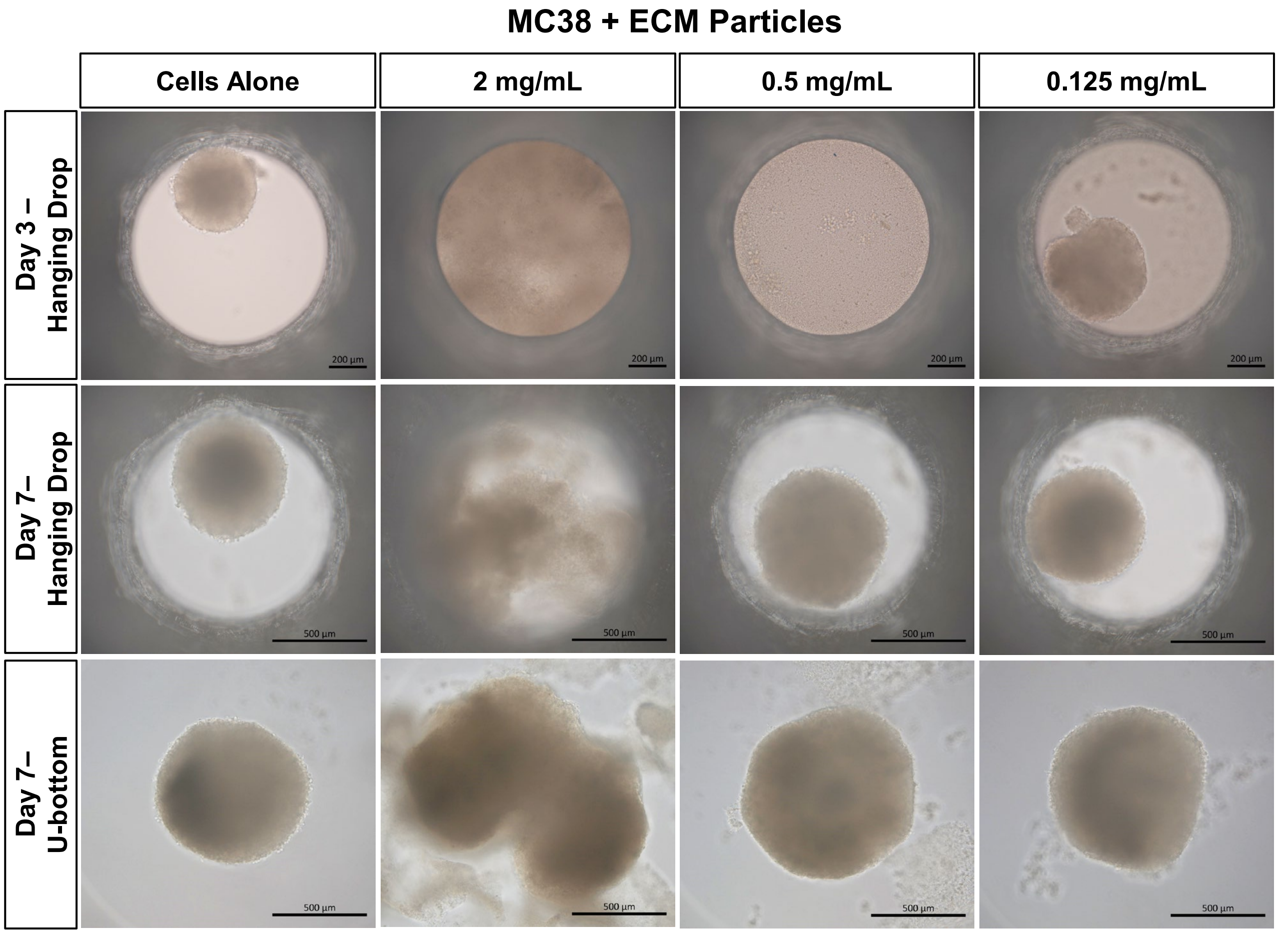
MC38 + ECM Particles
Cells Alone
2 mg/mL
0.5 mg/mL
0.125 mg/mL
Day 3 – Hanging Drop
Day 7– Hanging Drop
Day 7– U-bottom
200 µm
500 µm

S2c

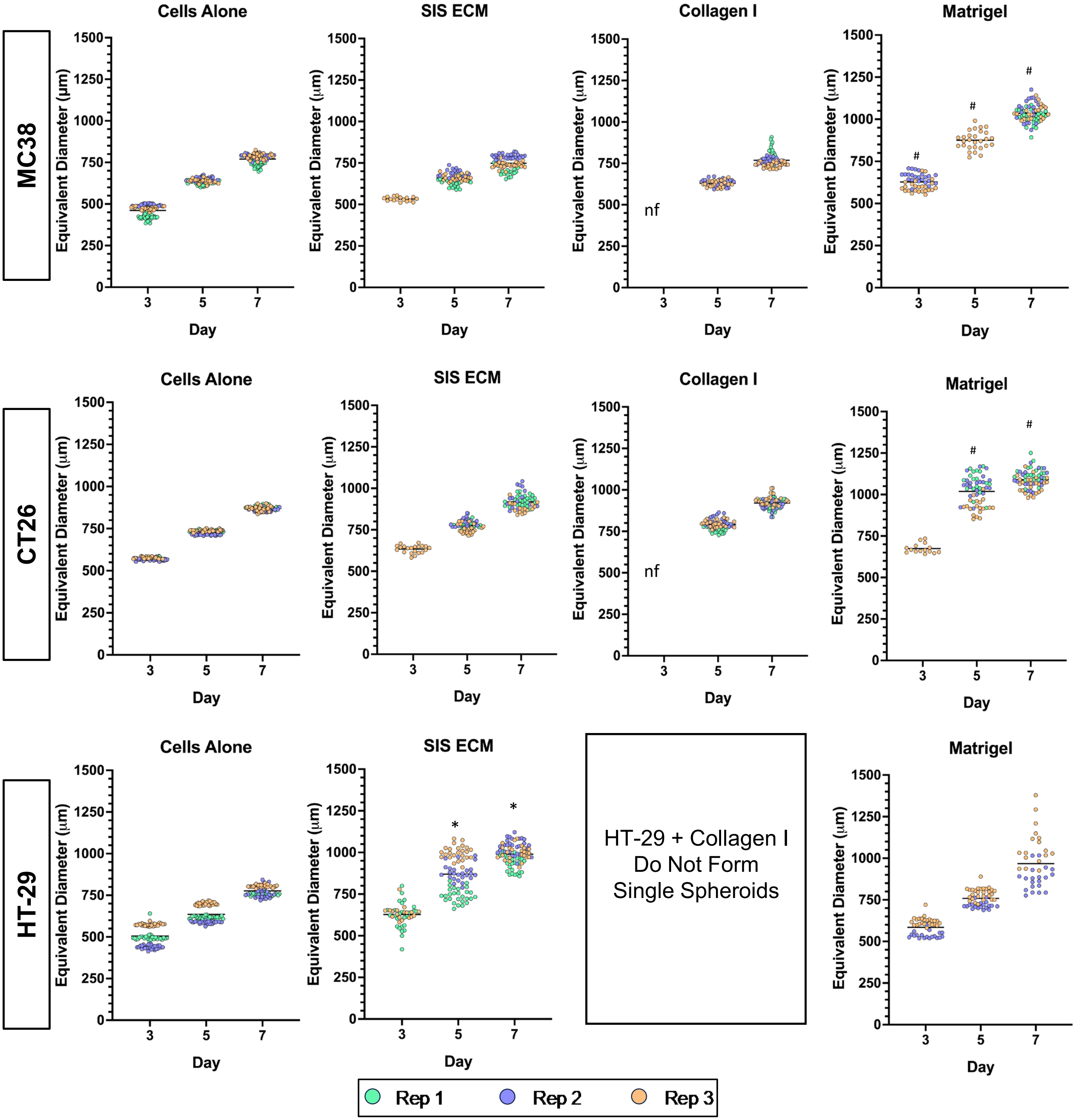
Cells Alone
SIS ECM
Collagen I
Matrigel
MC38
CT26
HT-29
Equivalent Diameter (µm)
Day
nf
#
*
HT-29 + Collagen I Do Not Form Single Spheroids
Rep 1
Rep 2
Rep 3

S2d

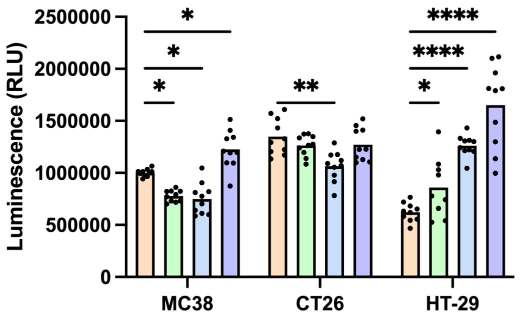
Luminescence (RLU)
*
**
****
MC38
CT26
HT-29

S2e

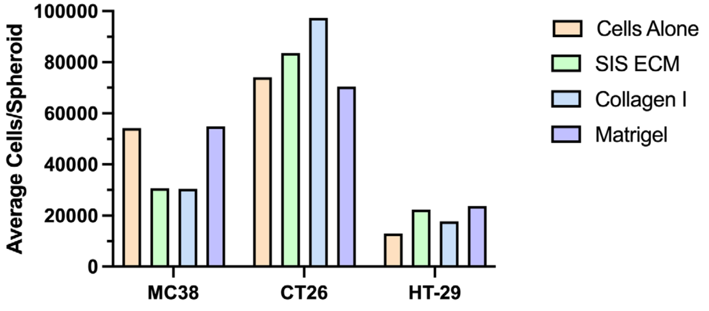
Average Cells/Spheroid
MC38
CT26
HT-29
Cells Alone
SIS ECM
Collagen I
Matrigel

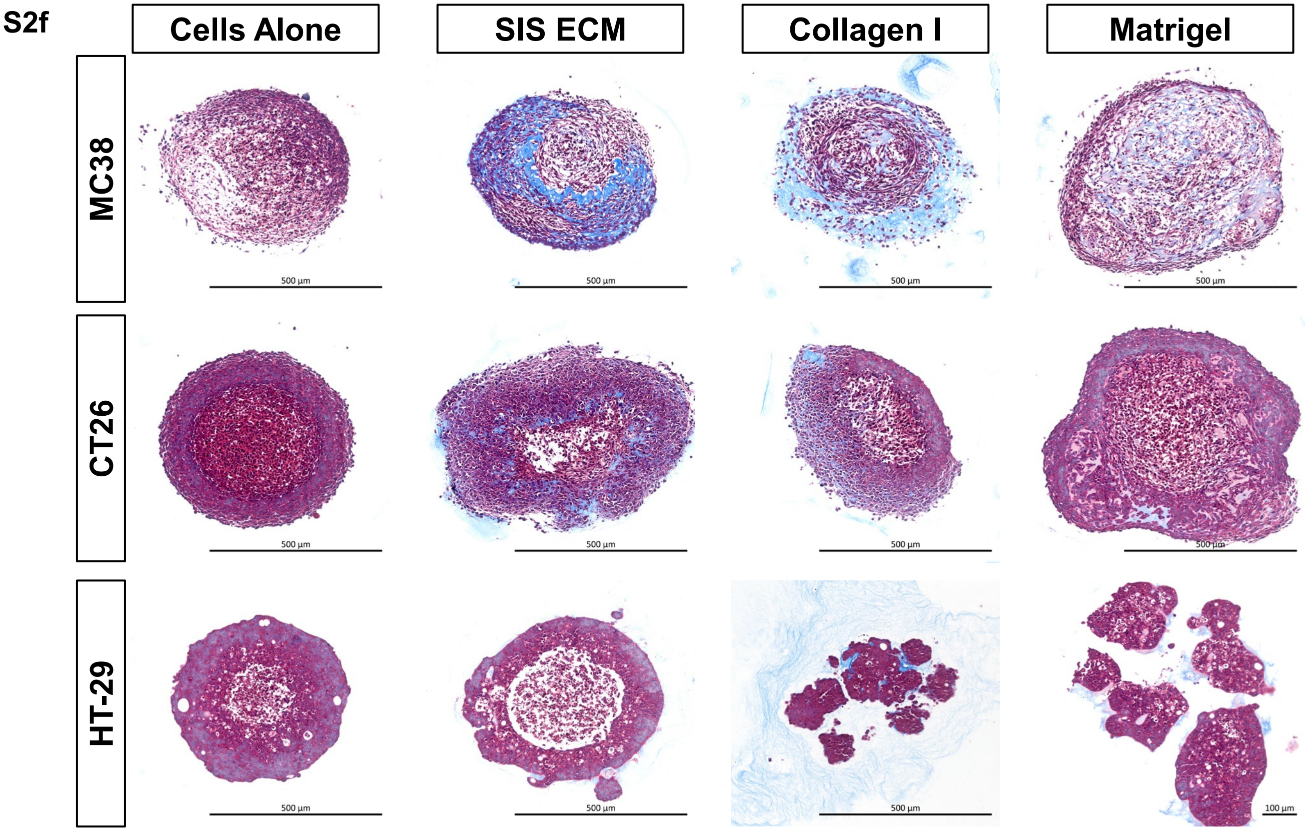
S2f
Cells Alone
SIS ECM
Collagen I
Matrigel
MC38
CT26
HT-29
500 µm
100 µm

**Figure S2**: a. Masson's Trichrome stained spheroid sections show varying concentrations of SIS ECM digest and Type I Collagen integrate within MC38 MatriSpheres. b. SIS ECM particles were combined with MC38 cells to form spheroids via hanging drop or ultra-low attachment round-bottom 96-well plates. c. Quantification of equivalent diameters at days 3, 5, and 7 of culture, n >= 37. Statistics are one-way ANOVA followed by Sídák's multiple comparisons test. # = Matrigel statistically different from cells alone on the same day, * = SIS ECM statistically different from cells alone on the same day. d. Quantification of metabolic activity via CellTiter-Glo 3D assay at day 7. Statistics were calculated by a two-way ANOVA followed by a Tukey's multiple comparisons test. e. Average number of cells per spheroid counted from manual dissociation of spheroids at day 7. f. Representative images of Masson's Trichrome stained spheroid sections at day 7. Scale bars are indicated on the images. Statistical significance where $p < 0.05$ is denoted with *, ≤ 0.01 with **, ≤ 0.001 with ***, and ≤ 0.0001 with ****.

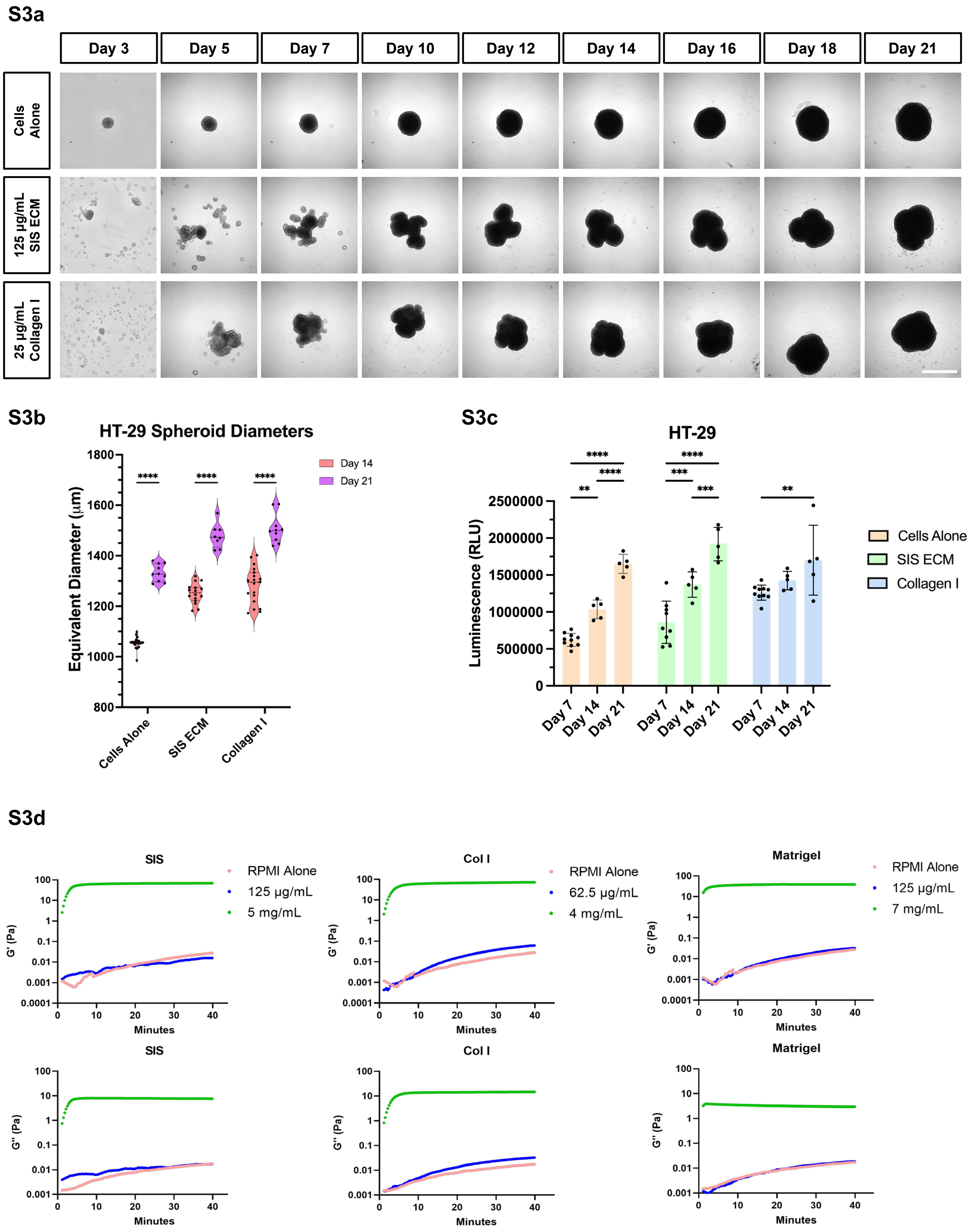
S3a
Day 3
Day 5
Day 7
Day 10
Day 12
Day 14
Day 16
Day 18
Day 21
Cells Alone
125 µg/mL SIS ECM
25 µg/mL Collagen I
S3b
HT-29 Spheroid Diameters
Equivalent Diameter (µm)
Day 14
Day 21
****
Cells Alone
SIS ECM
Collagen I
S3c
HT-29
Luminescence (RLU)
Cells Alone
SIS ECM
Collagen I
Day 7
Day 14
Day 21
S3d
SIS
Col I
Matrigel
RPMI Alone
125 µg/mL
5 mg/mL
62.5 µg/mL
4 mg/mL
7 mg/mL
G' (Pa)
G'' (Pa)
Minutes

**S3e**

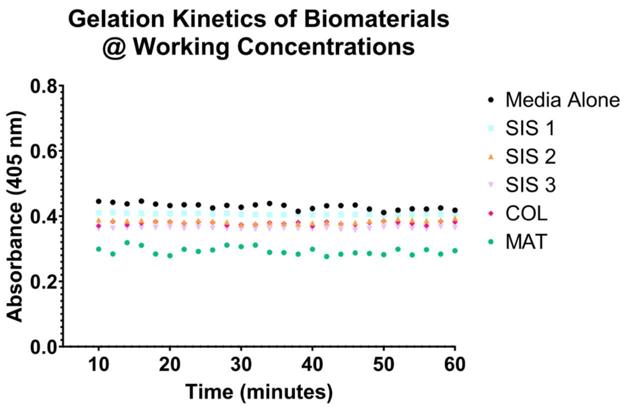

Gelation Kinetics of Biomaterials @ Working Concentrations
Absorbance (405 nm)
0.8
0.6
0.4
0.2
0.0
10
20
30
40
50
60
Time (minutes)
Media Alone
SIS 1
SIS 2
SIS 3
COL
MAT


**S3f**

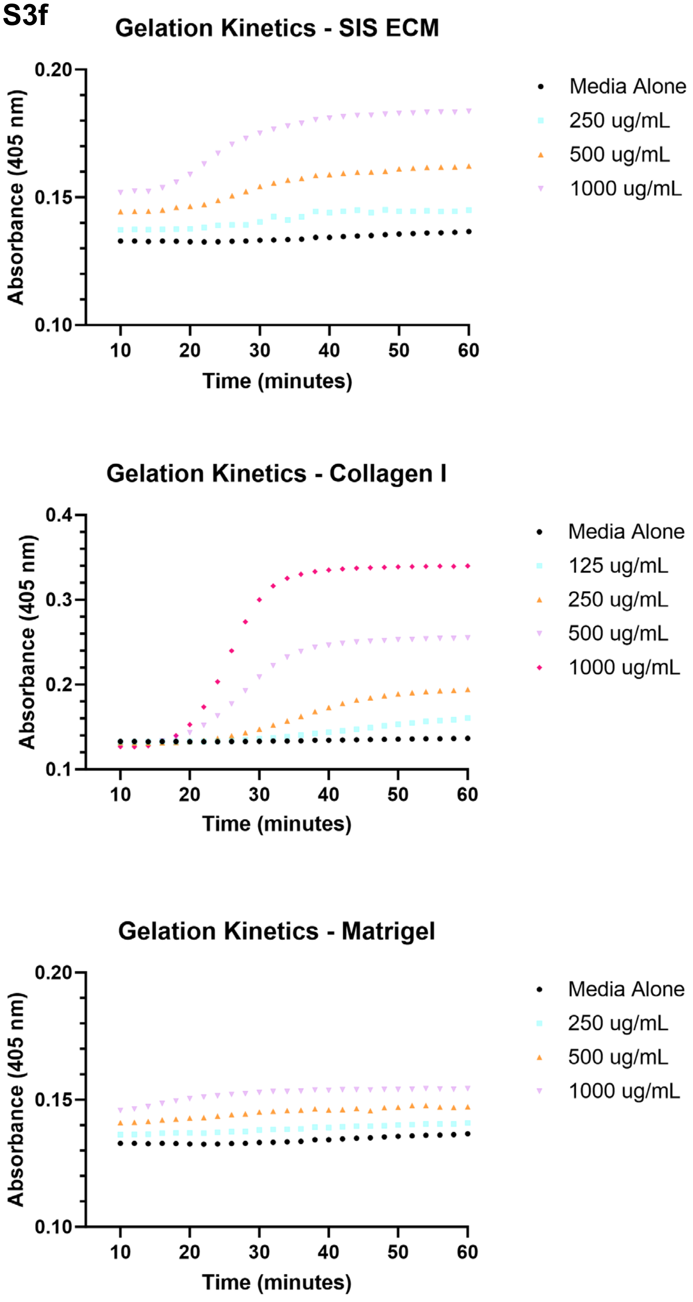

Gelation Kinetics - SIS ECM
Absorbance (405 nm)
0.20
0.15
0.10
10
20
30
40
50
60
Time (minutes)
Media Alone
250 ug/mL
500 ug/mL
1000 ug/mL
Gelation Kinetics - Collagen I
Absorbance (405 nm)
0.4
0.3
0.2
0.1
Time (minutes)
Media Alone
125 ug/mL
250 ug/mL
500 ug/mL
1000 ug/mL
Gelation Kinetics - Matrigel
Absorbance (405 nm)
0.20
0.15
0.10
Time (minutes)
Media Alone
250 ug/mL
500 ug/mL
1000 ug/mL

**Figure S3**: A. Representative images of HT-29 spheroids grown for 21 days. b. Quantification of HT-29 MatriSphere diameters at days 14 and 21. Statistics were calculated by a two-way ANOVA followed by a Sídák’s multiple comparisons test. c. Quantification of metabolic activity of spheroids via CellTiter-Glo 3D. d. Rheological characterization of cell culture media and ECM biomaterials at experimental and traditional hydrogel formation concentrations. e. Gelation kinetics curves showing turbidity of ECM biomaterials at working ECM concentrations in comparison to control RPMI media. f. Gelation kinetics curves at varying ECM concentration by ECM biomaterial. Statistical significance where $p < 0.05$ is denoted with *, ≤ 0.01 with **, ≤ 0.001 with ***, and ≤ 0.0001 with ****.

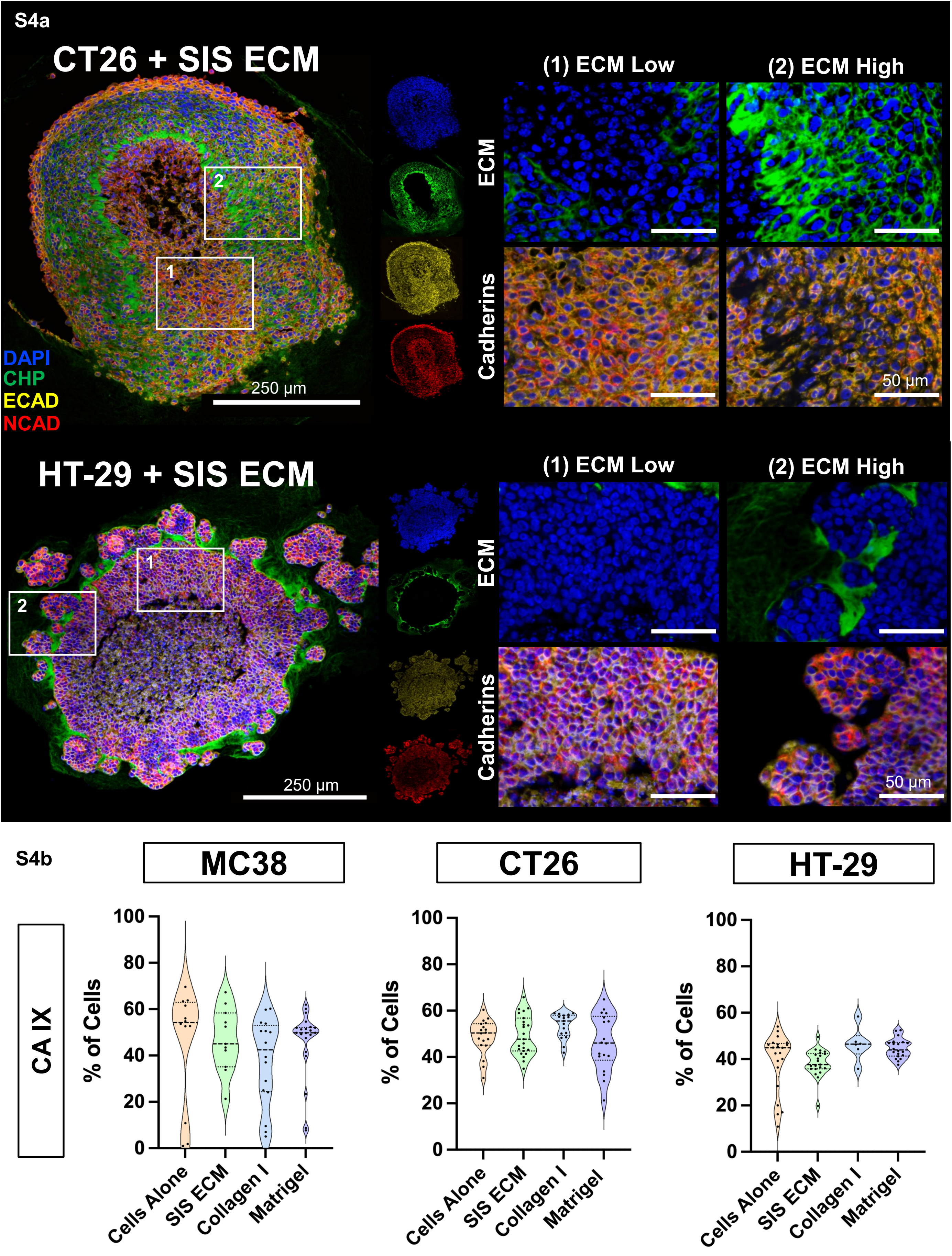

S4a
CT26 + SIS ECM
(1) ECM Low
(2) ECM High
ECM
Cadherins
DAPI
CHP
ECAD
NCAD
250 µm
50 µm
HT-29 + SIS ECM
(1) ECM Low
(2) ECM High
ECM
Cadherins
250 µm
50 µm
S4b
MC38
CT26
HT-29
CA IX
% of Cells
0
20
40
60
80
100
Cells Alone
SIS ECM
Collagen I
Matrigel

**S4c**

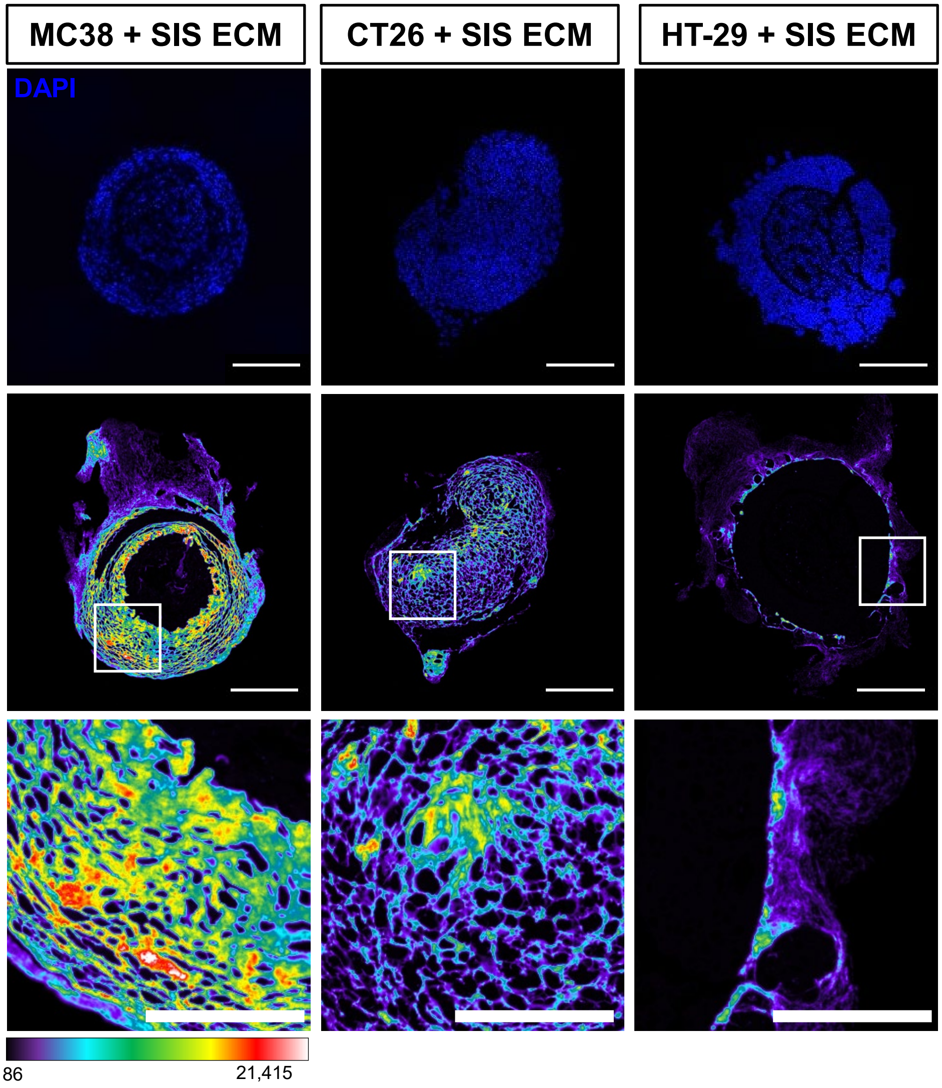

## MC38

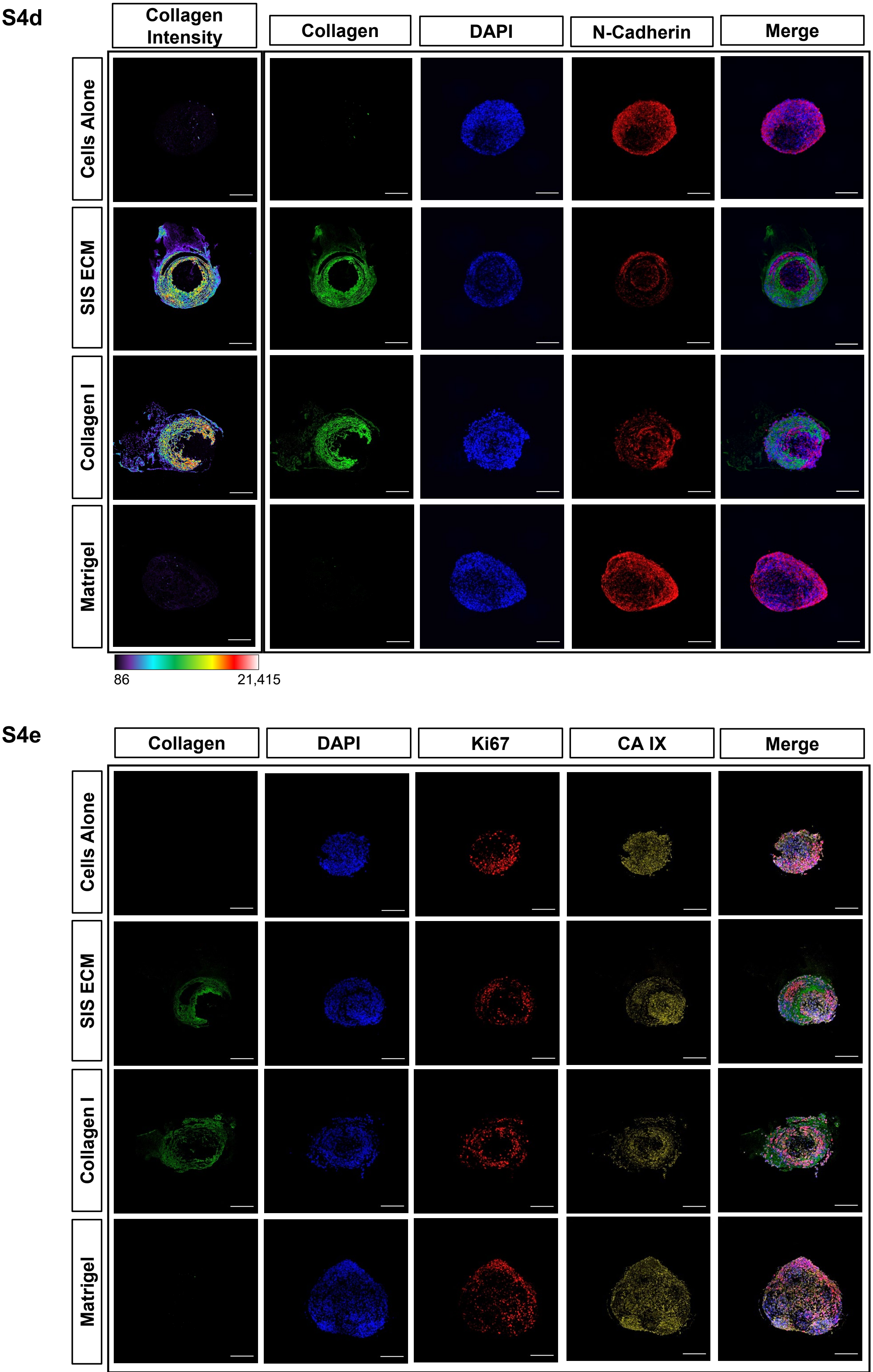

S4d
Collagen Intensity
Collagen
DAPI
N-Cadherin
Merge
Cells Alone
SIS ECM
Collagen I
Matrigel
86
21,415
S4e
Collagen
DAPI
Ki67
CA IX
Merge
Cells Alone
SIS ECM
Collagen I
Matrigel

# CT26

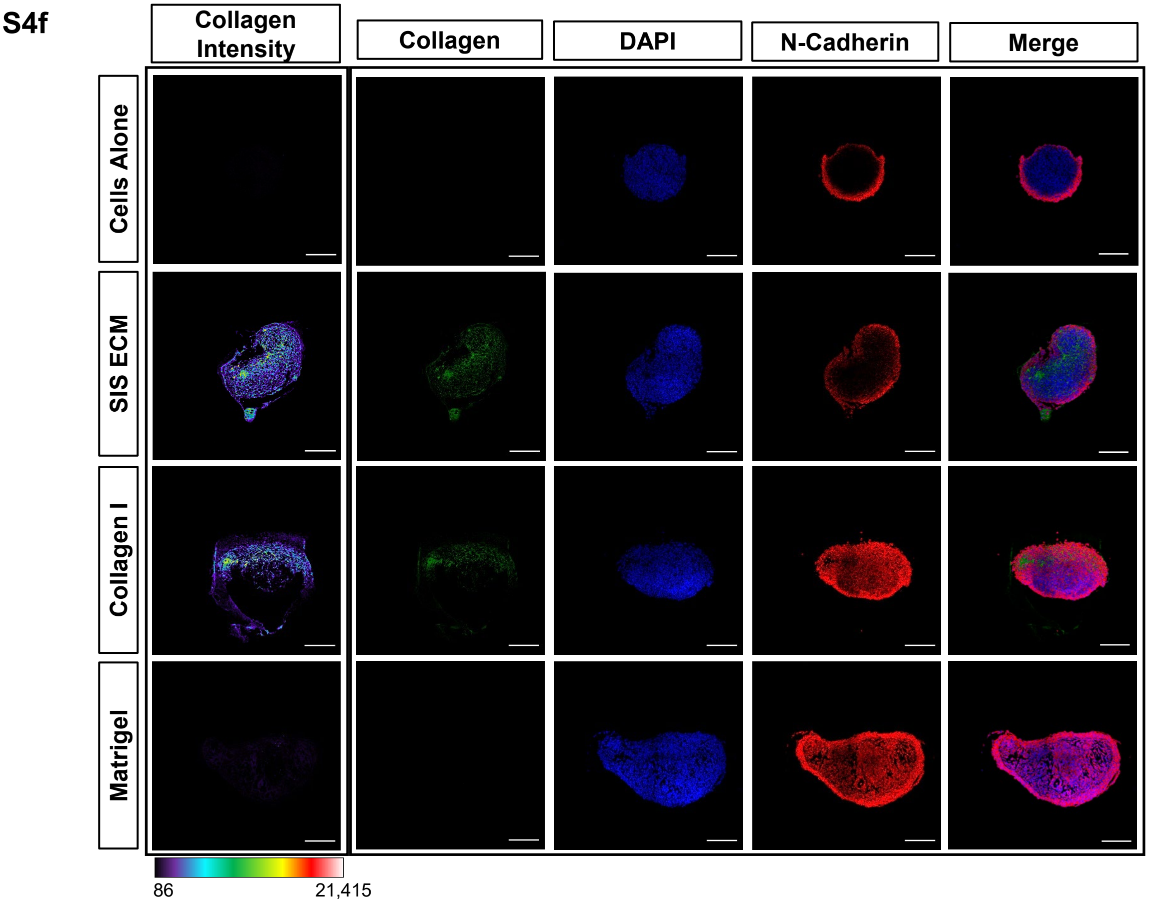
S4f
Collagen Intensity
Collagen
DAPI
N-Cadherin
Merge
Cells Alone
SIS ECM
Collagen I
Matrigel
86
21,415

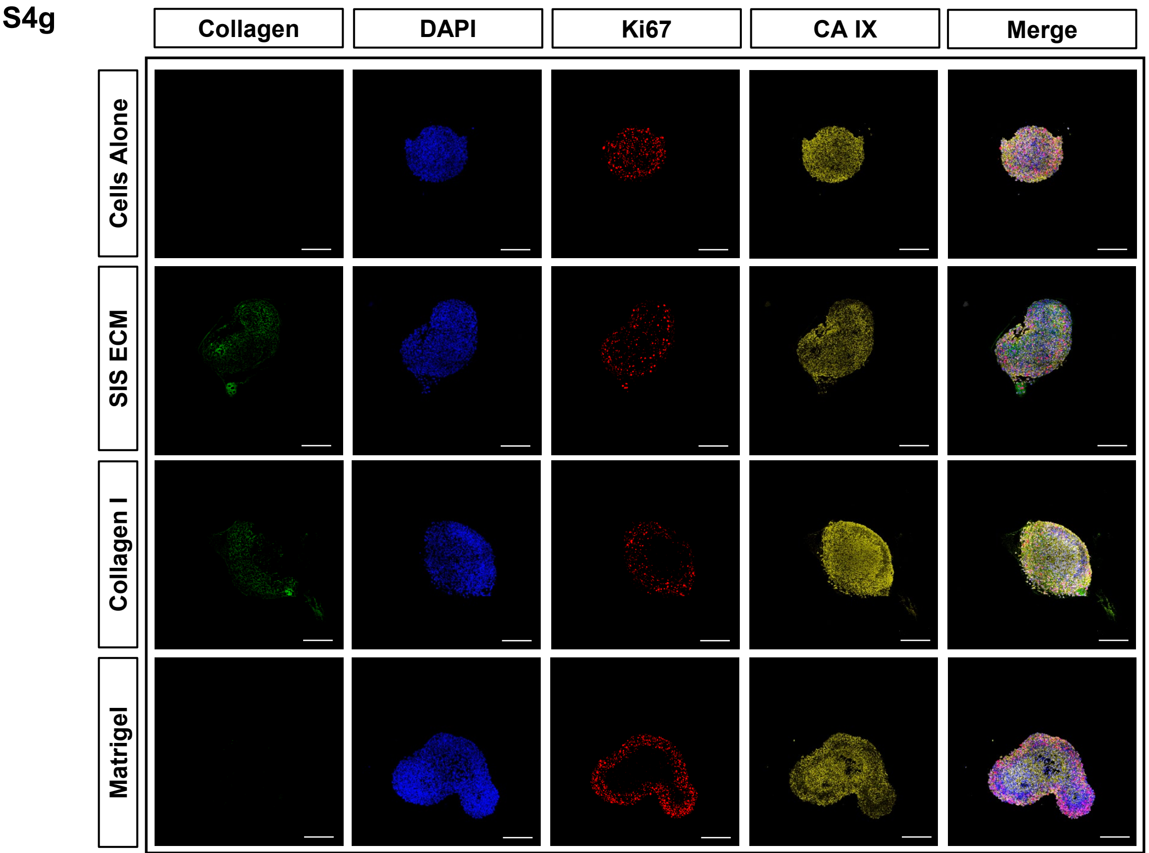
S4g
Collagen
DAPI
Ki67
CA IX
Merge
Cells Alone
SIS ECM
Collagen I
Matrigel

## HT-29

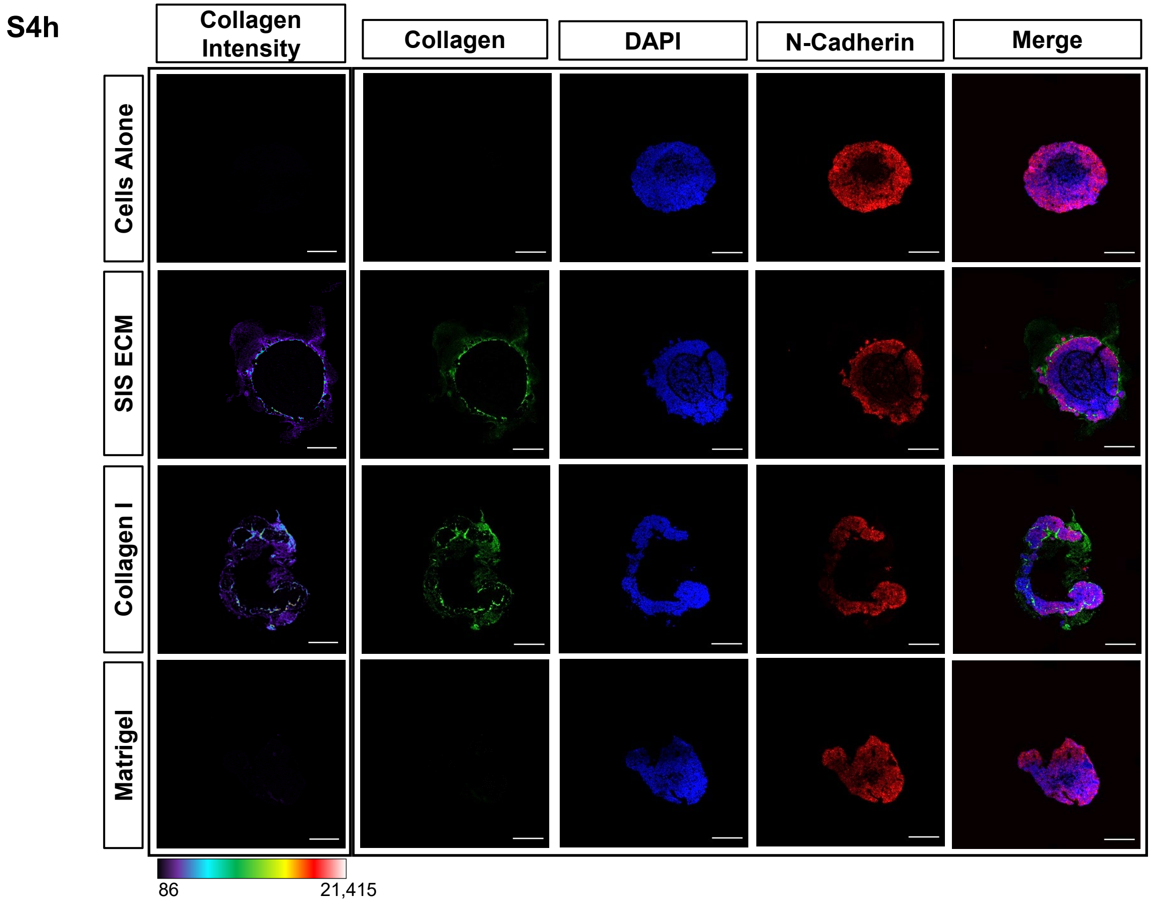
S4h
Collagen Intensity
Collagen
DAPI
N-Cadherin
Merge
Cells Alone
SIS ECM
Collagen I
Matrigel
86
21,415

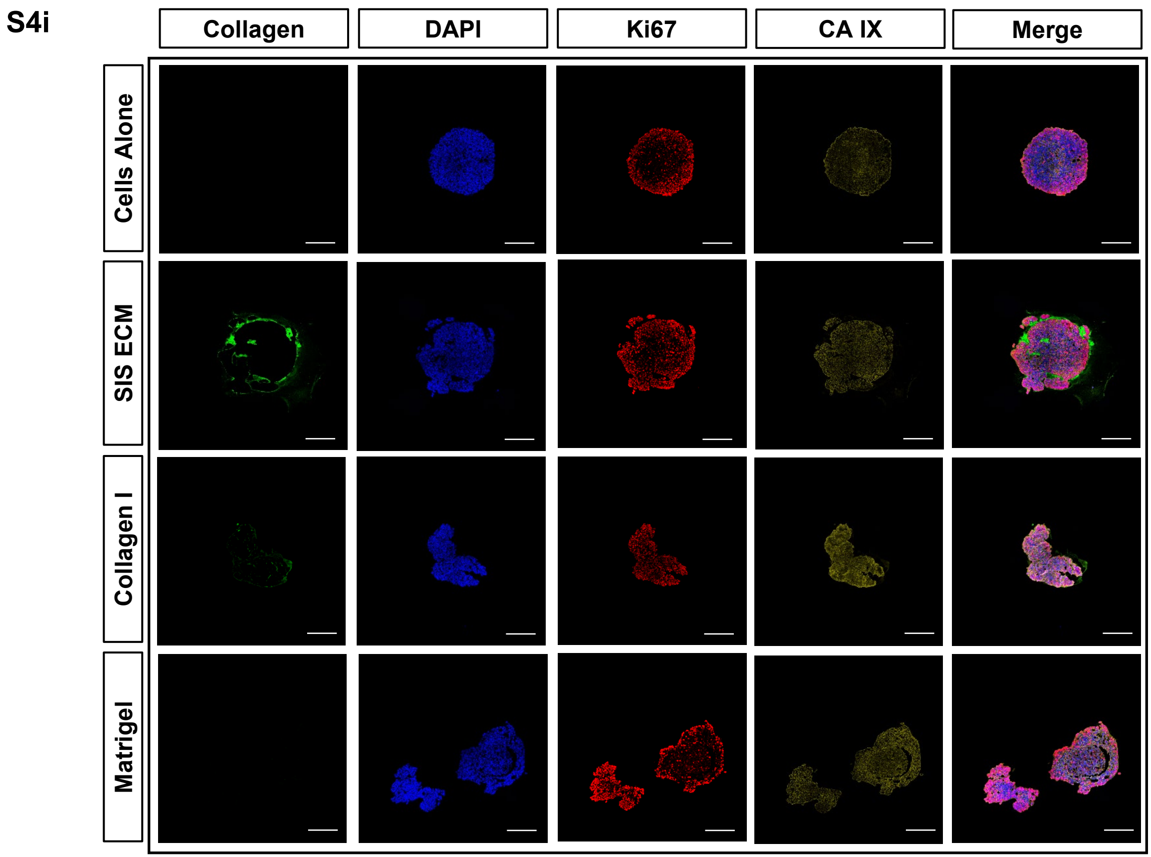
S4i
Collagen
DAPI
Ki67
CA IX
Merge
Cells Alone
SIS ECM
Collagen I
Matrigel

**Figure S4**: a. Representative images of CT26 + SIS ECM & HT-29 + SIS ECM spheroid sections stained for E-CAD, N-CAD, CHP, and DAPI. b. Quantification of CA IX staining from representative spheroid sections. Statistics were calculated by a one-way ANOVA followed by a Tukey's multiple comparisons test. c. Heatmaps of SIS ECM-incorporated spheroid sections stained with CHP to visualize denatured fibrillar collagen regions. d. Representative images of MC38 spheroid sections showing collagen intensity, CHP, DAPI, N-CAD, and the merged image. e. Representative images of MC38 spheroid sections showing, CHP, DAPI, Ki67, CA IX, and the merged image. f. Representative images of CT26 spheroid sections showing collagen intensity, CHP, DAPI, N-CAD, and the merged image. g. Representative images of CT26 spheroid sections showing, CHP, DAPI, Ki67, CA IX, and the merged image. h. Representative images of HT-29 spheroid sections showing collagen intensity, CHP, DAPI, N-CAD, and the merged image. i. Representative images of HT-29 spheroid sections showing, CHP, DAPI, Ki67, CA IX, and the merged image. Statistical significance where $p < 0.05$ is denoted with *, ≤ 0.01 with **, ≤ 0.001 with ***, and ≤ 0.0001 with ****.

S5a

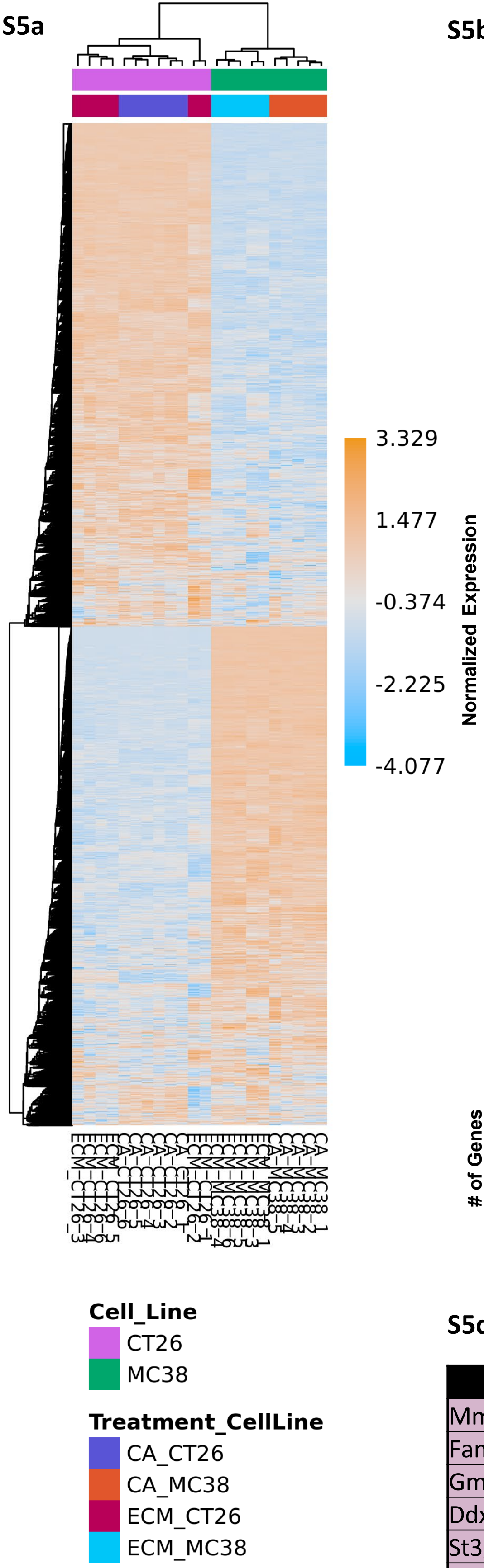


S5b

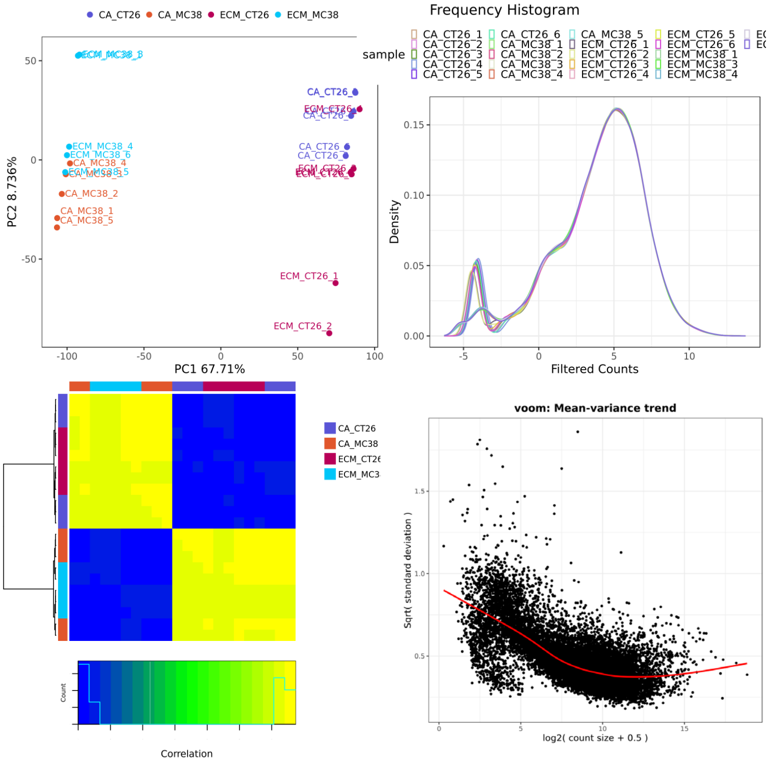


| | ECM_MC38 -CA_MC38 | ECM_CT26 -CA_CT26 |
|---|---|---|
| *upreg>1.2, pval<0.05* | 938 | 1066 |
| *downreg<-1.2, pval<0.05* | 1069 | 916 |
| *upreg>1.2, pval<0.01* | 580 | 764 |
| *downreg<-1.2, pval<0.01* | 667 | 576 |
| *upreg>1.2, adjpval<0.05* | 431 | 678 |
| *downreg<-1.2, adjpval<0.05* | 506 | 494 |
| *upreg>1.2, adjpval<0.01* | 178 | 418 |
| *downreg<-1.2, adjpval<0.01* | 216 | 262 |

Please note Sample size:
CA_CT26=6
CA_MC38=5
ECM_CT26=6
ECM_MC38=5

Contrasts:
ECM_MC38-CA_MC38 : c(ECM_MC38 = 5, CA_MC38 = 5)
ECM_CT26-CA_CT26 : c(ECM_CT26 = 6, CA_CT26 = 6)

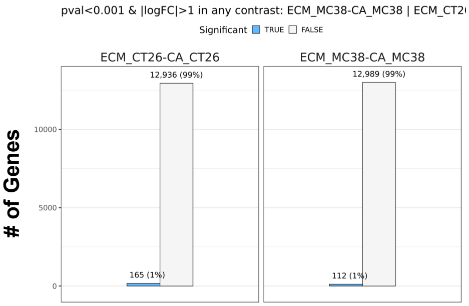


S5c

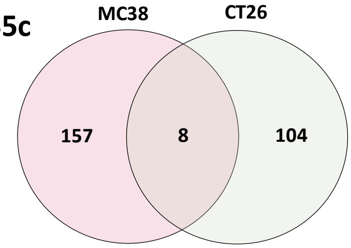


| Genes Influenced by ECM | |
|---|---|
| Mmp13 | St3gal4 |
| Fam102a | Dlgap2 |
| Gm8995 | Dlx2 |
| Ddx41 | Aspn |

S5d

| | SIS ECM vs Cells Alone | | | |
|---|---|---|---|---|
| | MC38 | | CT26 | |
| Genes | Fold Change | Adj p-value | Fold Change | Adj p-value |
| Mmp13 | 2.90 | 6.30E-03 | 14.22 | 2.42E-13 |
| Fam102a | 2.12 | 1.34E-08 | 2.61 | 8.98E-11 |
| Gm8995 | 2.08 | 6.68E-03 | 2.72 | 4.71E-09 |
| Ddx41 | 2.06 | 6.10E-08 | 2.00 | 1.91E-08 |
| St3gal4 | 2.05 | 1.44E-08 | 2.09 | 9.51E-09 |
| Dlgap2 | 2.03 | 2.54E-03 | 3.75 | 4.30E-03 |
| Dlx2 | -3.99 | 6.19E-04 | -5.05 | 1.64E-04 |
| Aspn | -6.13 | 5.29E-07 | -2.01 | 7.96E-03 |

S5e

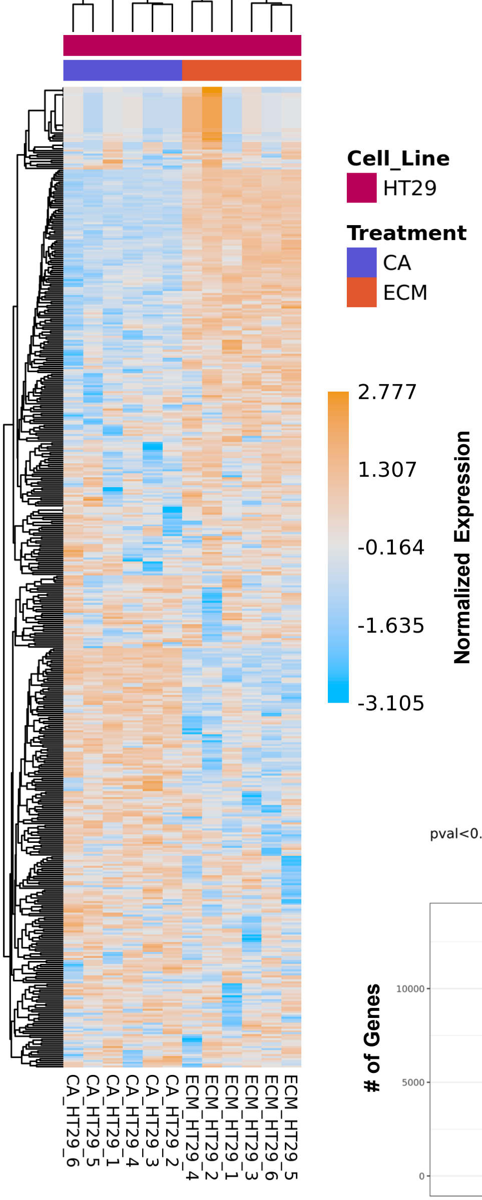
Cell_Line
HT29
Treatment
CA
ECM
2.777
1.307
-0.164
-1.635
-3.105
Normalized Expression
CA_HT29_6
CA_HT29_5
CA_HT29_1
CA_HT29_4
CA_HT29_3
CA_HT29_2
ECM_HT29_4
ECM_HT29_2
ECM_HT29_1
ECM_HT29_3
ECM_HT29_6
ECM_HT29_5

S5f

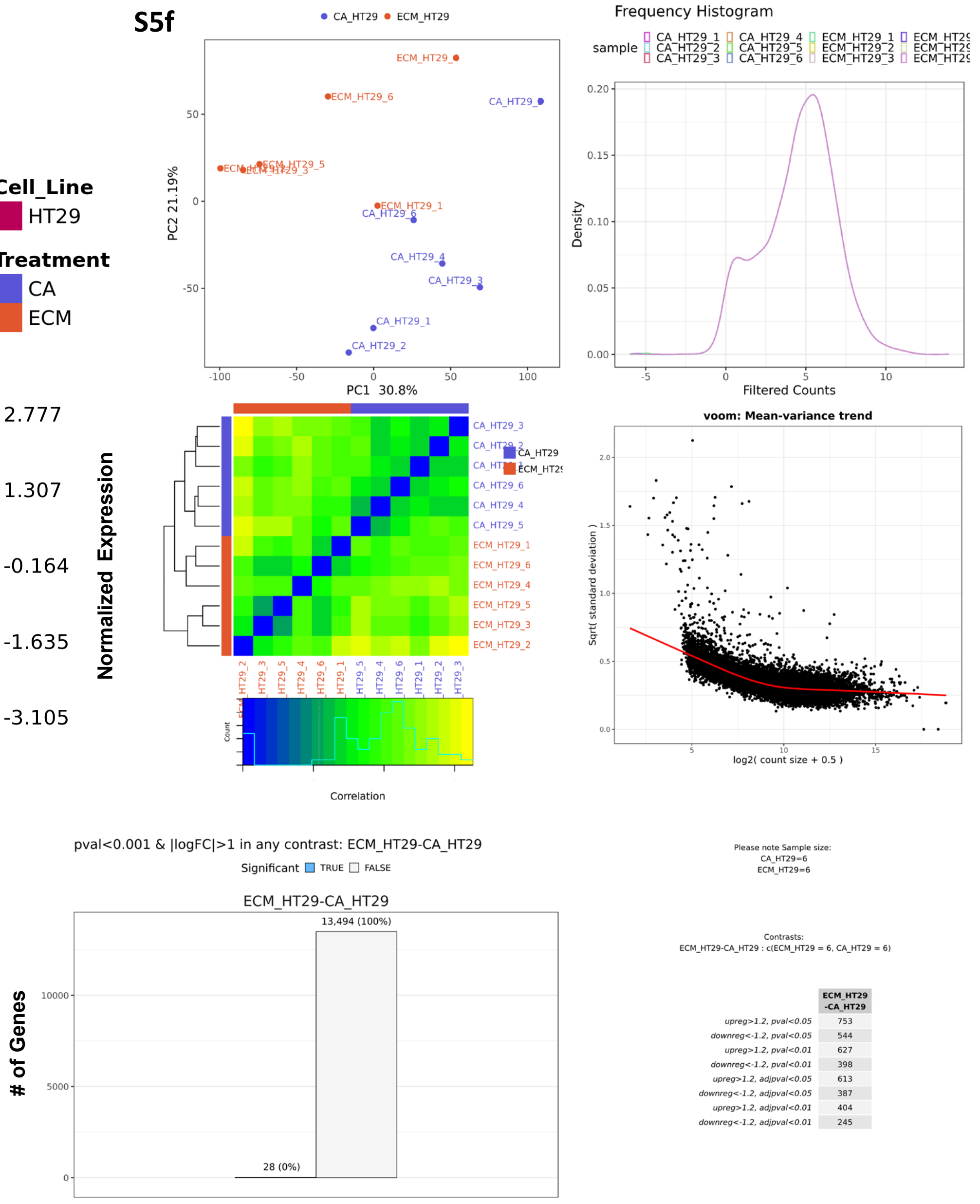
CA_HT29 ECM_HT29
PC1 30.8%
PC2 21.19%
Frequency Histogram
sample
Density
Filtered Counts
voom: Mean-variance trend
Sqrt( standard deviation )
log2( count size + 0.5 )
Correlation
Count
pval<0.001 & |logFC|>1 in any contrast: ECM_HT29-CA_HT29
Significant TRUE FALSE
ECM_HT29-CA_HT29
13,494 (100%)
28 (0%)
### of Genes
Please note Sample size:
CA_HT29=6
ECM_HT29=6
Contrasts:
ECM_HT29-CA_HT29 : c(ECM_HT29 = 6, CA_HT29 = 6)
ECM_HT29-CA_HT29
upreg>1.2, pval<0.05 753
downreg<-1.2, pval<0.05 544
upreg>1.2, pval<0.01 627
downreg<-1.2, pval<0.01 398
upreg>1.2, adjpval<0.05 613
downreg<-1.2, adjpval<0.05 387
upreg>1.2, adjpval<0.01 404
downreg<-1.2, adjpval<0.01 245

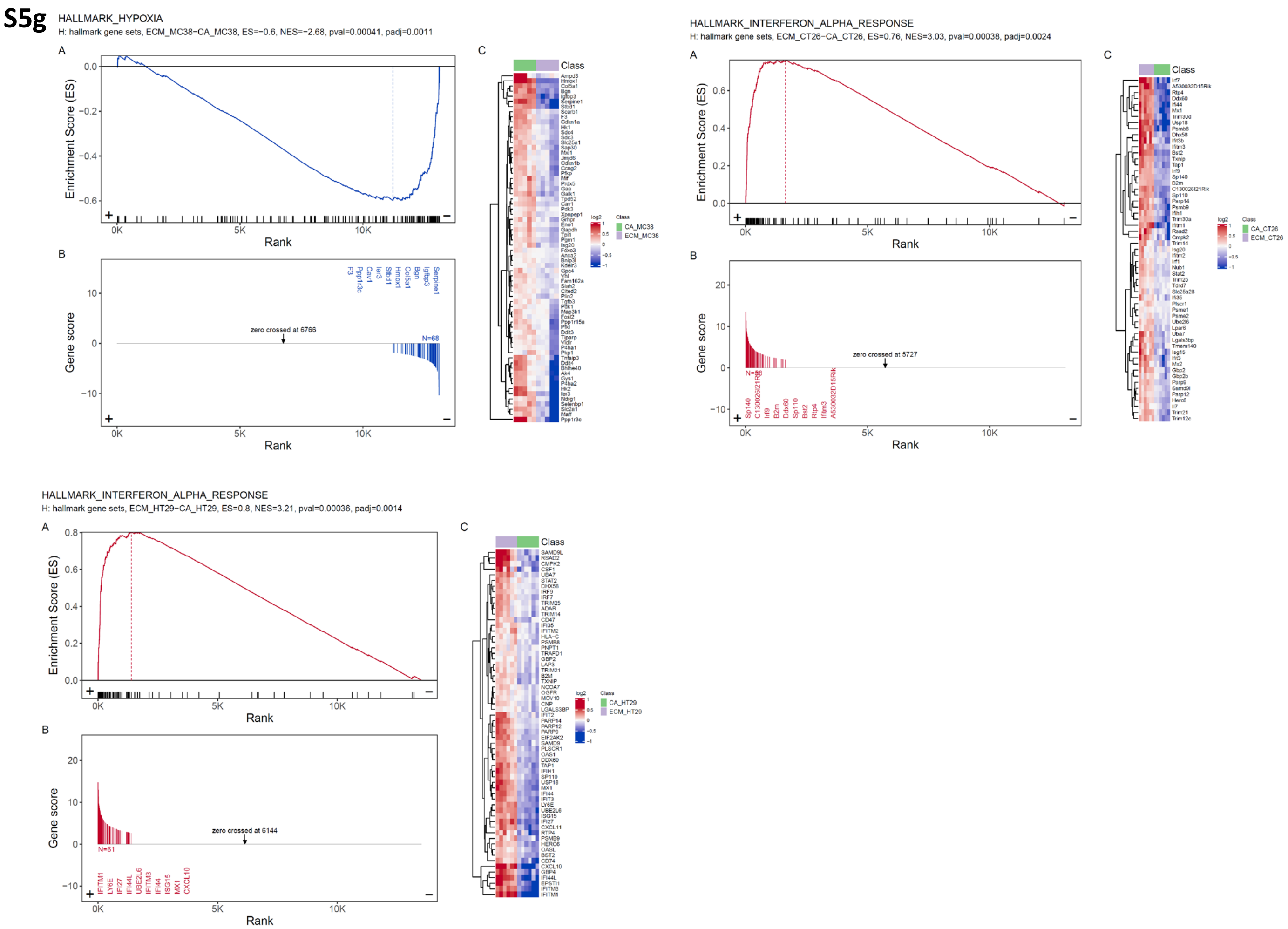
S5g
HALLMARK_HYPOXIA
H: hallmark gene sets, ECM_MC38−CA_MC38, ES=−0.6, NES=−2.68, pval=0.00041, padj=0.0011
A
Enrichment Score (ES)
Rank
B
Gene score
zero crossed at 6766
N=68
C
Class
CA_MC38
ECM_MC38
HALLMARK_INTERFERON_ALPHA_RESPONSE
H: hallmark gene sets, ECM_CT26−CA_CT26, ES=0.76, NES=3.03, pval=0.00038, padj=0.0024
A
Enrichment Score (ES)
Rank
B
Gene score
zero crossed at 5727
C
Class
CA_CT26
ECM_CT26
HALLMARK_INTERFERON_ALPHA_RESPONSE
H: hallmark gene sets, ECM_HT29−CA_HT29, ES=0.8, NES=3.21, pval=0.00036, padj=0.0014
A
Enrichment Score (ES)
Rank
B
Gene score
zero crossed at 6144
N=61
C
Class
CA_HT29
ECM_HT29

**Figure S5**: a. Heatmap showing normalized gene expression and hierarchical clustering of mouse CRC lines (MC38 and CT26) by sample. b. PCA plot, frequency histogram and number of mouse genes across different significant thresholds. c. Venn diagram of significantly overlapping genes between MC38 and CT26 d. table of corresponding fold change and adj. p-value of each gene. e. Heatmap showing normalized gene expression and hierarchical clustering of human CRC line HT-29 by sample. f. PCA plot, frequency histogram and number of human genes across different significant thresholds g. GSEA plots showing the top gene set within each cell line based upon enrichment score (ES). ES and gene score plots showing a heatmap of the leading-edge genes between treatments (SIS ECM vs. cells alone spheroids).

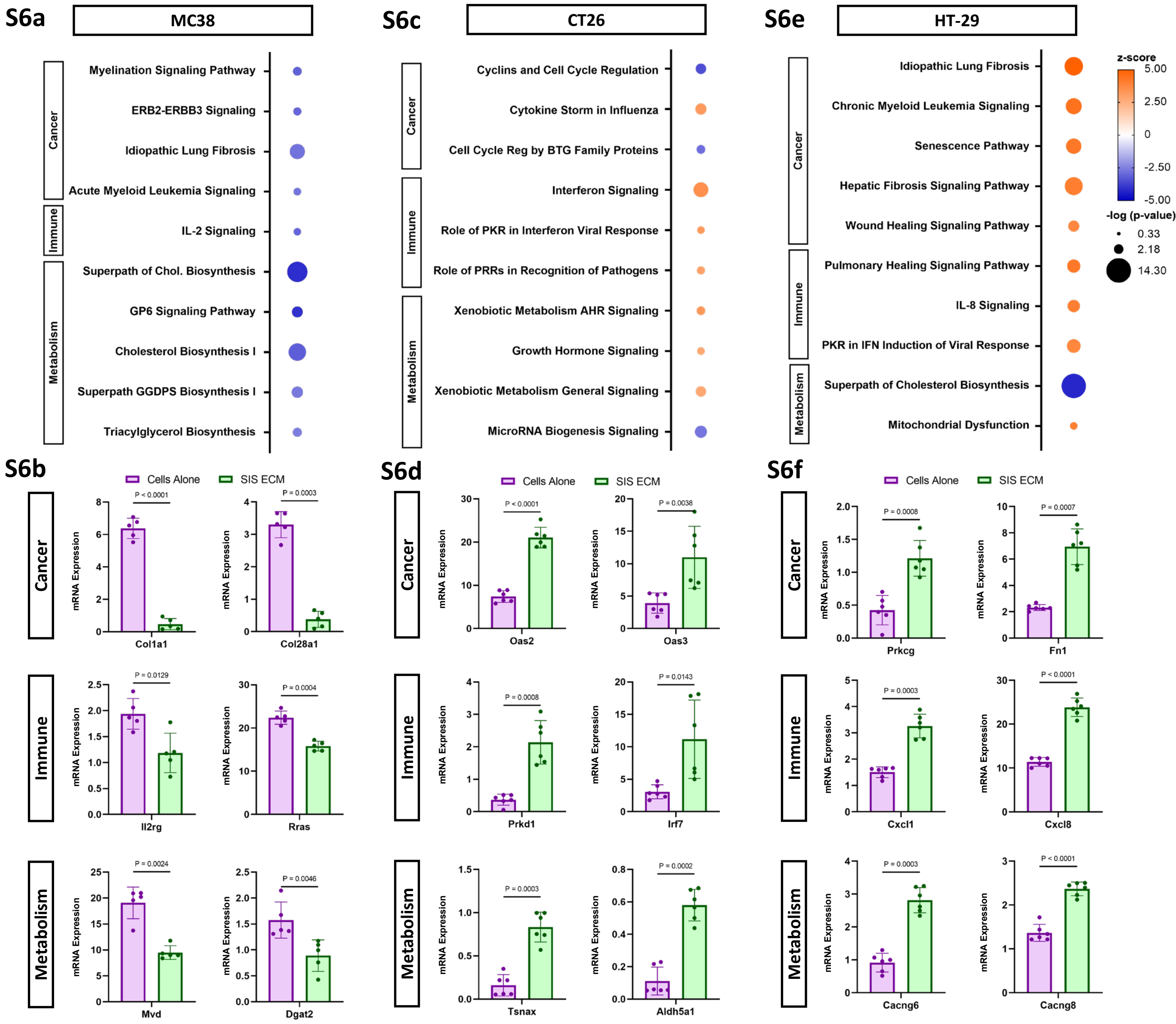

S6a
MC38
Cancer
Immune
Metabolism
Myelination Signaling Pathway
ERB2-ERBB3 Signaling
Idiopathic Lung Fibrosis
Acute Myeloid Leukemia Signaling
IL-2 Signaling
Superpath of Chol. Biosynthesis
GP6 Signaling Pathway
Cholesterol Biosynthesis I
Superpath GGDPS Biosynthesis I
Triacylglycerol Biosynthesis
S6c
CT26
Cyclins and Cell Cycle Regulation
Cytokine Storm in Influenza
Cell Cycle Reg by BTG Family Proteins
Interferon Signaling
Role of PKR in Interferon Viral Response
Role of PRRs in Recognition of Pathogens
Xenobiotic Metabolism AHR Signaling
Growth Hormone Signaling
Xenobiotic Metabolism General Signaling
MicroRNA Biogenesis Signaling
S6e
HT-29
Idiopathic Lung Fibrosis
Chronic Myeloid Leukemia Signaling
Senescence Pathway
Hepatic Fibrosis Signaling Pathway
Wound Healing Signaling Pathway
Pulmonary Healing Signaling Pathway
IL-8 Signaling
PKR in IFN Induction of Viral Response
Superpath of Cholesterol Biosynthesis
Mitochondrial Dysfunction
z-score
5.00
2.50
0
-2.50
-5.00
-log (p-value)
0.33
2.18
14.30
S6b
Cells Alone
SIS ECM
mRNA Expression
P < 0.0001
Col1a1
P = 0.0003
Col28a1
P = 0.0129
Il2rg
P = 0.0004
Rras
P = 0.0024
Mvd
P = 0.0046
Dgat2
S6d
P < 0.0001
Oas2
P = 0.0038
Oas3
P = 0.0008
Prkd1
P = 0.0143
Irf7
P = 0.0003
Tsnax
P = 0.0002
Aldh5a1
S6f
P = 0.0008
Prkcg
P = 0.0007
Fn1
P = 0.0003
Cxcl1
P < 0.0001
Cxcl8
P = 0.0003
Cacng6
P < 0.0001
Cacng8

**S6g**

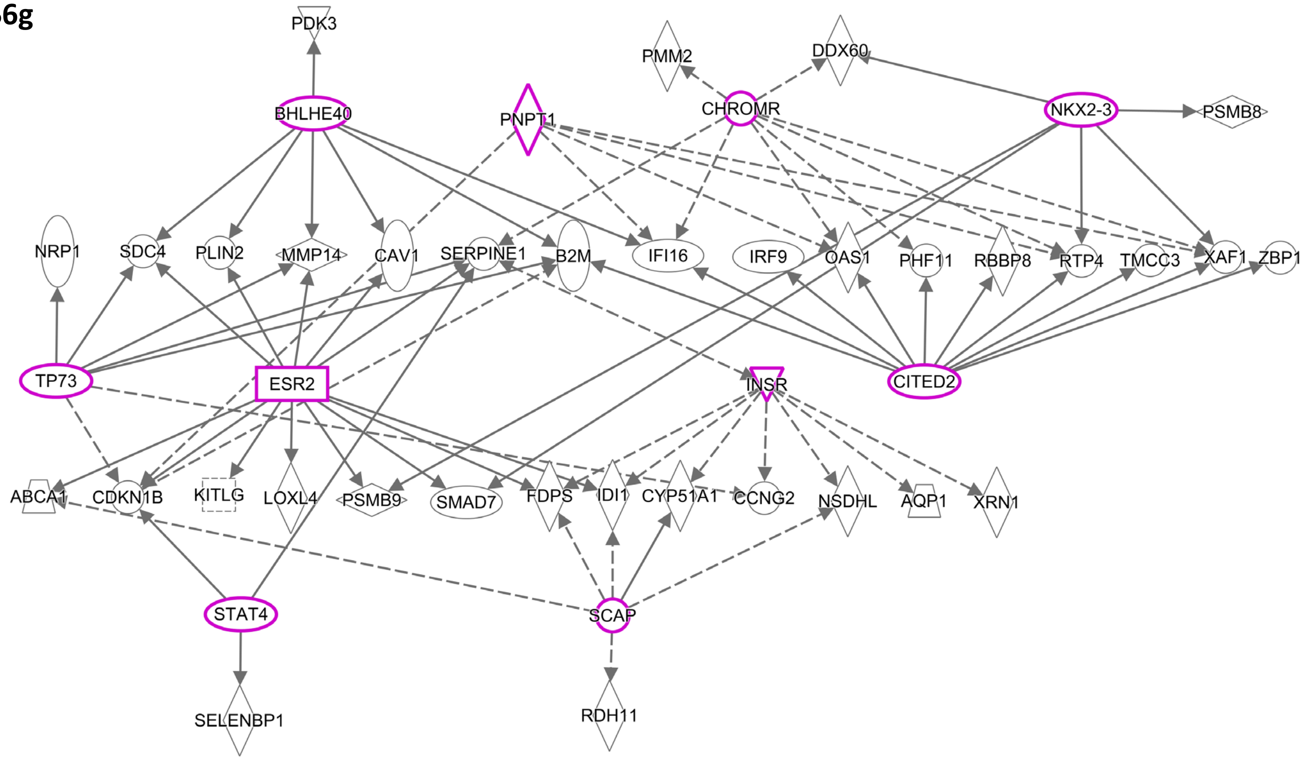
PDK3
PMM2
DDX60
BHLHE40
PNPT1
CHROMR
NKX2-3
PSMB8
NRP1
SDC4
PLIN2
MMP14
CAV1
SERPINE1
B2M
IFI16
IRF9
OAS1
PHF11
RBBP8
RTP4
TMCC3
XAF1
ZBP1
TP73
ESR2
INSR
CITED2
ABCA1
CDKN1B
KITLG
LOXL4
PSMB9
SMAD7
FDPS
IDI1
CYP51A1
CCNG2
NSDHL
AQP1
XRN1
STAT4
SCAP
SELENBP1
RDH11

**S6h**

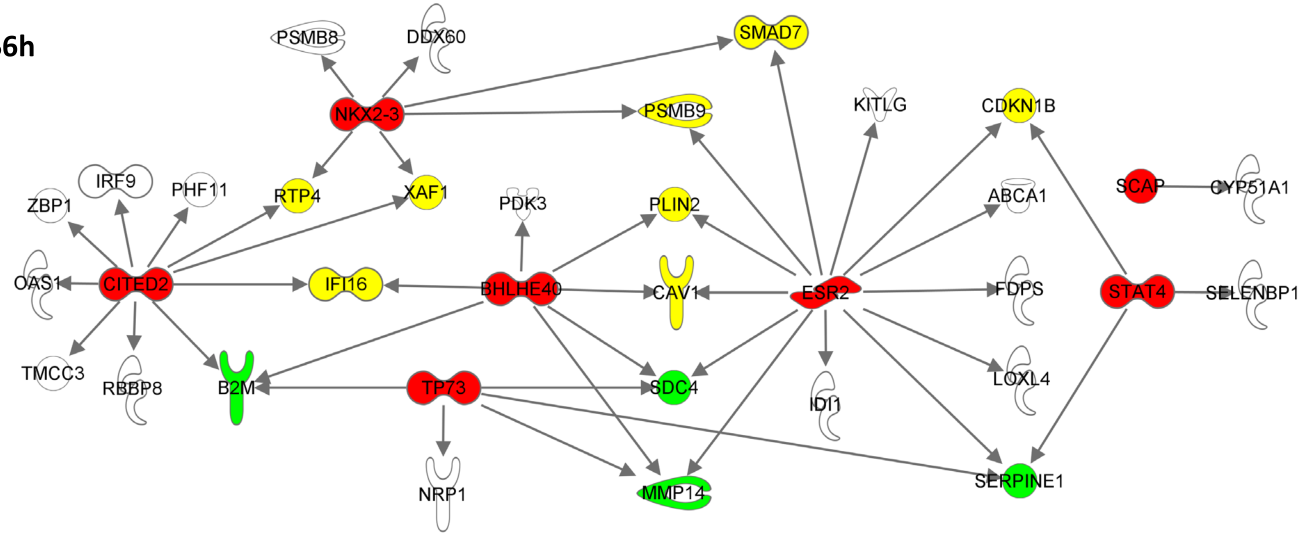
PSMB8
DDX60
SMAD7
NKX2-3
PSMB9
KITLG
CDKN1B
IRF9
ZBP1
PHF11
RTP4
XAF1
PDK3
PLIN2
ABCA1
SCAP
CYP51A1
OAS1
CITED2
IFI16
BHLHE40
CAV1
ESR2
FDPS
STAT4
SELENBP1
TMCC3
RBBP8
B2M
TP73
SDC4
IDI1
LOXL4
NRP1
MMP14
SERPINE1

**S6i**

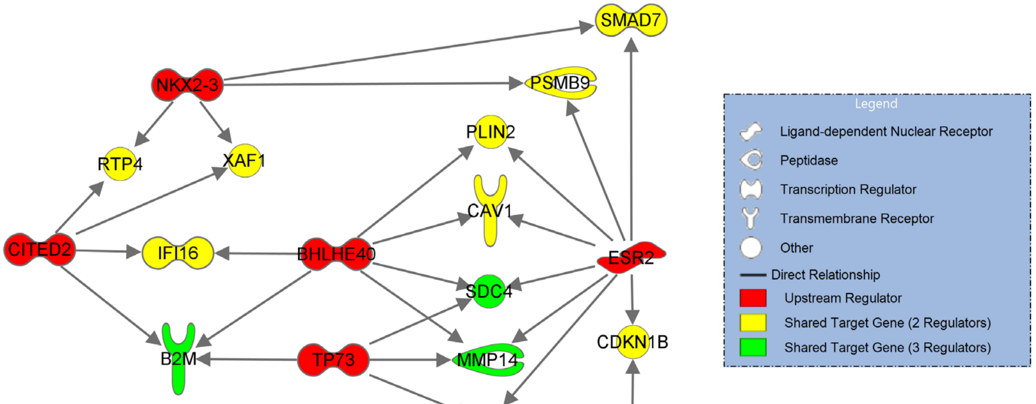
SMAD7
NKX2-3
PSMB9
PLIN2
RTP4
XAF1
CAV1
CITED2
IFI16
BHLHE40
ESR2
SDC4
B2M
TP73
MMP14
CDKN1B
SERPINE1
STAT4
Legend
Ligand-dependent Nuclear Receptor
Peptidase
Transcription Regulator
Transmembrane Receptor
Other
Direct Relationship
Upstream Regulator
Shared Target Gene (2 Regulators)
Shared Target Gene (3 Regulators)

**S6j**

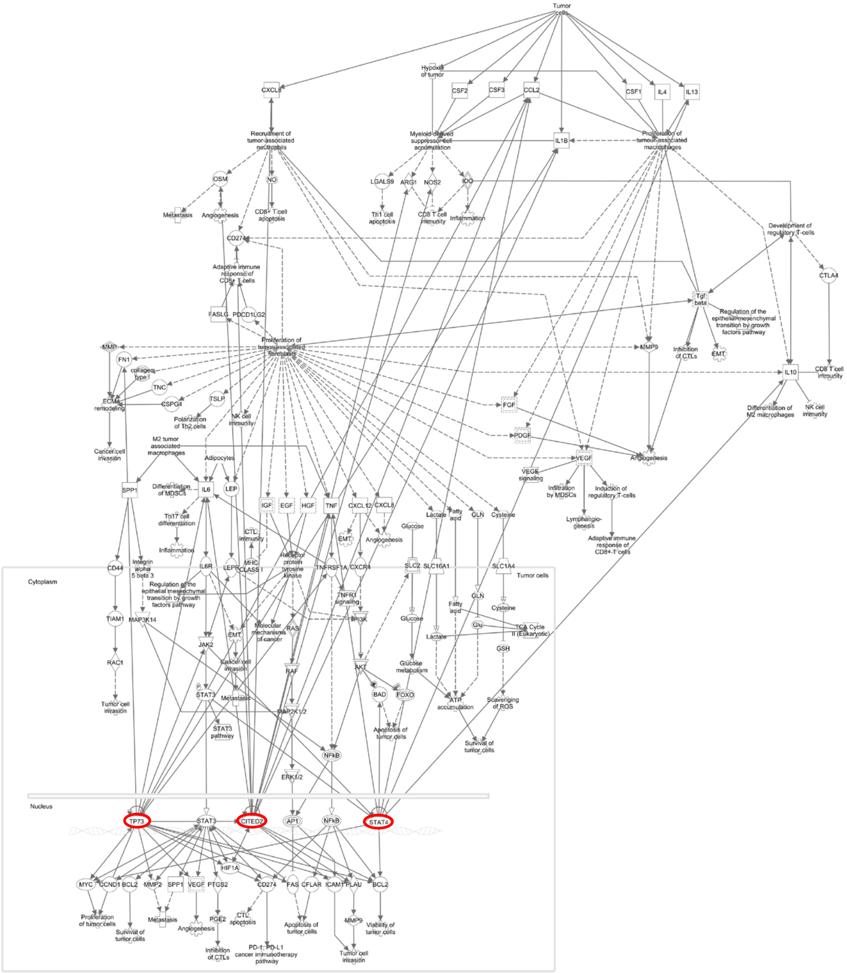

**S6k**

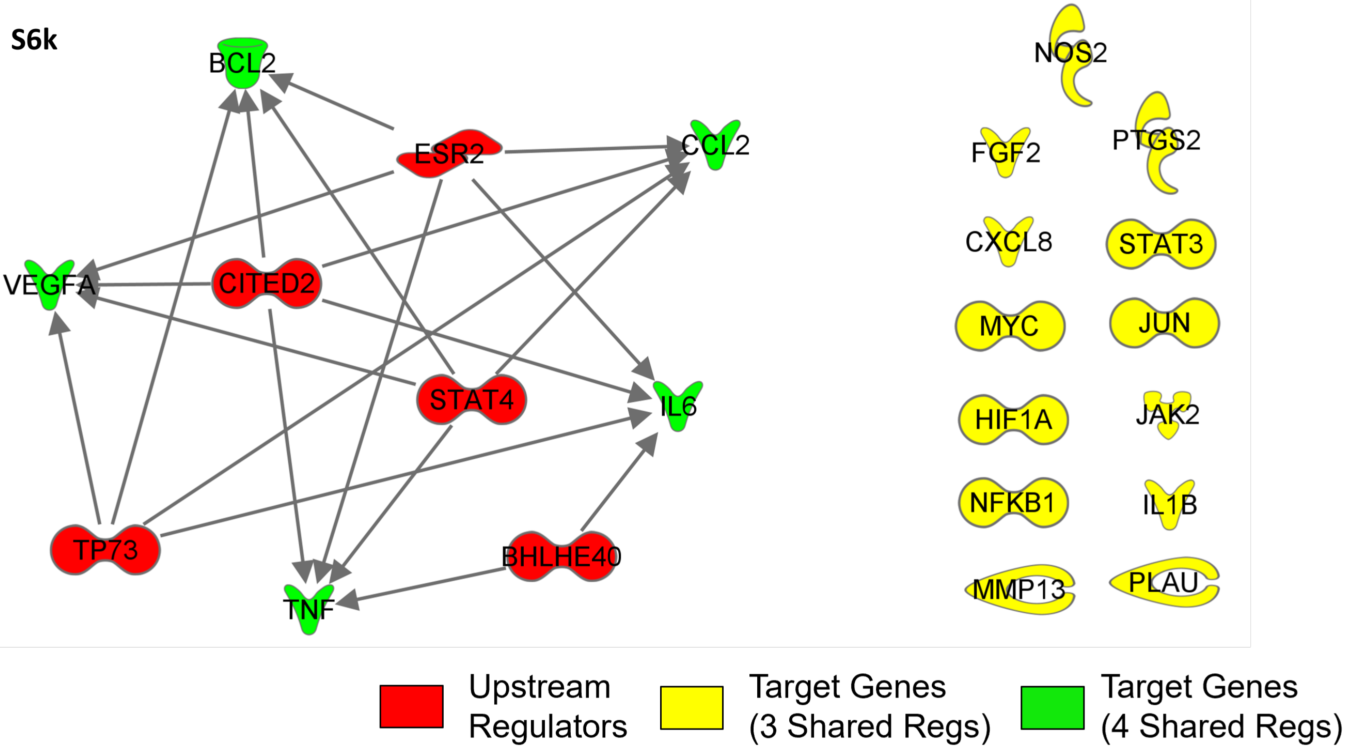
BCL2
ESR2
CCL2
VEGFA
CITED2
STAT4
IL6
TP73
BHLHE40
TNF
NOS2
FGF2
PTGS2
CXCL8
STAT3
MYC
JUN
HIF1A
JAK2
NFKB1
IL1B
MMP13
PLAU
Upstream Regulators
Target Genes (3 Shared Regs)
Target Genes (4 Shared Regs)

**Figure S6**: Bubble plots of the top 10 IPA predicted activation and top 2 genes identified within the assigned categories by cell line a-b. MC38 c-d. CT26 e-f. HT-29 g. All conserved upstream regulators and their downstream targets across CRC cell lines predicted by IPA. h. Only direct interactions between upstream regulators and target genes i. Direct interactions between gene targets with multiple shared regulators j. IPA tumor microenvironment pathway showing an example of interaction between target genes with 3 identified upstream regulators k. Interaction network between the identified upstream regulators and their most common targets within the TME pathways.

S7

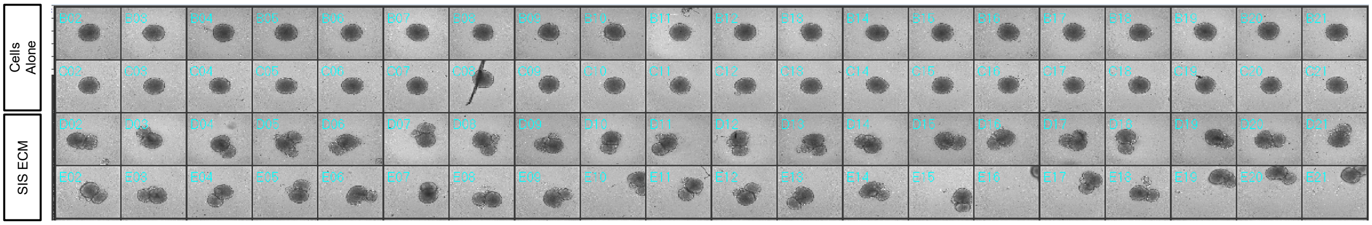

**Figure S7**: Representative images of HT-29 cells alone spheroids and MatriSpheres cultured in 384-well plates for 7 days.

**S8**

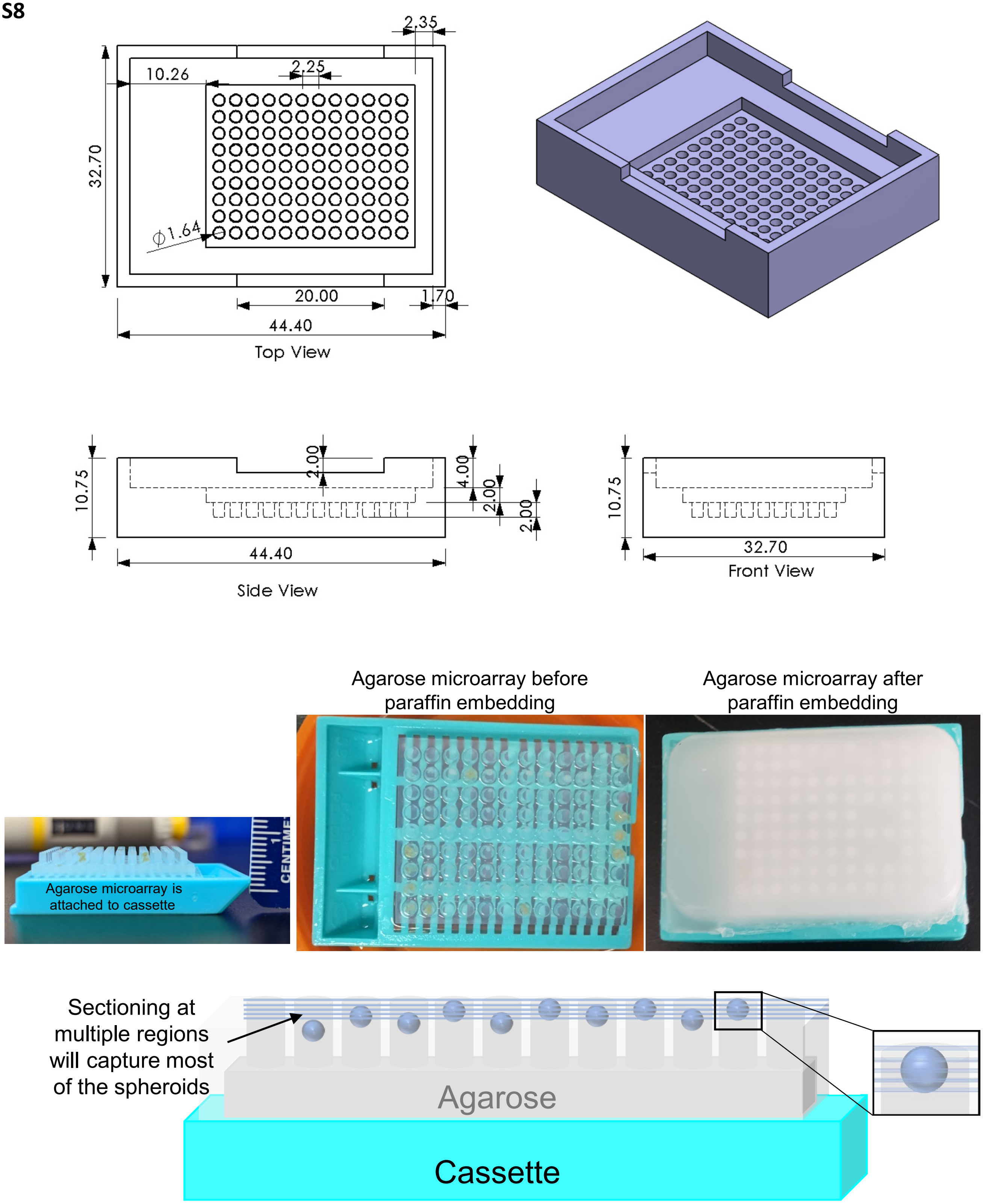
2.35
10.26
2.25
32.70
ϕ1.64
20.00
1.70
44.40
Top View
10.75
2.00
4.00
2.00
2.00
44.40
Side View
10.75
32.70
Front View
Agarose microarray before paraffin embedding
Agarose microarray after paraffin embedding
Agarose microarray is attached to cassette
Sectioning at multiple regions will capture most of the spheroids
Agarose
Cassette

**Figure S8**: Schematic design of the 3D printed spheroid microarray mold, agarose embedding of spheroids and sectioning diagram.

**SVid 1**: MC38 cells alone spheroid formation from initial seeding to day 3 of culture acquired by time-lapse imaging.

**SVid 2**: MC38 + SIS ECM MatriSphere formation from initial seeding to day 3 of culture acquired by time-lapse imaging.

**SVid 3**: MC38 + Type I Collagen MatriSphere formation from initial seeding to day 3 of culture acquired by time-lapse imaging.

**SVid 4**: MC38 + Matrigel MatriSphere formation from initial seeding to day 3 of culture acquired by time-lapse imaging.

**SVid 5**: CT26 cells alone spheroid formation from initial seeding to day 3 of culture acquired by time-lapse imaging.

**SVid 6**: CT26 + SIS ECM MatriSphere formation from initial seeding to day 3 of culture acquired by time-lapse imaging.

**SVid 7**: CT26 + Type I Collagen MatriSphere formation from initial seeding to day 3 of culture acquired by time-lapse imaging.

**SVid 8**: CT26 + Matrigel MatriSphere formation from initial seeding to day 3 of culture acquired by time-lapse imaging.

**SVid 9**: HT-29 cells alone spheroid formation from initial seeding to day 3 of culture acquired by time-lapse imaging.

**SVid 10**: HT-29 + SIS ECM MatriSphere formation from initial seeding to day 3 of culture acquired by time-lapse imaging.

**SVid 11**: HT-29 + Type I Collagen MatriSphere formation from initial seeding to day 3 of culture acquired by time-lapse imaging.

**SVid 12**: HT-29 + Matrigel MatriSphere formation from initial seeding to day 3 of culture acquired by time-lapse imaging.
